# TMEM55B downregulation promotes neurodegeneration in Alzheimer’s disease models

**DOI:** 10.64898/2026.10.21.760809

**Authors:** Bidisha Roy, Viorica Raluca Contu, Maciej Jeziorek, Wilma J. Friedman, Ching-On Wong

## Abstract

Lysosomal distribution is critical for numerous biological functions within the cell, including protein degradation in autophagosomes, cholesterol homeostasis, and cell invasion. Changes in the positioning of lysosomes and dysregulation of lysosomal trafficking are known to contribute to different human pathologies, such as lysosomal storage disease, neurodegeneration, and cancer. Here we report the critical role of the lysosomal **T**ransmembrane **P**rotein **55B** (TMEM55B) in mediating neurodegeneration. Using human clinical-pathological datasets, transgenic flies and iPSC-derived neurons, we demonstrate that loss of function of dTMEM55 potentiates neuronal degeneration. We found that TMEM55B levels are lowered in severe Alzheimer’s Disease (AD) cases, validated from microarray, single neuron and total RNA Seq datasets from brain sections of the ROSMAP cohort. Our results demonstrates that loss of function of dTMEM55 results in accumulation of lysosomes in the fly’s larval neuronal axons, enhances Tau-mediated retinal degeneration, locomotory dysfunction, lethality, and cell death in the fly AD model. The iPSC-derived AD neurons exhibit reduced TMEM55B levels, autophagic flux defects and increased accumulation of phosphorylated Tau in the lysosomes, confirming the observations in the fly. Overall, our findings indicate that TMEM55B downregulation could contribute to neurodegeneration through defects in lysosomal transport, potentially hindering autophagic clearance of Tau. Our findings have significant relevance in validating the role of lysosomal proteins in normal lysosomal function and maintaining healthy neurons. This work supports continued research into lysosome-mediated neurodegeneration and may provide a basis for exploring potential therapeutic approaches for lysosome-associated neuronal disorders.

**Highlights:**

- TMEM55B expression is significantly reduced in postmortem brain tissues of patients with severe AD.
- Genetic loss of TMEM55B enhances Tau-mediated neurodegeneration in *Drosophila*, indicating a conserved neuroprotective function.
- TMEM55B regulates neuronal lysosomal positioning across species, and its loss disrupts lysosomal transport.
- iPSC-derived Alzheimer’s disease neurons exhibit late-stage TMEM55B downregulation concomitant with increased accumulation of pathological phosphorylated Tau.
- Impaired lysosomal trafficking provides a mechanistic link between TMEM55B loss and progressive neurodegeneration.

## Introduction

Alzheimer’s disease (AD) is one of the most common types of dementia. An estimated 5.8 million Americans aged 65 years and older were living with Alzheimer’s dementia in 2020, a number projected to rise to approximately 13.8 million by mid-century (2050–2060) in the absence of effective preventive or disease-modifying interventions (2020). Currently, there is no single effective, disease-modifying therapy available for AD, and therefore, understanding the molecular mechanisms underlying the pathophysiology is important, and continuous positive efforts should be implemented to identify new approaches for the effective prevention and treatment of this disease.

Dysfunctional autophagosomes and lysosomes are recognized as early pathological features in Alzheimer’s disease (AD), preceding the classical deposition of amyloid-β plaques and Tau-containing neurofibrillary tangles (Nixon and Yang 2011). Swelling of endosomal compartments enriched in amyloid precursor protein (APP) peptides (Takahashi et al. 2002), lysosomal deficits, and abnormal accumulation of autophagic vacuoles have been observed in human AD brains and corresponding mouse models (Cataldo et al. 1997), (Nixon and Yang 2011), (Nixon et al. 2005). Genetic studies further support lysosomal involvement, as polymorphisms in several lysosomal cathepsin genes are associated with increased AD susceptibility (Bhojak et al. 2001), (Papassotiropoulos et al. 1999). In addition, deletion of the lysosomal protease inhibitors cystatin B/C has been reported to ameliorate AD-related phenotypes in mice (Mi et al. 2007), (Yang et al. 2011), (Yang et al. 2014). Together, impairment of the autophagosome–lysosomal system and the resulting disruption of intracellular signaling and trafficking have been strongly linked to neurodegeneration (Sorkin and von Zastrow 2009; Dobrowolski and De Robertis 2011), (Taelman et al. 2010), (Killisch et al. 1991), (Lipinski et al. 2010), (Nixon 2013).

Autophagy is essential for the removal of protein aggregates, damaged proteins, and dysfunctional organelles, whose accumulation is implicated in AD and other neurodegenerative disorders (Nixon 2013). Autophagic activity depends on lysosomal function, which is regulated by cellular nutrient status. Lysosomes act as amino-acid sensors through a complex involving vacuolar ATPase, Ragulator, and Rag GTPases that recruit mTORC1 to lysosomal membranes (Laplante and Sabatini 2012). Activation of mTORC1 by Rheb downstream of growth-factor signaling (Inoki et al. 2003), (Tee et al. 2003), controls lysosomal and autophagosome biogenesis via TFEB and the CLEAR gene network (Sardiello et al. 2009), (Settembre et al. 2012).

The transcription factors TFEB and TFE3 act as master regulators of lysosomal biogenesis (Sardiello et al. 2009), (Martina et al. 2014). Lysosomal positioning, modulated by cellular stimuli, regulates autophagic flux, and starvation-induced perinuclear lysosome–autophagosome fusion is critical for cargo degradation (Kimura et al. 2008), (Korolchuk et al. 2011).

TMEM55B is a lysosomal protein implicated in regulating lysosomal positioning and phosphatidylinositol 5-monophosphate abundance in the nucleus (Ungewickell et al. 2005), (Zou et al. 2007). It recruits the dynein adaptor JIP4 to lysosomes, promoting retrograde transport, while depletion of TMEM55B or JIP4 causes peripheral lysosome accumulation (Willett et al. 2017). TMEM55B is a TFEB/TFE3 target, with expression induced by starvation or lysosomal stress, suggesting transcriptional regulation of lysosomal positioning. Previous studies have suggested TMEM55B as a potential target of TFEB and TFE3 (Sardiello et al. 2009{Pastore, 2016 #194; Martina et al. 2016; Pastore et al. 2016). To investigate this, Puertollano and coworkers, transfected HeLa cells with adenoviruses expressing constitutively active TFEB or TFE3. Activation of TFEB and TFE3 resulted in a significant increase in TMEM55B mRNA and protein levels, supporting a role for these transcription factors in modulating TMEM55B expression (Willett et al. 2017). Post-translational modifications, including S-palmitoylation and Erk/MAPK-mediated phosphorylation, further modulate TMEM55B function (Takemasu et al. 2019). TMEM55B also regulates autophagosome–lysosome fusion, lysosomal repair, and TFE3 nuclear translocation under stress (Jeong et al. 2024), interacts with amino-acid sensing complexes (Hashimoto et al. 2018), and modulates cholesterol metabolism via PI(4,5)P2 phosphatase activity (Qin et al. 2020).

Dobrowlowski et al. (Nnah et al. 2019) identified TFEB-regulated endocytic genes, including MCOLN1, which mediates lysosomal Ca²⁺ release to activate calcineurin, promoting TFEB nuclear translocation, autophagy, and lysosomal biogenesis (Medina and Ballabio 2015), (Medina et al. 2015). In contrast, the role of TMEM55B in lysosomal positioning and neurodegeneration remains largely unexplored. While several studies have investigated the contribution of MCOLN1 to neurodegenerative processes (Gao et al. 2024), the mechanistic and functional involvement of TMEM55B in this context is still unknown. Therefore, we focused on TMEM55B as a novel and unexplored candidate in lysosome-mediated neurodegeneration.

Even though few groups have investigated putative roles of this molecule in autophagy, a clear mechanistic underpinning in its role in regulating Tau mediated neurodegeneration using in vitro, in vivo, patient derived clinical datasets and iPSC derived differentiated neurons have not been previously shown. In this study, we report the critical role of the lysosomal transmembrane protein TMEM55B in mediating neurodegeneration in transgenic *in vivo* fly and *in vitro* iPSC cells derived from Alzheimer’s disease patients. TMEM55 levels are lowered in severe AD cases, validated from single neuronal cell RNA Seq, bulk or total RNA Seq and microarray dataset from dorsolateral pre-frontal lobe brain sections of the ROSMAP cohort.

In this study, we observed that the loss of function of dTMEM55 [*Drosophila TMEM55*] leads to locomotion defects, enhances lethality, accumulation of lysosomes in the axonal nerves, enhances Tau-mediated neurodegeneration. Biochemical analysis revealed that downregulation of dTMEM55 leads to elevated levels of Tau in fly brains. Additionally, we observed that TMEM55B is lowered in iPSC-derived AD neurons in an age-dependent manner. In these iPSC-derived AD neurons we also confirmed, autophagic flux defects, elevation of lysosomal phosphorylated-Tau and accumulation of axonal lysosomes. Our results showed that the knockdown of TMEM55B led to lysosomal positioning defects in HEK293T cells and iPSC-derived AD neurons. Additionally, TMEM55-mediated neurodegeneration phenotypes were enhanced in loss of function MCOLN1 fly models, confirming that both TMEM55 and MCOLN1 proteins additively aid in modulating the neurodegenerative phenotypes.

Overall, our data suggests that the downregulation of TMEM55 can potentially cause neurodegeneration in AD cases, due to lysosome transport defects, preventing the autophagic clearance of Tau. Inefficient clearance of Tau due to autophagic flux defects leads to its accumulation, leading to neuronal degeneration.

## Materials and Methods

### Clinical Dataset Evaluation

We have analysed two independent AD patient cohorts in our studies. The Religious Orders Study (ROS) is a longitudinal clinical-pathologic cohort study of aging and AD (AD). In this study, the AD status was determined by a computer algorithm based on cognitive test performance with a series of discrete clinical judgments made in series by a neuropsychologist and a clinician. In addition, a battery of 21 cognitive performance tests was administered each year to assess cognitive abilities and determine these diagnostic classifications. Upon death, a post-mortem neuropathological evaluation was performed that includes a uniform structured assessment of AD pathology. The procedures follow those outlined by the pathologic dataset recommended by the National AD Coordinating Centre and pathologic diagnoses of AD use NIA-Reagan and modified CERAD criteria, and the staging of neurofibrillary pathology uses Braak Staging. The ROS study is described in detail by Bennett and co-workers (Bennett et al. 2012a). The Memory and Aging Project (MAP) is a longitudinal, epidemiologic clinical-pathologic cohort study of common chronic conditions of aging with an emphasis on decline in cognitive function and risk of AD. This study was designed to complement the ROS study by enrolling individuals with a wider range of life experiences and socioeconomic status into a study of similar structure and design as ROS. Diagnoses of dementia and AD were performed in an identical manner to the ROS study (described above). The MAP study is described in detail by Bennett and coworkers (Bennett et al. 2012a; Bennett et al. 2012b). We have used the ROSMAP clinical dataset, the microarray dataset and the gene expression array dataset for our studies. The Aging, Dementia and Traumatic Brain Injury (TBI) Project is a detailed neuropathological, molecular and transcriptomic characterization of brains of control and probable / possible AD cases from a unique aged population based cohort from the Adult Changes in Thought (ACT) study. Dementia is diagnosed at consensus conferences using DSM-IV criteria (Widiger et al. 1991) and AD using probable or possible AD as defined by NINCDS-ADRDA criteria (McKhann et al. 1984). We have used the clinical-pathological, microarray, RNA Seq and single cell RNA Seq dataset for our analysis.

### ROSMAP differential gene expression analysis

Processed microarray and RNA-Seq datasets were downloaded from the AMP-AD portal Sage Bionetworks and analysed for levels of TMEM55B, MAPT, MCOLN1 and RPL13a mRNAs. [AMP-AD Knowledge Portal: ROSMAP RNA-Seq available at https://www.synapse.org/#!Synapse:syn3388564 and AMP-AD Knowledge Portal: ROSMAP Microarray available at https://www.synapse.org/#!Synapse:syn3800853, (Myers et al. 2007)].

### Single Cell RNA Seq Analysis

Data Analysis was performed using the ROSMAP ‘The Single-cell transcriptomic analysis of Alzheimer’s disease (snRNAseqPFC_BA10) study’ (Mathys et al. 2019). Forty eight brain specimens were selected from the Religious Orders Study and Memory and Aging Project (ROSMAP), based on the following selection criteria: 24 individuals with elevated β-amyloid and other pathological hallmarks of AD, and 24 individuals with no or very low Aβ burden or other pathologies. Data consists of 80,660 droplet-based single-nucleus RNA-seq (snRNA-seq) profiles generated from the prefrontal cortex (Brodmann area 10).

### Fly Husbandry

Flies were maintained at 25°C on a 12-hour light/dark cycle and raised on standard cornmeal media. A complete list and source of stocks used can be found in Table 2.

**Table 1:**
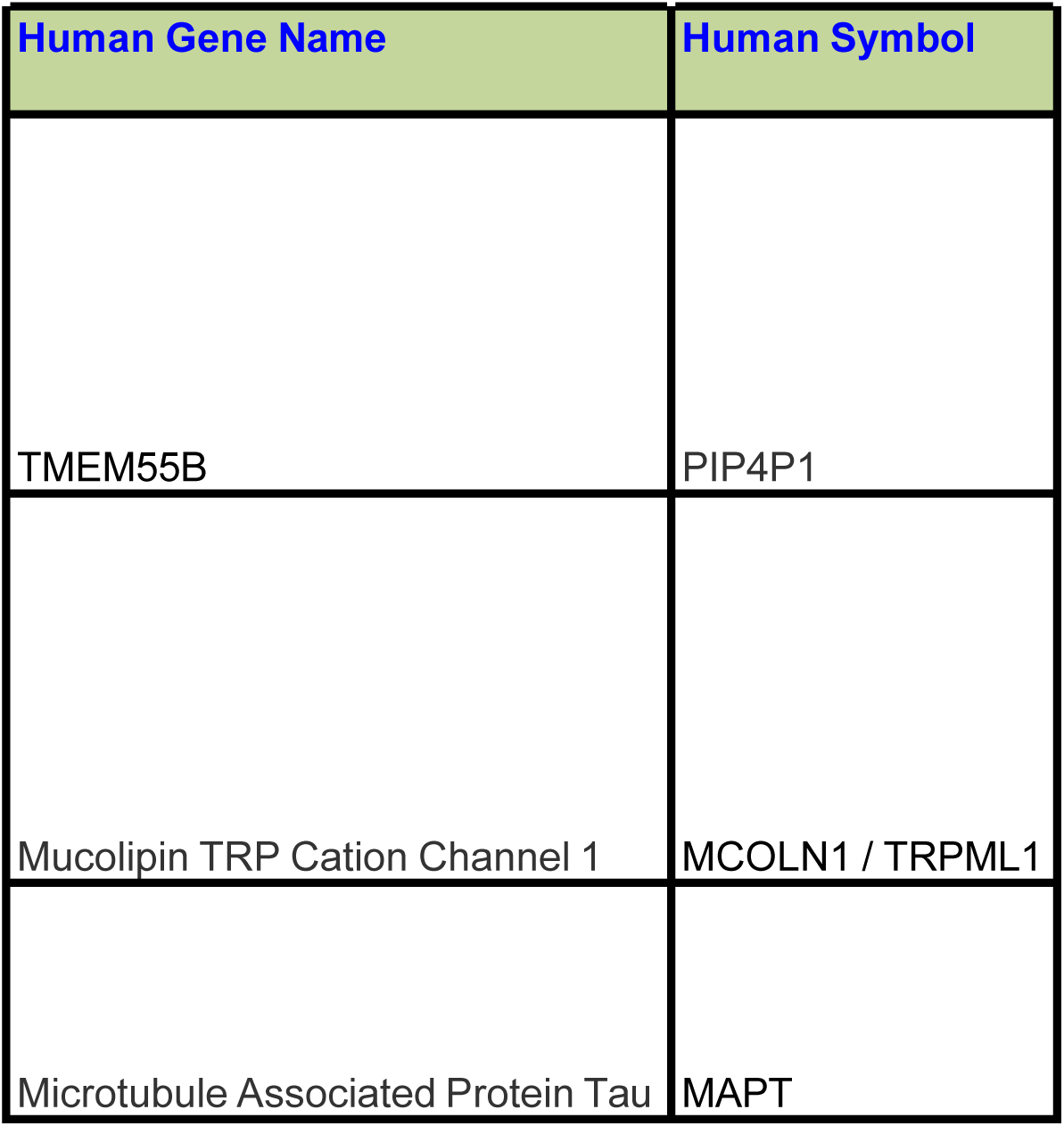
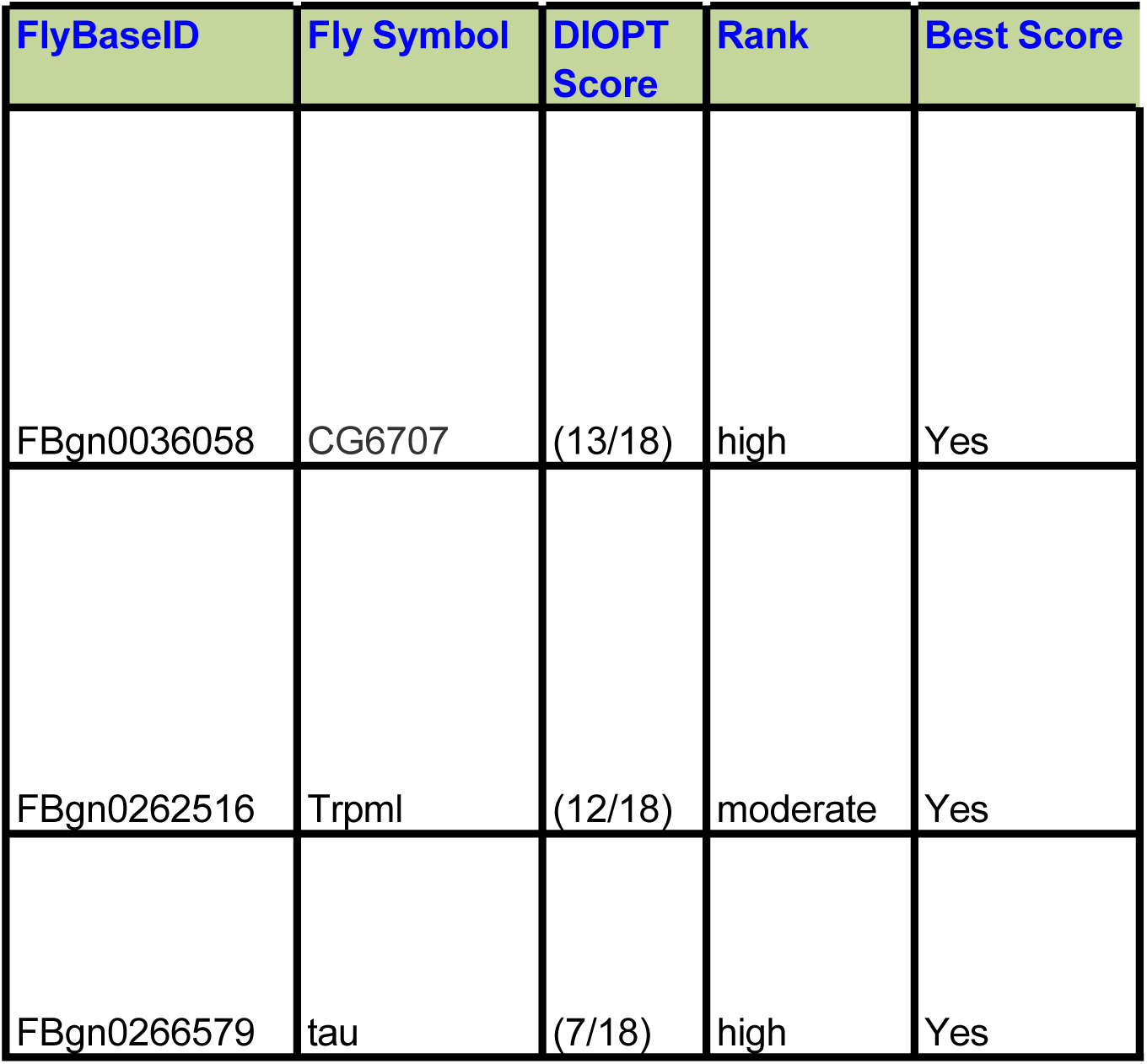
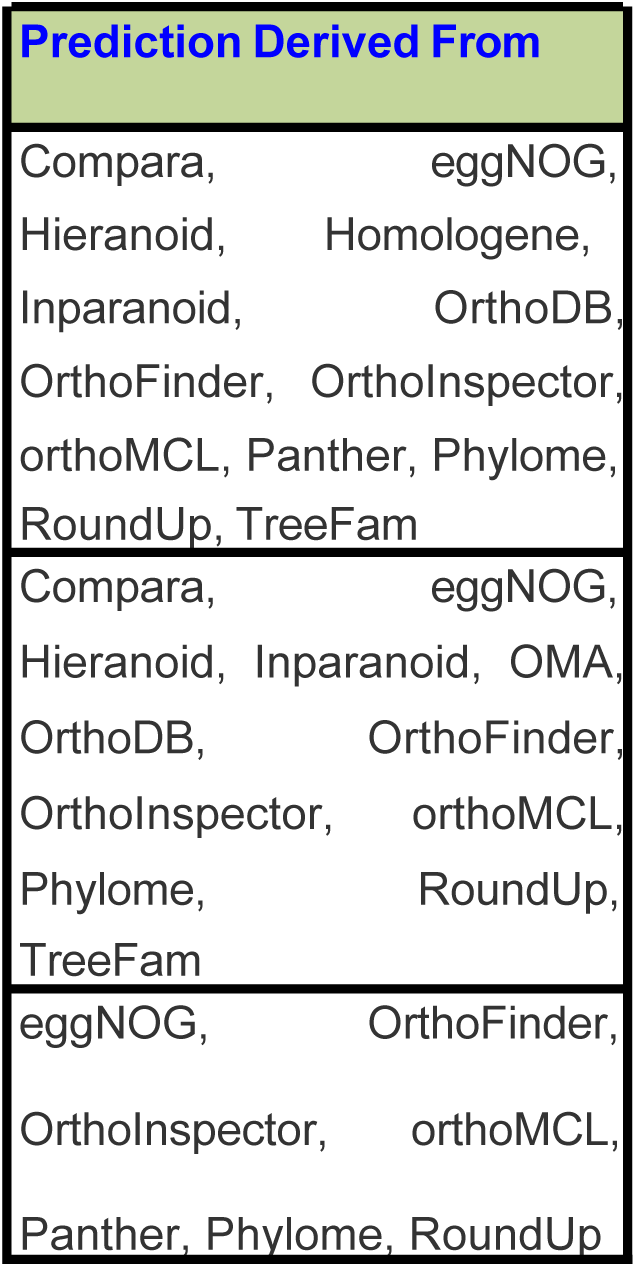
DIOPT-based identification of *Drosophila* orthologs for selected human genes associated with lysosomal and neurodegenerative pathways. This table shows the extent of homology and orthology between specific human genes and their Drosophila melanogaster counterparts using the DIOPT (DRSC Integrative Ortholog Prediction Tool) database. We compared human TMEM55B (PIP4P1), human MCOLN1/TRPML1, and human MAPT to the fly genes CG6707, Trpml, and tau, respectively. The DIOPT scores and ranking categories indicate varying levels of confidence across the group. TMEM55B and MCOLN1 appear strongly to moderately conserved, while MAPT shows a lower but still significant orthologous connection. These findings reinforce the idea that *Drosophila* is a reliable model system for studying the conserved molecular mechanisms that contribute to human disease.

**Table 2:** *Drosophila* fly lines and genetic reagents used in this study. This table lists the various fly stocks used in the functional analysis of genese associated with lysosome mediated neurodegeneration. It includes mutant alleles, RNAi lines, GAL4 drivers, humanized transgenic constructs, and transgenic Tau fly lines. Most stocks were acquired from the Bloomington Drosophila Stock Center, few specialized constructs were provided as gifts from collaborators and few transgenics were constructed by cloning. Stock numbers are provided where available to ensure reproducibility and facilitate future experimental validation.

| Genotype | Source |
| --- | --- |
| CG6707<sup>c01122</sup> /TM6Tb | Bloomington Drosophila Stock Center |
| UAS CG6707 RNAi | Bloomington Drosophila Stock Center |
| OK371 GAL4 | Bloomington Drosophila Stock Center |
| Elav Gal4; gl Tau 1.1 | Bloomington Drosophila Stock Center |
| gl Tau p301L | Bloomington Drosophila Stock Center |
| UAS Tau 1.13 | Bloomington Drosophila Stock Center |
| UAS hLAMP1.HA | Bloomington Drosophila Stock Center |
| C155 Gal4 or Elav Gal4 | Bloomington Drosophila Stock Center |
| UAS dtmem55.mCherry | gift from C.O.Wong |
| Trpml1 / TM6Tb | Bloomington Drosophila Stock Center |
| UAS hTMEM55 1+2 | gift from C.O.Wong |
| UAS hTMEM55 3 | gift from C.O.Wong |
| UAS hTMEM55 4 | gift from C.O.Wong |

nts used in this study.
| Stock Number |
| --- |
| BDSC#85012 |
| BDSC#28316 |
| BDSC#26160 |
| BDSC#51360 |
| BDSC#51377 |
| BDSC#51362 |
| BDSC#86301 |
| BDSC#458 |
| N/A |
| BDSC#28992 |
| N/A |
| N/A |
| N/A |

### Fly strains

Generation of dTMEM55 and human TMEM55B transgenic fly strains: Full-length *dTMEM55* cDNA was PCR-amplified from the cDNA clone IP16043 obtained from the Drosophila Genomics Resource Center, and subcloned into *pmCherry-C1* vector (Clontech). Both human *TMEM55B* cDNA and *mCherry-dTMEM55* cDNA were PCR-amplified and subcloned into *pUAS-attB* vector, respectively. Transgenesis was performed by docking *UAS-hTMEM55B* and *UAS-mCherry-dTMEM55* transgenes to *VK00001* on the 2^nd^ chromosome, respectively.

### Locomotor Assay

Adult males 3-, 10- or 21-days post eclosion (depending on the experiment) were selected for behavioural analysis. A single fly was placed in an empty vial. The fly was gently knocked down to the bottom of tube and allowed 10 s to climb up to the vial top. The distance travelled by individual flies in a 10 s period was measured. All analyses were performed with the experimenter blinded to genotype (Roy and Jackson 2014). For all experiments, climbing assay readouts are presented as both bar graphs and scatter plots showing individual flies. While bar graphs provide a clear summary of central tendency and facilitate comparison of group means across genotypes and ages, they can obscure inter-individual variability inherent to climbing assays and other neurobehavioral measurements. Scatter plots complement summary statistics by displaying individual measurements, allowing visualization of behavioral heterogeneity, data distribution, and potential outliers. Together, these features provide a more transparent and rigorous assessment of age- and genotype-dependent neurobehavioral phenotypes.

### Lethality Assay

8-10 vials were set for every data point on the graphs. 10-12 adult flies were placed in each food vial. They were transferred to freshly prepared vials every 3 days. The number of dead and alive flies were recorded on every transfer. Percentage lethality was calculated as [Total number of dead flies / Total number of flies] x 100 (Bauer et al. 2004; Leung et al. 2024).

### Survival Assay

One-to-three-day-old adult flies were collected in vials (≤20 flies per vial) containing 3% agar dissolved in water. At least four vials of flies were set up for each experimental condition. Fly vials were kept in an incubator set at 25°C and 12-h light-dark cycle. Dead flies in each vial were counted every day until all flies were dead (Leung et al. 2024). TD50 values for each genotype tested are provided in Table 3.

**Table 3.** Median lifespan (TD50) analysis of *Drosophila* genotypes used in aging studies. In this table, the TD50 values, representing time to attain 50% survival for different *Drosophila melanogaster* genotypes, were evaluated using lifespan assays. Data analysis was performed to compare survival across flies across various genetic backgrounds like single heterozygous mutant [CG6707c / + or Trpml 1 / + flies] and double mutant lines, [like CG6707c / Trpml 1 fly lines]. Single heterozygous CG6707c / + and Trpml 1 / + mutants exhibited longer median lifespans, whereas double mutant flies, CG6707c / Trpml 1 showed reduced survival, suggesting that disruption of both these genes additively may negatively impact longevity. The number of flies tested for each genotype is also provided, supporting statistical vigor and robustness of the aging analysis.

| Genotype | TD50 |
| --- | --- |
| CG6707c / + | 58 |
| Trpml 1 /+ | 55 |
| CG6707c / Trpml 1 | 41 |

types used in aging studies.
| No. of flies |
| --- |
| 81 |
| 67 |
| 76 |

### Retinal degeneration

Thoracic-abdominal segments of flies 7-10 days post eclosion were glued onto a microscope slide with transparent nail polish. The head of the fly was positioned such that the right eye was facing the objective lens of the microscope and the left side of the head stuck to the nail polish. In order to accurately quantitate retinal degeneration, we used Leica MZ275 microscope and LAS EZ Leica’s Application Suite Software. The extent of eye area undergoing retinal degeneration (rough pattern with fused ommatidia) was analysed using Image J’s Analyze Area function. Quantitative Bar graphs representing Percent (%) eye roughness = [Area of eye with roughness pattern / Total retinal area] X 100 (Roy and Jackson 2014).

### Immunohistochemistry

Dissection and immunostaining of adult and larval brain and larval fillet preparation were carried out using modifications of published protocols (Langan and Slater 1991). For adult brain, aged flies were anesthetized, decapitated and their heads incubated in fixative (4% paraformaldehyde + PBS + 0.1% Triton X-100) on ice for 30 min. The heads were then dissected to remove the entire brain. For the larval brain, brains were dissected out from third instar wandering larvae in ice cold PBS. For larval NMJ dissection, wandering third instar larvae were pinned on Sylguard plates and subjected to fillet preparation in ice cold PBS. Adult brain, larval brain and larval fillets were then fixed in the above-mentioned fixative for 2 hr on ice. After fixation, brains were washed four times for 15 min each with wash buffer (0.1% PTX = PBS + 0.1% Triton X-100) before incubation in blocking buffer (PBS + 0.1% Triton X-100 + 5% goat serum) for 1 h at room temperature. Tissues were incubated with the primary antibodies overnight at 4°C. This was followed by incubation with fluorescently labelled secondary antibody for 2 h at room temperature. Brains were washed four times for 15 min each in wash buffer and mounted on slides with Vectashield used as an anti-fading agent. The list of primary and secondary antibodies, their source and dilution, used in this study is listed in Table 4.

**Table 4.** Reagents, antibodies, and chemical compounds used for molecular and cellular experiments. This table summarizes the essential experimental reagents utilized in our research study, including primary and secondary antibodies, fluorescent secondary dyes, chemical inhibitors, and molecular biology reagents. Items such as anti-tau antibodies, various lysosomal markers, and compounds for autophagy or endolysosomal pathway analysis are also listed for their company sources and catalog information. The listing of detailed reagent specifications ensures reproducibility in addition to providing a comprehensive reference for the biochemical, imaging, and genetic experiments conducted.

| ***Antibody*** |
| --- |
| *Anti GFP, Rabbit* |
| *Anti GFP, Mouse* |
| *Anti Cleaved Caspase3, Rabbit* |
| *Anti Tau, Mouse* |
| *Anti Tau, Rabbit* |
| *Anti Tau, Mouse* |
| *Anti phospho-Tau (AT8),*<br>*mouse,Phospho-Tau (Ser202,*<br>*Thr205) Monoclonal Antibody* |
| *Anti HA, Mouse* |
| *GAPDH, Mouse* |
| *Anti m-Cherry, Rabbit* |
| *Anti Elav, Rat* |
| *Anti HRP 647* |
| *Alexa anti mouse 488* |
| *Alexa anti Rabbit 488* |
| *Alexa anti mouse 568* |
| *Alexa anti Rabbit 568* |
| *Alexa anti Rat 488* |
| *Anti TMEM55B, Rabbit* |
| *IR Dye 800CW, Goat anti mouse* |
| *IR Dye 680LT, Goat anti rabbit* |
| *Anti LAMP2, Mouse* |
| *Anti Rab 7, Mouse* |
| *Anti MAP2, Chicken* |
| *Beta-actin, Mouse* |
| *OCT-4 antibody, Mouse* |
| *Tuj-1 [anti-Neuronal Class IIIβ-*<br>*Tubulin], Mouse* |
| *LC3 a/b, Rabbit* |

| **Reagents** |
| --- |
| ***SuperScript™ IV First-Strand***<br>***Synthesis System*** |
| ***QuantiTect Reverse Transcription Kit*** |
| ***Luna Universal qPCR Master Mix*** |
| ***DeadEnd™ Fluorometric TUNEL System*** |
| ***Vectashield antifade mounting media with DAPI*** |
| ***Membrane Blocking Solution*** |
| ***Basic chemicals for Westernblotting, Immunohistochemistry and RT PCR*** |
| ***Western Blot related TrisGlycineSDS, TrisGlycine buffer, Nitrocellulose membranes*** |
| ***TRIzol*** |
| ***Chloroquine*** |
| ***Lipofectamine RNAiMAX*** |
| ***siControl or siTMEM55B*** |
| ***PlasmoTest™ - Mycoplasma Detection Kit*** |

Reagents, antibodies, and chemical compounds used for molecular and cellular experiments.
| **Source** | **Application, Dilution** |
| --- | --- |
| *Thermofisher Scientific, Cat No. A11122* | *IHC: 1:500* |
| *DSHB, Cat 12A6-s* | *IHC : 1: 100* |
| *Cell Signaling Technology, Cat No. D175, 5A1E* | *IHC : 1:100; WB : 1:1000* |
| *Thermofisher Scientific, Cat No. 13-6400* | *IHC : 1:100; WB : 1:1000* |
| *Abcam, Cat No. ab76128* | *IHC : 1:50 ; WB : 1:1000* |
| *DSHB, Cat No. 5A6* | *WB : 1:1000* |
| *Thermofisher Scientific, Cat No. MN 1020* | *IHC : 1: 200; WB : 1:1000* |
| *Sigma, Cat No. H9658* | *IHC : 1: 200* |
| *Invitrogen, Cat No. MA5-15738-HRP* | *WB : 1: 2000* |
| *Cell Signaling Technology, Cat No. 43590* | *IHC : 1: 200* |
| *DSHB, Cat No. 7E8A10* | *IHC : 1:20* |
| *Thermofisher Scientific, Cat No. A17331* | *IHC : 1:200* |
| *Thermofisher Scientific, Cat No. A32723* | *IHC : 1:200* |
| *Thermofisher Scientific, Cat No. A32731* | *IHC : 1:200* |
| *Thermofisher Scientific, Cat No. A-11004* | *IHC : 1:200* |
| *Thermofisher Scientific, Cat No. A-110011* | *IHC : 1:200* |
| *Thermofisher Scientific, Cat No. A-A48262* | *IHC : 1:200* |
| *ProteinTech, Cat No. 23992-1-AP* | *IHC : 1: 200; WB : 1:1000* |
| *Licor, Cat No. 926-32210* | *WB : 1: 10,000* |
| *Licor, Cat No. 926-68021* | *WB : 1: 20,000* |
| *DSHB, Cat No. H4B4* | *IHC : 1:20* |
| *Cell Science Technology, Cat No. 95746* | *IHC : 1:100* |
| *Abcam, Cat No. ab5392* | *IHC : 1:500* |
| *DSHB, Cat No. JLA20* | *WB : 1:1000* |
| *Santa Cruz Biotechnologt, Cat No. Sc - 5729* | *WB : 1:500* |
| *Covance, Cat No. MMS-435P* | *WB : 1:1000* |
| *Cell Signaling Technology, Cat No. 4108* | *WB : 1 :1000* |

| **Source** | **Application** |
| --- | --- |
| *Invitrogen, 18091050* | *RT PCR* |
| **Qiagen, Cat No. 205313** | **RT PCR** |
| **New England Biolabs, M3003L** | **RT PCR** |
| **Promega, Cat No. G3250** | **Tunel Assay** |
| **Vector Laboratories, H-1200** |  |
| **Life technologies, Cat No. 000105** | **Western Blot** |
| **Fisher Scientific** | **Western Blot,**<br>**Immunohistochemistry, RT**<br>**PCR** |
| **Fisher Scientific** | **Western Blot** |
| **Invitrogen, 15596018** | **RT PCR** |
| **Sigma Aldrich** | **Autophagic Flux Assay** |
| **Life Technologies, 13778075** | **Cell Transfection** |
| **Ambion Silencer** | **Cell transfection and gene**<br>**knockdown** |
| **Invivogen, rep-pt1** | **Cell-based assay for the**<br>**detection of mycoplasma**<br>**in cell cultures** |

### Western Blotting

Freshly eclosed adult flies were collected and heads were dissected and homogenized in a lysis buffer consisting of 10 mM Tris–Cl (pH 7.4) + NaCl (0.8 M) + EGTA (1 mM, pH 8.0) + 10% sucrose + protease inhibitor cocktail and phosphatase inhibitor cocktail (Roche Applied Science, Indianapolis, IN, USA)(Roy and Jackson 2014). Extracts were mixed with 6X Laemmli SDS Sample Buffer with 9% v/v beta mercaptoethanol and separated by 15% SDS–PAGE. Proteins were transferred to nitrocellulose membranes, blocked in membrane blocking solution, immunoblotted using the desired primary antibodies. The membranes were incubated with the primary antibody overnight, washed in 1 X PBS + 0.1% Tween, incubated in Licor secondary antibodies. The immunoblots were scanned using the Biorad ChemiDoc^TM^ MP Imaging System. Densities of bands were measured using NIH ImageJ (Roy and Jackson 2014). The list of primary and secondary antibodies, their source and dilution, used in this study is listed in Table 4.

### qRT-PCR

Real time qRT-PCR (**Q**uantitative **R**everse **T**ranscription **P**olymerase **C**hain **R**eaction) was performed as previously described (Aizawa et al. 2016; Takahashi et al. 2017; Contu et al. 2023). Total RNA was isolated from 10 to 20 heads of 13 days old flies using the TRIzol Reagent (Invitrogen, 15596018) according with the manufacturer’s instructions. cDNA was then synthesized using SuperScript^TM^ Ⅳ First-Strand Synthesis System (Invitrogen, 18091050) or QuantiTect Reverse Transcription Kit (Qiagen, 205313) and then employed as a template for qPCR on a CFX384 Touch Real-Time PCR Detection System (Bio-Rad). Luna Universal qPCR Master Mix (New England Biolabs, M3003L) was used for performing the quantitative PCR. The levels of each target RNA were normalized to those of *rp49*. The following primers were used for the detection of respective RNAs: *TMEM55B* forward: 5′-GTTCGGACACGTTTTTGTTCAA −3′, reverse: 5′-TGCCGATAGTTACAGCGATGC - 3′; *rp49* forward: 5′-ATCGGTTACGGATCGAACAA-3′, reverse: 5′-GACAATCTCCTTGCGCTTCT-3′

### Cell death assay in the adult brain

For apoptotic cell death visualization, primary antibody anti-cleaved caspase-3 was incubated overnight in PBT containing 5% normal goat serum (Roy and Jackson 2014), (Bawa et al. 2020). Brains were then washed and incubated with Alexa 568 secondary antibody (Invitrogen), washed again and mounted with Vectashield mounting media containing DAPI (Vector Laboratories). Slides were scanned using the LSM510 confocal laser-scanning microscope. Number of cleaved caspase-3 positive cells was counted using the Image J software on 6 micron stacked confocal images. Cell death was also visualized in 21 days or 28 days adult brains of flies, using terminal deoxynucleotidyl transferase biotin dUTP nick end labeling (TUNEL) labeling according to manufacturer’s 20 instructions (DeadEnd**^TM^** Fluorometric TUNEL System, G3250 Promega) [(Bawa et al. 2020), (Dias-Santagata et al. 2007), (Vasudevan and Ryoo 2016)]. Number of TUNEL-positive cells throughout the brain optic lobe was quantified with Image J software.

### Quantification of LAMP1 positive lysosomes in the neuronal axons

For quantification of human LAMP1 positive lysosomes, Z stack images were processed using the ImageJ background correction, color depth reduction to 8 bits and adjustment of the detection threshold. Manual counting using Image J’s multipoint tool aided in quantifying the total number of LAMP1 positive organelles in the third instar wandering larval nerves. These nerves are known to innervate the ventral nerve chord of the larvae and are comprising of axons of motor neurons. The nerve area was measured, by selecting first the area of quantitation using Image J’s free hand tool and subsequently, Analyze, Measure tool. Data represented on the y-axis as Total number of LAMP1 positive lysosomes / Total area of the larval ventral chord nerves.

### Human induced pluripotent stem cell (hiPSC) culture and generation of neurons from human iPSCs

#### A. Differentiation of Human iPSCs into induced neurons

Human induced pluripotent stem cells (hiPSCs) were cultured and differentiated into neurons following a previously established protocol (Hurley et al. 2023). All cell lines were maintained at 37 °C in a humidified atmosphere containing 5% CO₂ and were routinely tested for mycoplasma contamination using the MycoAlert™ PLUS Mycoplasma Detection Kit (PlasmoTest™ - Mycoplasma Detection Kit, invivogen Cat.code: rep-pt1) according to the manufacturer’s instructions. Cells were passaged upon reaching 70–90% confluence, and only cultures maintained for ≤2 months were used for experiments (Contu et al. 2017). Female control and CRISPR/Cas9-edited hiPSC lines harboring a heterozygous presenilin-1 (PSEN1) M146L mutation (Hurley et al. 2023),(Reddy et al. 2016) were maintained in StemFlex™ Medium (Thermo Fisher Scientific; catalog no. A3349401) on six-well plates coated overnight with 50 µg/mL Matrigel (Corning® Matrigel® Matrix) diluted in DMEM/F-12 (Gibco^TM^, Thermofisher Scientific). Cells were routinely passaged using Versene (Thermo Fisher Scientific; catalog no. 15040066), and colonies were regularly examined for characteristic morphology to confirm maintenance of an undifferentiated state. For neuronal differentiation, hiPSCs were dissociated into single cells using Versene followed by Accutase (Sigma-Aldrich; catalog no. A6964) and plated onto coverslips at a density of 60,000 cells per well in 24-well plates or 120,000 cells per well in 12-well plates. Cells were seeded in StemFlex medium supplemented with 10 µM Y-27632 (Sigma-Aldrich; catalog no. Y0503) and 5 µM polybrene (Sigma-Aldrich; catalog no. TR-1003). The following day (day 1), cells were transduced with lentiviral vectors expressing neuronal transcription factors, including NGN2, in StemFlex medium. On day 2, cultures were transitioned to neuronal induction medium (NIM; DMEM/F-12 supplemented with 1% GlutaMAX, 1% non-essential amino acids, and 1% N-2), and NGN2 expression was induced by the addition of 2 µg/mL doxycycline. On day 3, the medium was replaced with fresh NIM containing 2 µg/mL doxycycline, and 1 µg/mL puromycin was added to select for transduced cells. On day 4, cells were washed twice with DMEM/F-12 and switched to neuronal differentiation medium (NDM; Neurobasal medium supplemented with 2% B-27, 1% GlutaMAX, 1% CultureOne, and 200 µM ascorbic acid) containing 2 µg/mL doxycycline and 1 µg/mL puromycin. On day 5, cultures were washed with DMEM/F-12 and maintained in NDM supplemented with 2 µg/mL doxycycline. When residual undifferentiated or proliferating cells were observed, 2 µM cytosine β-D-arabinofuranoside (Ara-C; Millipore Sigma) was added and maintained until proliferating cells were eliminated. Beginning on day 7, half-medium changes were performed every 2–3 days using neuronal maturation medium (NMM; Neurobasal Plus supplemented with 2% B-27 Plus, 1% GlutaMAX, 1% CultureOne, and 200 µM ascorbic acid) containing 2 µg/mL doxycycline. Neuronal culture purity was assessed by immunocytochemistry and Western blot analysis using established neuronal and brain-specific markers. Differentiated neurons were aged for 20 or 40 days in vitro and subsequently used for Western blotting and immunocytochemistry experiments [Supplementary Figure 7A].

#### B. Coating

Glass coverslips (Nunc™ Thermanox™ Coverslips; Thermo Fisher Scientific) and culture plates were coated with poly-D-lysine (10 µg/mL; Thermo Fisher Scientific) prepared in PBS and incubated for 30 min at 37 °C. Following incubation, surfaces were washed extensively with PBS and distilled water, air-dried in a biosafety cabinet for 2 h, and then incubated overnight with Matrigel (100 µg/mL). This sequential coating strategy was critical for ensuring reproducible cell attachment, survival, and efficient neuronal differentiation across cultures [Supplementary Figure 10A].

### Immunocytochemistry of the human iPSC derived neurons

Neuronal cultures were prefixed by directly adding an equal volume of pre-warmed 4% paraformaldehyde (PFA) in phosphate-buffered saline (PBS; 37 °C) to the culture medium, yielding a final concentration of 2% PFA, and incubated for 10 min at room temperature (RT). Cells were then washed once with PBS and fixed with freshly prepared 4% PFA in PBS for an additional 15 min at RT. Following fixation, coverslips were washed once with PBS and twice with PBS containing 0.1% Tween-20 (PBS-T). Cells were permeabilized and blocked simultaneously for 30 min at RT in blocking buffer containing 0.2% Triton X-100. Primary antibodies were diluted in 1% bovine serum albumin (BSA) in PBS and applied to coverslips for overnight incubation at 4 °C. The following day, coverslips were washed with PBS-T at RT and incubated with appropriate fluorophore-conjugated secondary antibodies diluted in 1% BSA in PBS for 30 min at RT in the dark. After secondary antibody incubation, coverslips were washed four times with PBS-T and once with distilled water, then mounted using ProLong™ Gold Antifade Mountant with DAPI (Invitrogen™, Thermo Fisher Scientific). Mounted coverslips were cured overnight in the dark. Confocal images were acquired using a Zeiss LSM 510 confocal microscope and analyzed using ImageJ software. The list of primary and secondary antibodies, their source and dilution, used in this study is listed in Table 4.

### Protein Extraction and Western Blot Analysis of the human iPSC derived neurons

Human iPSC-derived neurons were lysed in SDS-based lysis buffer containing 3% SDS, 5% glycerol, 10 mM Tris-HCl (pH 7.8), 0.02% bromophenol blue, 2% β-mercaptoethanol, and protease inhibitors. Cell lysates were homogenized using QIAshredder spin columns (Qiagen) according to the manufacturer’s instructions by centrifugation, and the clarified lysates were collected. Equal amounts of protein were resolved by SDS–PAGE and subsequently transferred onto polyvinylidene difluoride (PVDF) membranes (Bio-Rad; catalog no. 1620177). Membranes were blocked for 1 h at room temperature in PBS containing 0.1% Tween-20 (PBS-T) and 1% skim milk, followed by overnight incubation at 4 °C with primary antibodies diluted either in PBS containing 3% bovine serum albumin or PBS-T containing 1% skim milk, as appropriate. Following primary antibody incubation, membranes were incubated with horseradish peroxidase (HRP)-conjugated secondary antibodies diluted 1:10,000 in PBS-T for 1 h at room temperature with gentle agitation. Membranes were washed four times with PBS-T for 5 min each and developed using Clarity™ Western ECL Substrate (Bio-Rad; 1:1 mixture). Chemiluminescent signals were detected using a Bio-Rad ChemiDoc™ MP Imaging System, and band intensities were quantified using NIH ImageJ software (Hase et al. 2020). The list of primary and secondary antibodies, their source and dilution, used in this study is listed in Table 4.

### Autophagic Flux assay of the human iPSC derived neurons

Autophagic flux was assessed by monitoring the turnover of microtubule-associated protein 1 light chain 3 (LC3-II) using Western blot analysis on iPSCs, Control and AD neurons grown in the presence or absence of the lysosomal degradation inhibitor chloroquine (Reddy et al. 2016; Mauthe et al. 2018). Chloroquine blocks lysosomal acidification and prevents autophagosome–lysosome degradation, leading to the accumulation of LC3-II when autophagic flux is active. Thus, autophagic flux was inferred from the increase in LC3-II levels upon chloroquine treatment relative to untreated conditions. The magnitude of autophagic flux was quantified as the difference in LC3-II abundance between chloroquine-treated and untreated samples. An increase in LC3-II levels following chloroquine treatment was interpreted as evidence of intact or enhanced autophagic flux, whereas a lack of significant LC3-II accumulation in the presence of chloroquine indicated impaired or reduced autophagic flux. Immunoblots were imaged using a Bio-Rad ChemiDoc™ MP Imaging System, and band intensities were quantified using NIH ImageJ software.

### HEK293T cell culture and immunocytochemistry

HEK293T cells were maintained in Dulbecco’s Modified Eagle Medium (DMEM) supplemented with 10% FBS, L-Glutamine, and MEM non-essential amino acids at 37°C in a humidified 5% CO2 atmosphere. Cells were passaged using 0.25% trypsin when 80–90% confluent. For siRNA-mediated knockdown of TMEM55B, HEK293T cells were transfected with 100nM predesigned siControl or siTMEM55B (Ambion Silencer) using Lipofectamine RNAiMAX (Life Technologies, 13778075) according to the manufacturer’s protocol and incubated for 72 h. The knockdown efficiency was evaluated by immunostaining. HEK293T cells were fixed with 4% PFA in PBS at room temperature for 12 minutes, permeabilized for 5 minutes with 0.5% Triton100 in PBS, blocked with 3% BSA in PBS and incubated with primary antibodies overnight at 4C (1:100 TMEM55B ProteinTech23992-1-AP, 1:100 Rab7 CST 95746, 1:20 Lamp2 DSHB H4B4). The next day, samples were washed, incubated with secondary antibodies (Jackson ImmunoResearch), and mounted with VECTASHIELD Vibrance mounting medium (H1800). Images were collected using Zeiss microscope LSM510 confocal microscope.

For quantification of TMEM55B levels, in the Control and si-RNA mediated TMEM55B knocked down HEK293T cells, Z stack images were processed using the ImageJ Stacks, Z-Project tool. The intensity of TMEM55B and DAPI in the HEK293T cells were measured using Image J’s free hand tool and subsequently, Analyze, Measure mean intensity tool. The data represented on the y-axis is the ratio of TMEM55B Intensity normalised to DAPI Intensity.

For quantification of lysosomal positioning in the Control and si-RNA mediated TMEM55B knocked down HEK293T cells, Z stack images were processed using the ImageJ Stacks, Z-Project tool. The distance of the Rab7 (late endosomal and lysosomal marker) from the nucleus was measured using Image J’s straight line tool and subsequently, Analyze, Measure length tool. It was then normalised with the diameter of the nucleus (marked by DAPI) of the cell. The data represented on the y-axis is the ratio of Rab7 staining length from the nucleus and the diameter of the nucleus.

### Statistical Analysis and Graphs

Statistical analysis and graphs were plotted using GraphPad Prism 8 software.

### Image generation Software

Schematics shown in Figure 11 and Supplementary Figure 9A were partially created using BioRender (https://biorender.com), with the majority of illustrations manually drawn using Microsoft Paint and PowerPoint.

## Results

### TMEM55B RNA levels are lowered in severe AD patients in the clinical ROSMAP microarray and RNA Seq dataset generated from dorso-prefrontal cortical tissues of adult brain

As a first step, we wanted to evaluate if there are any changes in the TMEM55B RNA levels in the AD brains with respect to the control counterparts. Using clinical dataset of AD patients [ROSMAP patient cohort], the patients were first stratified using clinical-pathological hallmark, as mild, moderate and severe AD patients [Supplementary Fig 1A, B and C]. Severe AD patients were identified by clinical –pathological criteria of very high BRAAK score, low CERAD score, high CDx number and low MMSE score [Supplementary Figure 2E]. Microarray gene expression and total RNA sequencing analysis using the dorso-prefrontal cortical regions of adult human brain revealed that the TMEM55B RNA levels were significantly lowered only in the severe AD patients [Figure 1Ci, Supplementary Figure 2Ci]. However, housekeeping gene levels like RPL13A and GAPDH remain unaffected in both the Control and AD patient samples [Figure 1Cii, Supplementary Figure 2Cii]. We also exhibit that lysosomal associated gene MCOLN1 RNA [Figure 1D] level is lowered in the severe AD patients. Additionally, AD associated gene MAPT RNA level is upregulated in the severe AD patients [Figure 1E]. These results indicate that TMEM55B levels are dysregulated in severe AD patients and not in mild or moderate AD cases.

**Figure 1:**
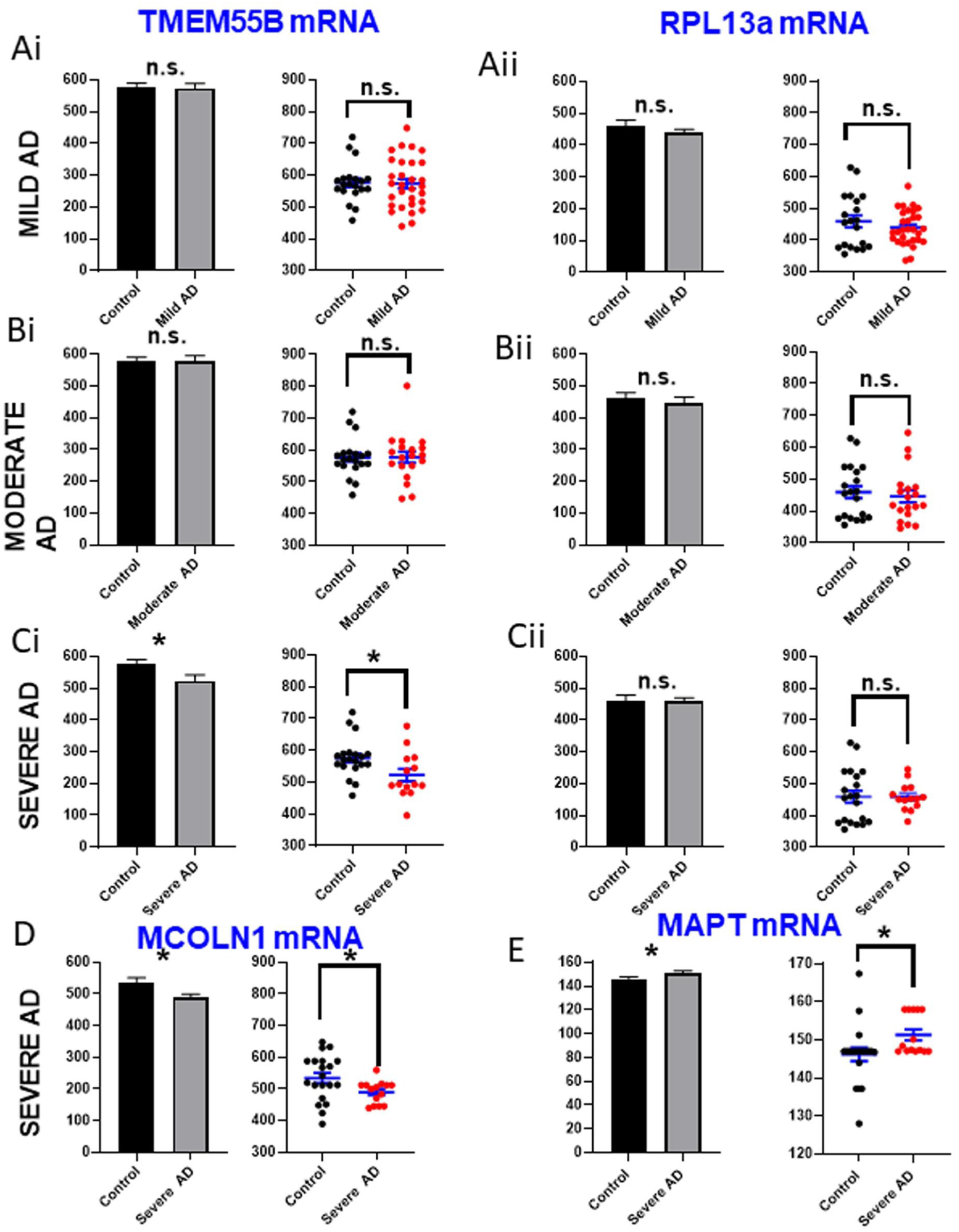
TMEM55B RNA levels are lowered in severe AD patients in the ROSMAP microarray dataset. (Ai-Cii) Bar graphs and scatter plots representing quantitative mRNA levels of TMEM55B and housekeeping gene RPL13a in mild (Ai, Aii) moderate (Bi, Bii), and severe AD (Ci, Cii) cases respectively, using the microarray dataset of ROSMAP studies. Similarly, MCOLN1 and MAPT mRNA levels are compared between control and severe AD patients in (D) and (E) respectively. Control patients: N= 20; Severe AD patients: N=14; Moderate AD patients: 19; Mild AD patients: 30 patients. *: p < 0.05; **: p < 0.01; ***: p < 0.001; n.s: p ≥ 0.05. p-value was calculated using Prism 7’s two-tailed Unpaired t-test.

### TMEM55B RNA levels are lowered in neuronal nucleus of severe AD patients

Since total RNA Seq or microarray ROSMAP dataset used total lysates from the dorso-prefrontal cortex, we wanted to specifically look into the levels of TMEM55B in the neuronal population. Using single-nucleus RNA Seq dataset from the ROS-MAP AD patient consortium, we evaluated TMEM55B levels in the nucleus of neuronal cells present in the dorso-prefrontal cortex of the AD and control patients. Because bulk RNA-seq and microarray analyses from the ROS-MAP dataset were performed on total lysates of the dorsolateral prefrontal cortex, cell type–specific changes in TMEM55B expression cannot be distinguished and may be confounded by contributions from non-neuronal cell types, including glia. To specifically assess neuronal TMEM55B expression, we analyzed single-nucleus RNA-seq data from the ROS-MAP Alzheimer’s disease consortium. This approach enables cell type–resolved interrogation of gene expression, allowing evaluation of TMEM55B levels specifically within neuronal nuclei from the dorsolateral prefrontal cortex of Alzheimer’s disease and control individuals, without contamination from other brain cell populations. We found the levels of TMEM55B levels were significantly lowered in the neuronal nucleus of AD patients in comparison to the control [Supplementary Figure 3]. This indicates that TMEM55B levels are lowered in the neuronal population of AD patients, and hints towards a possible link of this TMEM55B downregulation in neuronal degeneration.

### Loss of function of dTMEM55 leads to locomotion defects in 10 days old flies and enhanced lethality

Since human TMEM55B shares 99% homology with *Drosophila* dTMEM55 (CG6707) [Table 1], we decided to model TMEM55B function and its relationship with neurodegeneration in *Drosophila*. We utilized a transposon insertion allele of *dTMEM55* (CG6707c / +) and a transgenic *dTMEM55*-targeting RNAi line (UAS CG6707 RNAi) to evaluate how TMEM55B deficiency affects neurodegeneration in AD and understand the mechanisms underlying abnormal Tau accumulation [a key neuropathological hallmark of AD]. Additionally, to ascertain that the studies in all the fly TMEM55B models are due to altered TMEM55B levels, we used RT-PCR to probe for TMEM55B RNA levels [Supplementary Figure 8, A-D]. We found that the TMEM55B levels were lowered in loss of function flies [CG6707c / + (Supp Figure 7A), OK371 Gal4 /+; UAS CG6707 RNAi/+ (Supp Figure 8C)] and elevated in the gain of function flies [CG6707 Dup / + (Supp Figure 8A; OK371 Gal4 / UAS *dTMEM55* (Supp Figure 8B)].

Altered locomotor function is a common phenotype in *Drosophila* models of neurodegeneration. The climbing assay is one of the simplest and routinely used method for measuring the motor function of flies. In this assay, flies are placed in an empty vial, and the vial is tapped to drop the flies to the bottom of the vial. One can either calculate the distance climbed by flies in a defined time period or alternatively one can calculate the percentage of flies that climbed to a certain height after a certain time. The assay is based on the negative geotaxis of flies. While healthy, young flies immediately start climbing the vial wall when they fall to the bottom, the rate and speed of climbing is reduced in older flies and flies that express a disease gene in their nervous system (Li et al. 2010). Previous study of Roy et al and others (Roy and Jackson 2014), (Nitta and Sugie 2022), have used locomotory assays as a means to evaluate neurodegeneration to probe Tau and alpha Synuclein interaction (Roy and Jackson 2014). In our current study, we have used similar paradigms like climbing assays to evaluate how downregulation of dTMEM55 affects age dependent locomotory defects in flies. Both *dTMEM55* heterozygous mutant flies [CG6707c / +, Figure 2Aii] and flies with RNAi mediated TMEM55B downregulation using a motor neuron [OK371 Gal4 /+; UAS CG6707 RNAi/+, Figure 2Bii] and pan-neuronal driver [Elav Gal4 / UAS CG6707 RNAi, Figure 2Cii] showed locomotory defects in 10 days old flies. However, the 3-day old flies in these transgenics did not show any defects in these climbing assays [Figure 2Ai, 2Bi and 2Ci]. In a similar fashion, the dTMEM55 knockout flies (CG6707 ^KO^ / +) also displayed climbing defects in older flies (10 days old) and not in the 3 days old younger flies [Supplementary Figure 7B, 7C]. Since behavioural output like locomotory or climbing assays are reliable readouts for neuronal dysfunction, all these results lead to the conclusion that downregulation of TMEM55B results in neuronal dysfunction in an age dependent manner. For all these experiments, climbing assay readouts are represented across individual flies using scatter plots [Supplementary Figure 4, 7C]. Scatter plots were used along with bar graphs to capture inter-individual variability and behavioral heterogeneity inherent to climbing assays and other neurobehavioral measurements. This representation allows direct visualization of data distribution and potential outliers, providing a more transparent and accurate assessment of age-dependent and genotype-specific neurobehavioral phenotypes that may be obscured by summary statistics alone.

**Figure 2:**
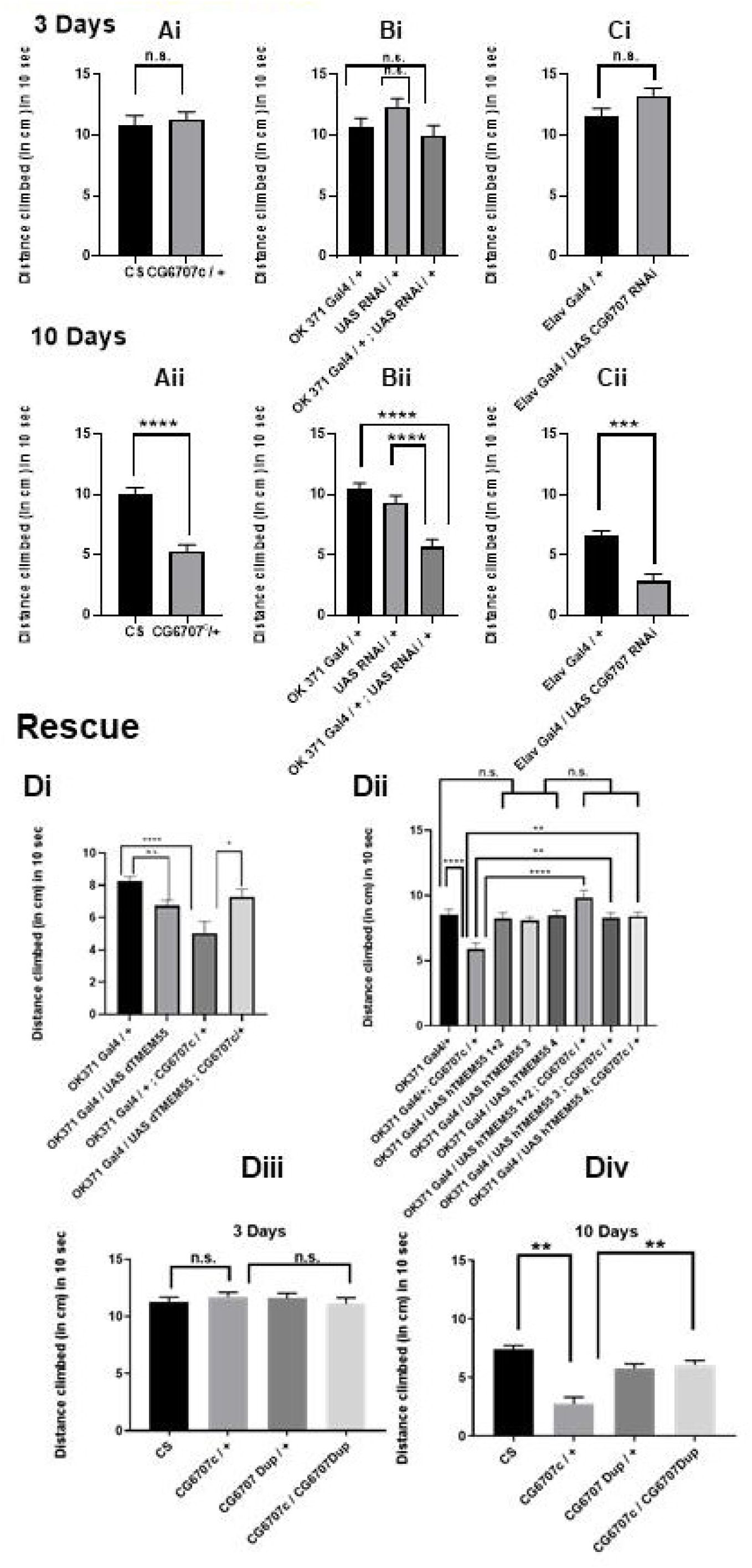
Loss of function of endogenous fly TMEM55 (dTMEM) leads to age-dependent locomotory defects and lethality. (Ai, Aii) Distance climbed by adult male flies (3 days and 10 days old) in 10 seconds compared between wild-type CS and loss of function piggyBac insertion dTMEM mutants (CG6707c or CG6707*^c01122^*), N > 16 flies. (Bi, Bii) Distance climbed by adult males (3 days and 10 days old) in 10 seconds compared in RNAi-mediated dTMEM knockdown flies in glutamatergic neurons with respect to the Gal4 driver and UAS responder alone, N > 16 flies. (Ci, Cii) Distance climbed by adult males (3 days and 10 days old) in 10 seconds compared in RNAi-mediated dTMEM knockdown flies only in the neurons with respect to the pan-neuronal Gal4 driver and UAS responder alone N > 16 flies. (Di) Overexpression of dTMEM in glutamatergic neurons rescues locomotory defects in CG6707c mutant flies, N = 11 males; (Dii) Overexpression of humanTMEM55 (driven by OK371 Gal4 driver and various UAS human TMEM55 lines) in glutamatergic neurons rescues locomotory defects in CG6707c mutant flies, N = 17 males. (Diii, Div): dTMEM duplication also leads to decline in the locomotory defects mediated by the loss of function of dTMEM in the CG6707c mutants, in age dependent manner. Genotypes: CS: Wild type; CG6707c / +: dTMEM heterozygous mutant; CGDup / +: heterozygous dTMEM duplication; CG6707c / CGDup: double transgenic with dTMEM mutation and dTMEM duplication, N = 17 males. *: p < 0.05; **: p < 0.01; ***: p < 0.001; n.s: p ≥ 0.05. p-value was calculated using Prism 7’s two-tailed Unpaired t-test and One-way ANOVA, multiple comparisons.

Notably, re-expression of dTMEM55 (OK371 Gal4 UAS *dTMEM55*; CG6707c /+) or human TMEM55B (OK371 Gal4 UAS *hTMEM55*; CG6707c /+) in the neurons of dTMEM55-deficient flies rescued the climbing defects [Figure 2Di–2Div]. This result indicates that the neuronal dysfunction phenotype is specifically associated with the loss of function of dTMEM55 and not any other gene. Additionally, rescue by the human TMEM55B indicates that the human TMEM55B shares similar functional attributes as dTMEM55. Therefore, the information obtained through the neuronal dysfunction phenotype from fly TMEM55B loss of function models can be extended to neuronal degeneration in human AD.

Lifespan or lethality assays in flies is another common read out for testing neuronal toxicity in evaluating any gene’s function in neurodegenerative disease (Nitta and Sugie 2022). We found that knockdown of *dTMEM55* in motor neurons enhanced lethality in 43 days old flies [OK371 Gal4/+; UAS CG6707 RNAi / +], Supplementary Figure 7A]. Lethality assay is one of the quantitative approaches to evaluate neurotoxicity and is widely used to assess the effect of various genes in modulating toxicity in various neurodegenerative diseases (Nitta and Sugie 2022). Therefore, these results conclude that TMEM55B potentiates neurodegeneration in AD by enhancing neuronal toxicity.

### Loss of dTMEM55 leads to accumulation of lysosomes in larval segmental nerve axons

Lysosomes are important for maintenance of protein and organelle homeostasis in cells. A lysosome’s optimum functioning is important for neuronal longevity and health. Maintenance of optimum axonal lysosomal transport is important for regulating neuronal development, its maintenance and functioning. It is also known that perturbations to optimal axonal lysosomes, or abnormal accumulation or decreased number of axonal lysosomes are linked to altered neuronal function and health. Any change in optimal axonal lysosome number has been known to lead to several neurological diseases including neurodegenerative diseases (Paumier and Gowrishankar 2024). Investigations in both human AD brain tissue and mouse models of AD have shown the excessive accumulation of lysosomes in swollen axons near extracellular amyloid aggregates, which formed part of the neuritic plaque (Terry et al. 1964), one of the major neuropathological hallmarks of AD. These multilamellar and multivesicular vesicles were observed by electron microscopy (Terry et al. 1964) and resembled axonal organelles that build up due to retrograde transport defects or blockage (Tsukita and Ishikawa 1980). Abnormal lysosome trafficking and signaling has been implicated in various diseases, including neurodegenerative diseases and various molecular players have been shown to have an important role in regulating these processes (Lie and Nixon 2019).

To assess whether downregulation of *dTMEM55* could lead to any lysosome transport defects, we assayed the number of lysosomes in the *Drosophila* larval segmental nerve axons. Lysosomes in these larval nerve axons were visualized by expressing HA-tagged LAMP1 in motor neurons. LAMP1, a lysosomal-associated membrane glycoprotein, serves as a canonical marker for lysosomes and autolysosomes and is widely employed to monitor autophagic flux and autophagosome–lysosome fusion. Its reliability is supported by structural, biochemical, and functional studies, including colocalization with LC3, lysosomal dynamics, and fusion events. LAMP1-based detection is formally recommended in the autophagy monitoring guidelines, reflecting its broad utility in assessing lysosomal integrity and autophagic progression across diverse cellular contexts (Klionsky et al. 2008a), (Klionsky et al. 2012), (Klionsky et al. 2016), (Klionsky et al. 2021), (Cheng et al. 2018). Deficiency of *dTMEM55* resulted in abnormal accumulation of lysosomes in the larval axons [OK371 Gal4;CG6707c + UAS Lamp1 HA, Figure 3Bi-ii, 3C,3D] in comparison to the control larval axons [OK371 Gal4 + UAS Lamp1 HA, Figure 3Ai-ii, 3C,3D]. These results indicate that lowered TMEM55B could lead to neurodegeneration by affecting normal axonal transport of lysosomes.

**Figure 3:**
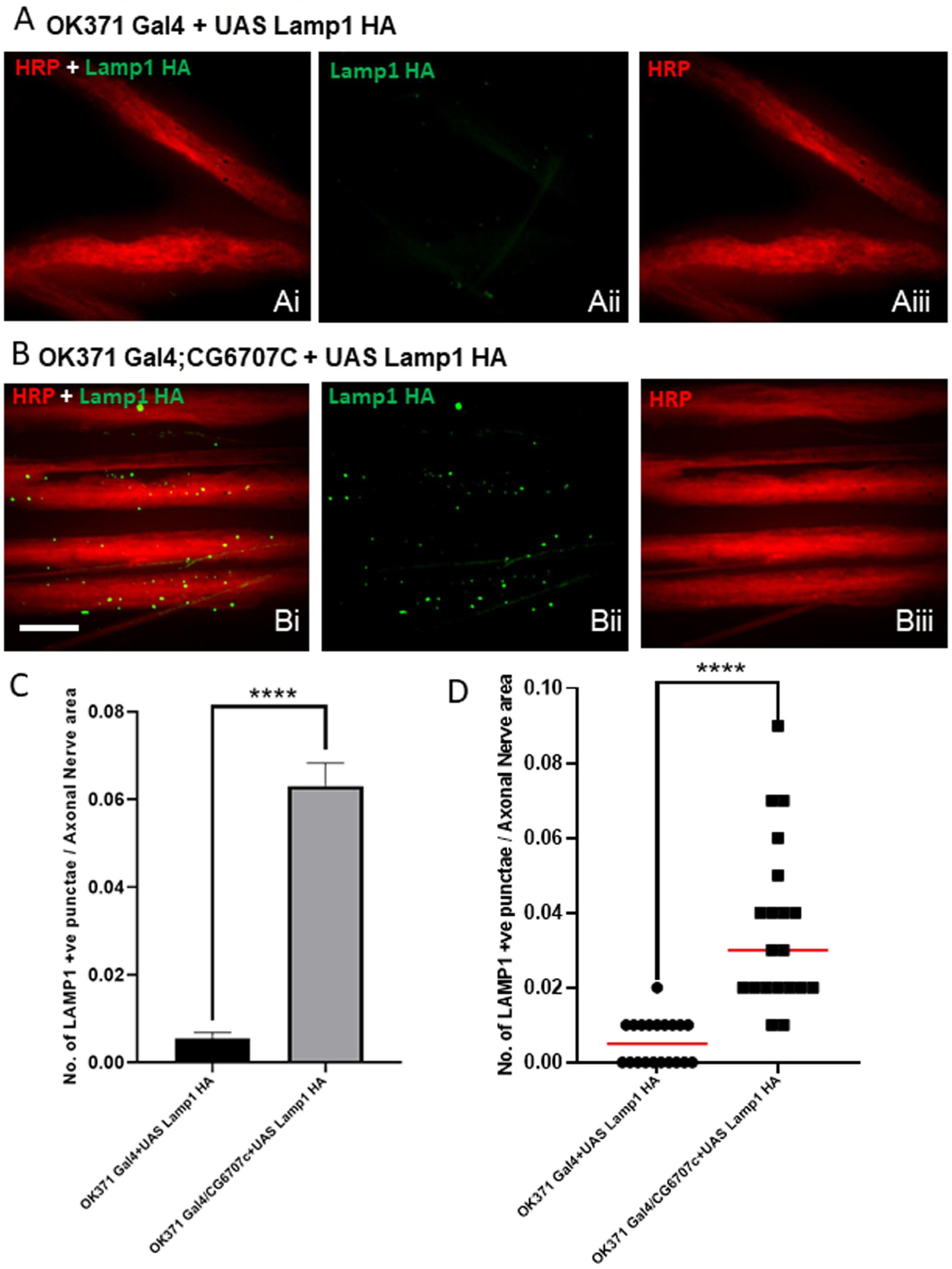
Loss of dTMEM leads to lysosomal accumulation in axons of larval segmental nerves. (Ai,Aii,Aiii,Bi,Bii,Biii) Confocal images of axons in the proximal (with respect to the ventral nerve cord)/anterior regions of the larval segmental nerve (A3-A4 segments) stained with antibodies against hemagglutinin or HA (green) and HRP (red). (Bi, Biii) Increased lysosomal accumulation (lysosome is tagged with HA) in the axons is observed in the dTMEM mutant larvae segmental nerves. (C): Bar graph with quantitation of lysosomal accumulation, Y axis: Total no. of lysosomal punctae / Axonal nerve area. (D): Scatter plot showing distribution of total number of lysosomal punctae per unit axonal nerve area in 20 axonal tracts. Each data point represents a single axonal tract. Genotypes: (Ai, Aii, Aiii) WT: OK6-GAL4/+; UAS hLAMP1 HA (Bi, Bii, Biii) OK6-GAL4/+; CG6707c / UAS hLAMP1 HA, [N = 20 axonal tracts across 4 animals]. *: p < 0.05; **: p < 0.01; ***: p < 0.001; n.s: p ≥ 0.05. p-value was calculated using Prism 7’s two-tailed Unpaired t-test. Scale bar, 20 µm.

### Downregulation of TMEM55B enhances Tau-mediated retinal degeneration, locomotory dysfunction, lethality and cell death

Age dependent neurodegeneration has been observed due to expression of β-amyloid (Aβ) or *Tau* encoded by the gene microtubule-associated protein tau (MAPT), both of which are associated with the prominent AD pathologies, in *Drosophila* brain (Iijima et al. 2004), (Finelli et al. 2004), (Crowther et al. 2005), (Wittmann et al. 2001). AD is an age dependent neurological disorder that causes irreversible loss of neurons, especially in the brain cortex and hippocampus. In the brains of AD patients, senile A beta plaques with aggregates of ß-amyloid peptides and neurofibrillary tangles (NFTs), comprised of aggregates of highly phosphorylated Tau protein, are observed. These are the major neuropathological hallmarks of AD (Scheltens et al. 2021). *Drosophila* is routinely used as a simple, genetically tractable in *vivo* model for studying the molecular pathogenesis of AD (Bouleau and Tricoire 2015), (Jeon et al. 2020).

Previously, Roy et al. (Roy and Jackson 2014), have used the rough eye phenotype as a readout to evaluate neurodegeneration to probe Tau and alpha Synuclein interaction. As a step forward, we wanted to evaluate whether downregulation of dTMEM55 modulates Tau toxicity. We downregulated dTMEM55 expression using a mutant dTMEM55 (heterozygous) fly [CG6707c/+] crossing it to a fly overexpressing human wild-type full-length (2N/4R) tau using a *gl*-Tau construct (human tau cDNA fused directly to the eye-specific *gl* promoter (Ambegaokar and Jackson 2011). *gl*-Tau misexpression causes a rough phenotype of the external eye associated with abnormalities of photoreceptor neurons and other cell types in the underlying retina (Jackson et al. 2002). In the *gl*-Tau flies, the degree of the rough eye phenotype is intermediate with mainly anterior patch of ommatidia being disrupted. These flies have reduced eyes, with the posterior region of the eye remaining more normal (Tau1.1 or *gl-*Tau 1.1 */ +*) Figure 4Ai). Misexpression of both human Tau with a concomitant downregulation of dTMEM55, resulted in a more severe rough eye, with the disrupted ommatidia extending farther posteriorly (Tau1.1 + dTMEM55 mut or *gl*-Tau1.1 /+; CG6707c / +, Figure 4Aii, 4Aiii). We also assayed the extent of rough eye phenotype in the *gl-*Tau flies, with the neuronal knockdown of dTMEM55. Additionally, we have shown that Tau mediated roughness increased significantly in eyes of flies with pan-neuronal *dTMEM55* knockdown (Figure 4Biv, 4Bv), in comparison to *gl-*Tau alone (Figure 4Biii).

**Figure 4:**
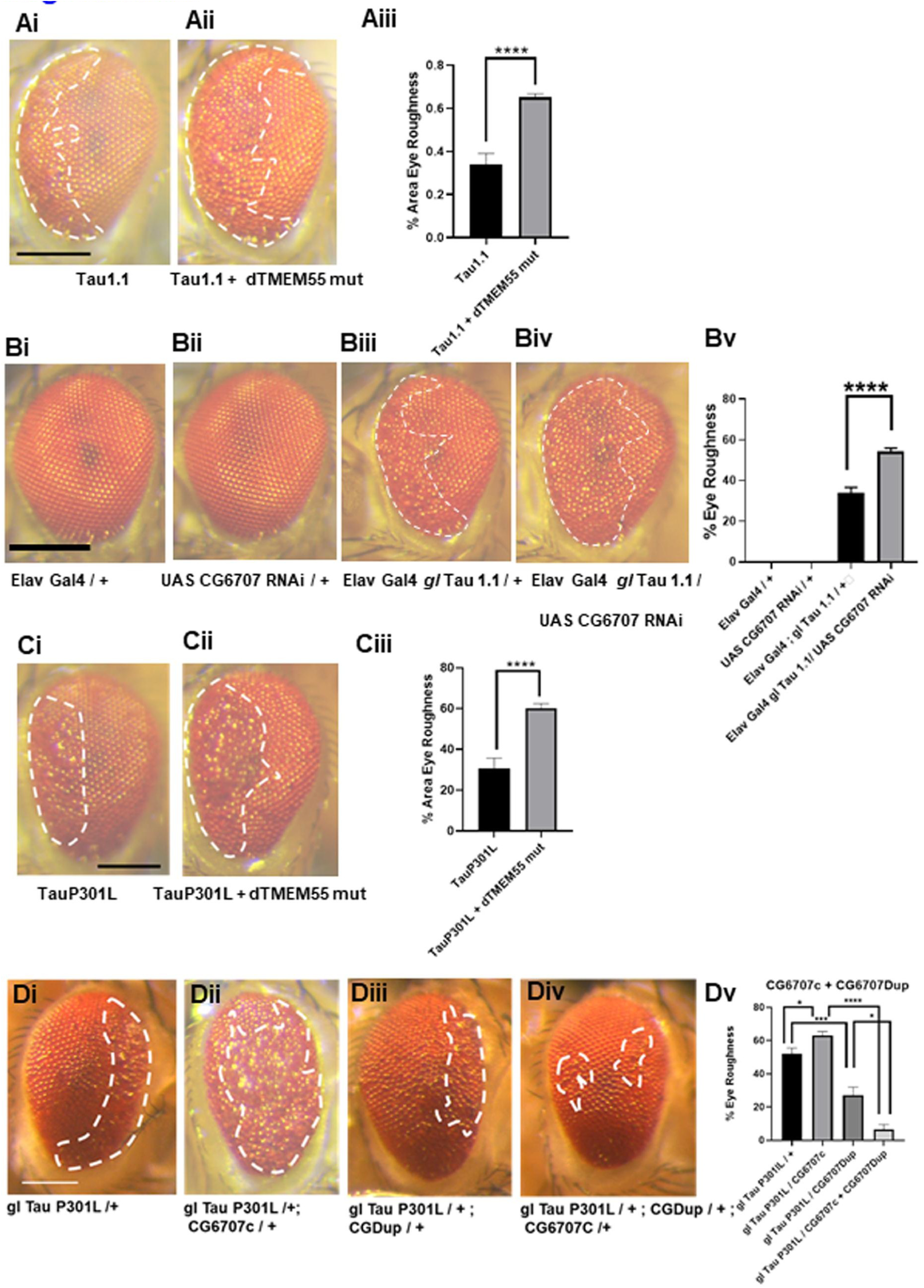
Downregulation of dTMEM enhances Tau mediated retinal degeneration. (Ai,Aii,Bi,Bii,Ci,Cii,Ciii,Civ) Light micrographs of fly eye to determine the extent of retinal degeneration or eye roughness. Downregulation of dTMEM enhances both wild type (gl Tau1.1) [Ai,Aii] and P301L mutant Tau (gl Tau p301L) [Bi,Bii] mediated retinal degeneration phenotypes. Downregulation of dTMEM specifically in the photoreceptor neurons using the pan-neuronal Elav Gal4 driver also enhances wild-type Tau-mediated retinal degeneration (Ci, Cii, Ciii, Civ). Aiii, Bi, Ciii: Bar graphs representing the extent of eye roughness in each of the genetically dTMEM downregulation strategies. Y axis: % eye roughness = [Area of roughness / Total Area of the retina] X 100.Genotypes: Ci: gl TauP301L / +; Cii: gl TauP301L / +; CG6707c / +; Ai: glTau1.1 / +; Aii: gl Tau1.1/+; CG6707c / +; Bi: Elav Gal4 / +; Bii: Elav Gal4 / +; UAS CG6707 RNAi / +; Biii: Elav Gal4; gl Tau1.1 / +; Biv: Elav Gal4 / +; gl Tau1.1 / +; UAS CG6707 RNAi / + [N > 5], *: p < 0.05; **: p < 0.01; ***: p < 0.001; n.s: p ≥ 0.05. p-value was calculated using Prism 7’s two-tailed Unpaired t-test and One-way ANOVA, multiple comparisons. Scale bar, 150 µm

We also observed that downregulation of dTMEM55 enhances eye roughness in flies mis-expressing *TauP301L* [a Tau mutation most frequently observed in patients with frontotemporal dementia with parkinsonism linked to chromosome 17 (FTDP-17), [TauP301L + dTMEM55mut or *gl*-TauP301L / +; CG6707c / +, Figure 4Cii, 4Ciii] in comparison to flies expressing TauP301L alone, without any *dTMEM55* downregulation [TauP301L or *gl*-TauP301L /+, Figure 4Ci, 4Ciii]. The specific role of *dTMEM55*’s downregulation in enhancing Tau mediated rough eye phenotype or retinal degeneration was shown by introducing an additional copy of *dTMEM55* through genomic fragment duplication [CGDup], which had the potential to restore the dTMEM55 to optimum levels, against the loss of function *dTMEM55* genetic background [*gl*-TauP301L / +; CGDup / +; CG6707c / +, Figure 4Div, 4Dv]. In these flies, the rough eye phenotype was significantly reduced [Figure 4Div, 4Dv] in comparison to flies where Tau was mis-expressed against a *dTMEM55* loss of function background [*gl*-TauP301L / +; CG6707c / +, Figure 4Diii, 4Dv]. All these results demonstrated that TMEM55B had the potential to enhance both wild type Tau and mutant Tau P301L phenotypes in a fly AD model, an indicator of its plausible role in potentiating Tau mediated neurodegeneration in Alzheimer’s disease. The severity of Tau-mediated eye roughness and retinal degeneration in transgenic flies with TMEM55B gene downregulation was quantitatively assessed across individual flies. Each data point in the scatter plot represents a single fly, allowing visualization of inter-individual variability in degenerative phenotypes rather than relying on averaged values alone (Supplementary Figure 5). The use of scatter plots enables detection of phenotypic heterogeneity, outliers, and the overall distribution of degeneration severity within each genotype, thereby providing a more accurate and biologically informative assessment of Tau-induced retinal degeneration at the individual-fly level.

Motor behaviour has been used to probe for neuronal function in various neurodegenerative models. Tau has previously been shown to be involved in axonal transport defects and locomotor dysfunction in a *Drosophila* model (Mudher et al. 2004). To examine whether downregulation of dTMEM55 modulates Tau mediated adult locomotor behaviour, we used the motor neuronal *OK371*-GAL4 driver to express Tau in postmitotic neurons. We observed that 21-day-old flies expressing Tau alone [OK371 Gal4 / UAS Tau 1.13] or the dTMEM55 (heterozygous) mutant [OK371 Gal4 / +; CG6707c / +] had severe motor deficits, with respect to the controls [Figure 5Aiii]. However, 21-day-old flies, misexpressing wild type Tau against a loss of function dTMEM55 genetic background [OK371 Gal4 / UAS Tau1.13; CG6707c / +] showed significantly higher motor behaviour defects than age matched flies expressing Tau alone or flies having only dTMEM55 downregulated. Scatter plots were used to assess Tau-mediated locomotory dysfunction following downregulation of TMEM55 by analyzing locomotor performance at the level of individual flies (Supplementary Figure 6). Each data point represents a single fly, allowing visualization of the full distribution of locomotory deficits rather than relying solely on mean values. This approach captures inter-individual variability, phenotypic heterogeneity, and potential outliers that may be obscured in simple bar graphs. Consequently, the use of scatter plots provides a more sensitive and biologically informative assessment of the extent to which TMEM55 downregulation enhances Tau-induced locomotory dysfunction.

**Figure 5:**
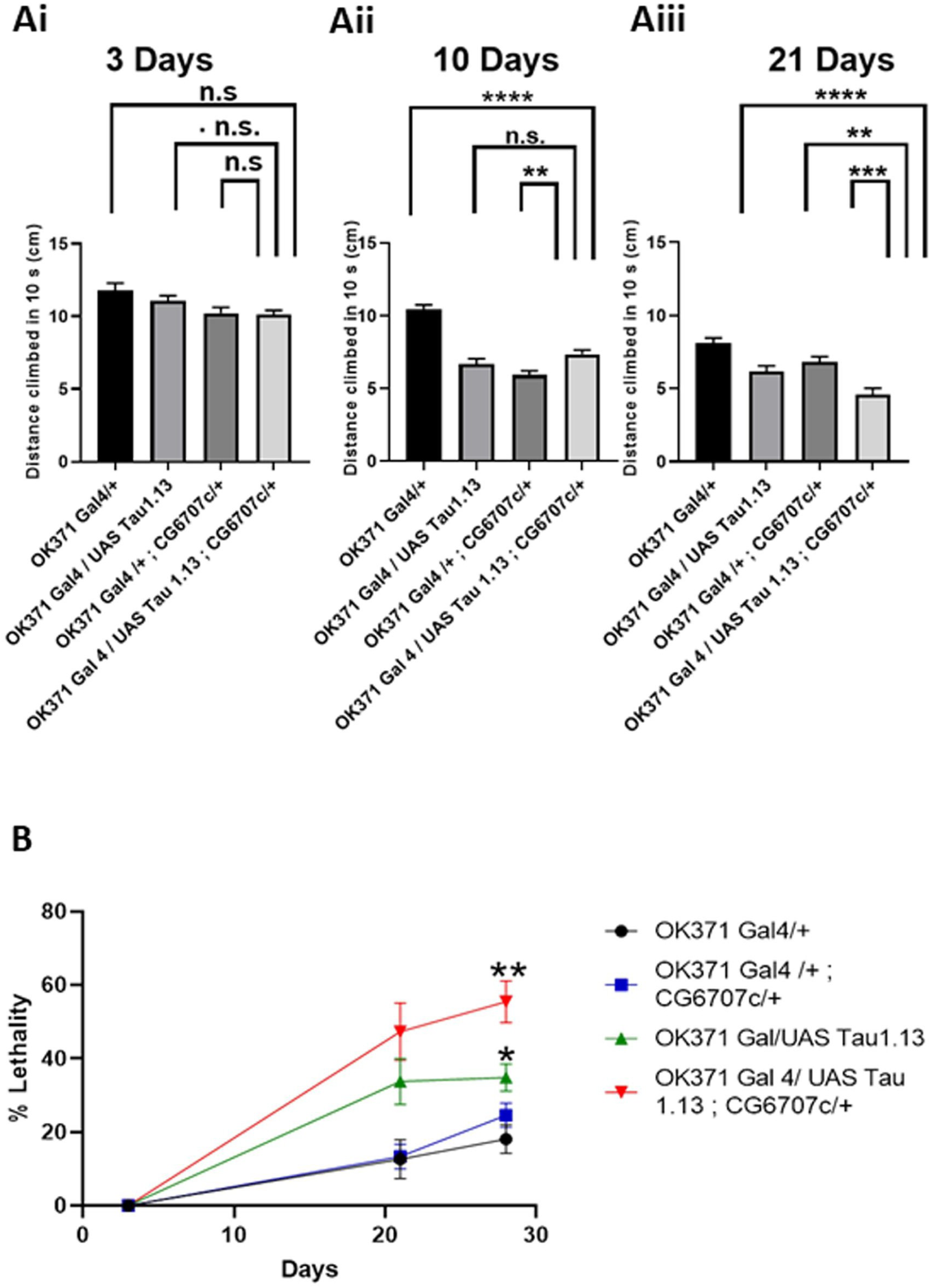
Downregulation of dTMEM enhances Tau-mediated locomotory dysfunction and lethality. (Ai, Aii, Aiii) Bar graphs representing the locomotory function of adult flies with a measurement of distance climbed by adult male flies (3 days, 10 days old, and 21 days) in 10 seconds compared between various genotypes, N = 17 males per genotype. (B) Line graphs depicting quantitative results of lethality assay comparing flies of various genotypes are listed below, N = 4 tubes [each tube = 10-12 flies]. *: p < 0.05; **: p < 0.01; ***: p < 0.001; n.s: p ≥ 0.05. p-value was calculated using Prism 7’s One-way ANOVA, multiple comparisons.

To assess whether downregulation of dTMEM55 enhances Tau mediated neuronal toxicity, we used fly lethality assay. Mis-expression of Tau alone using the motor neuronal driver [OK371 Gal4] enhanced lethality in 28-day-old flies [OK371 Gal4 / UAS Tau 1.13, Figure 5B]. However, this lethality was further exacerbated when Tau was misexpressed against a dTMEM55 loss of function genetic background (Figure 5B). This result demonstrates that downregulation of dTMEM55 enhances Tau mediated neuronal toxicity.

The TUNEL (Terminal deoxynucleotidyl transferase dUTP nick end labeling) assay is a widely employed method to detect DNA fragmentation, a hallmark of apoptotic cell death. In the context of Alzheimer’s disease (AD), TUNEL has been reliably used to identify neuronal apoptosis in both postmortem brain tissue and experimental models. Early studies demonstrated increased TUNEL-positive neurons in AD brains compared to age-matched controls, implicating apoptosis as a key contributor to neurodegeneration (Smale et al. 1995). More recent investigations continue to validate TUNEL as a robust tool for detecting cell death in AD, often in combination with complementary markers to confirm apoptotic pathways (Goel et al. 2022). Collectively, these studies support the use of the TUNEL assay as a reliable and sensitive method to assess neuronal loss in Alzheimer’s disease research. In *Drosophila* models of neurodegeneration, including tauopathy and amyloid-β overexpression models, TUNEL has been reliably used to quantify neuronal apoptosis. For example, flies expressing human tauR406W exhibit significantly increased TUNEL-positive neurons compared to controls, correlating DNA fragmentation with tau-mediated neurotoxicity (Wittmann et al. 2001). The TUNEL assay detects fragmented DNA, a hallmark of apoptosis, making it a sensitive and reliable method to identify dying cells in tissues. In the tauR406W *Drosophila* model, TUNEL staining specifically labelled apoptotic neurons in the brain, as confirmed by colocalization with the neuronal marker Elav and absence of signal in glial repo-expressing cells. This demonstrates that TUNEL can reliably distinguish cell types undergoing apoptosis. Moreover, changes in TUNEL-positive cell counts accurately reflected manipulations of oxidative stress pathways: downregulation of antioxidant defenses increased TUNEL-positive neurons, while antioxidant enzyme overexpression or vitamin E treatment decreased them (Dias-Santagata et al. 2007). Importantly, these changes occurred without altering tau protein levels, showing that TUNEL specifically reports the extent of apoptotic cell death, rather than changes in protein expression. Thus, TUNEL proved to be a quantitative and cell type–specific marker of apoptosis, enabling the study of neurodegenerative mechanisms and therapeutic interventions in this tauopathy model. These studies confirm that the TUNEL assay provides a robust and sensitive measure of neuronal cell death in *Drosophila* neurodegeneration models. We also observed that loss of dTMEM55 enhances Tau mediated cell death assayed by TUNEL staining (Supplementary Figure 9 Ai-iii, 9 B) in 21 days old fly brains.

All these assays, including retinal degeneration, climbing defects and lethality showed that the downregulation of dTMEM55 enhanced Tau mediated neuronal degeneration, dysfunction and toxicity in the fly transgenic AD models, indicating that these neuronal defects might lead to neurodegeneration in AD patients.

### Downregulation of dTMEM leads to elevated levels of Tau in fly brains

The microtubule-binding protein tau is a component of neurofibrillary tangles (NFT) in Alzheimer’s disease (AD) and related disorders that are collectively referred to as tauopathies (Morishima-Kawashima et al. 1995), (Johnson and Bailey 2002). In the NFT, Tau is hyperphosphorylated and it is widely believed that tau kinases such as glycogen synthase kinase-3β (GSK)-3β, microtubule affinity-regulating kinase (MARK) and cyclin-dependent kinase 5 (Cdk5) play a role in the formation of insoluble tau aggregates and NFT (Cho and Johnson 2003), (Augustinack et al. 2002). The phosphorylated Tau forms the NFT, which is one of the neuropathological hallmarks in AD. To assess whether downregulation of dTMEM55 leads to any change in the p-Tau (pSer202/pThr205) levels in our model of TMEM55B’s role in AD, we performed immunoblots with head extracts. We probed the blot with AT8, a monoclonal antibody that recognizes Tau phosphorylated at serine 202 and threonine 205, T-46, which detects total Tau, and GAPDH as a loading control. We detected an increase in AT8 signal in flies mis-expressing Tau along with downregulation dTMEM55 in comparison to flies only mis-expressing Tau alone without any dTMEM55 downregulation. This kind of elevated p-Tau levels was observed in flies with downregulated dTMEM55 [using the heterozygous *dTMEM55* mutant, CG6707c / +] with either wild type Tau [Tau 1.1 or *gl-*Tau1.1] or mutant Tau [P301L Tau or *gl*-P301L Tau] mis-expression [Figure 6A, 6Aiii (*gl*-Tau1.1 + CG6707c /+), Figure 6B, 6Biii (*gl*-Tau1.1 / +; CG6707c / +)]. This indicates that loss of function dTMEM55 elevates phosphorylated Tau levels, a clinical-pathological manifestation known to be associated with neurofibrillary tangles in AD.

**Figure 6:**
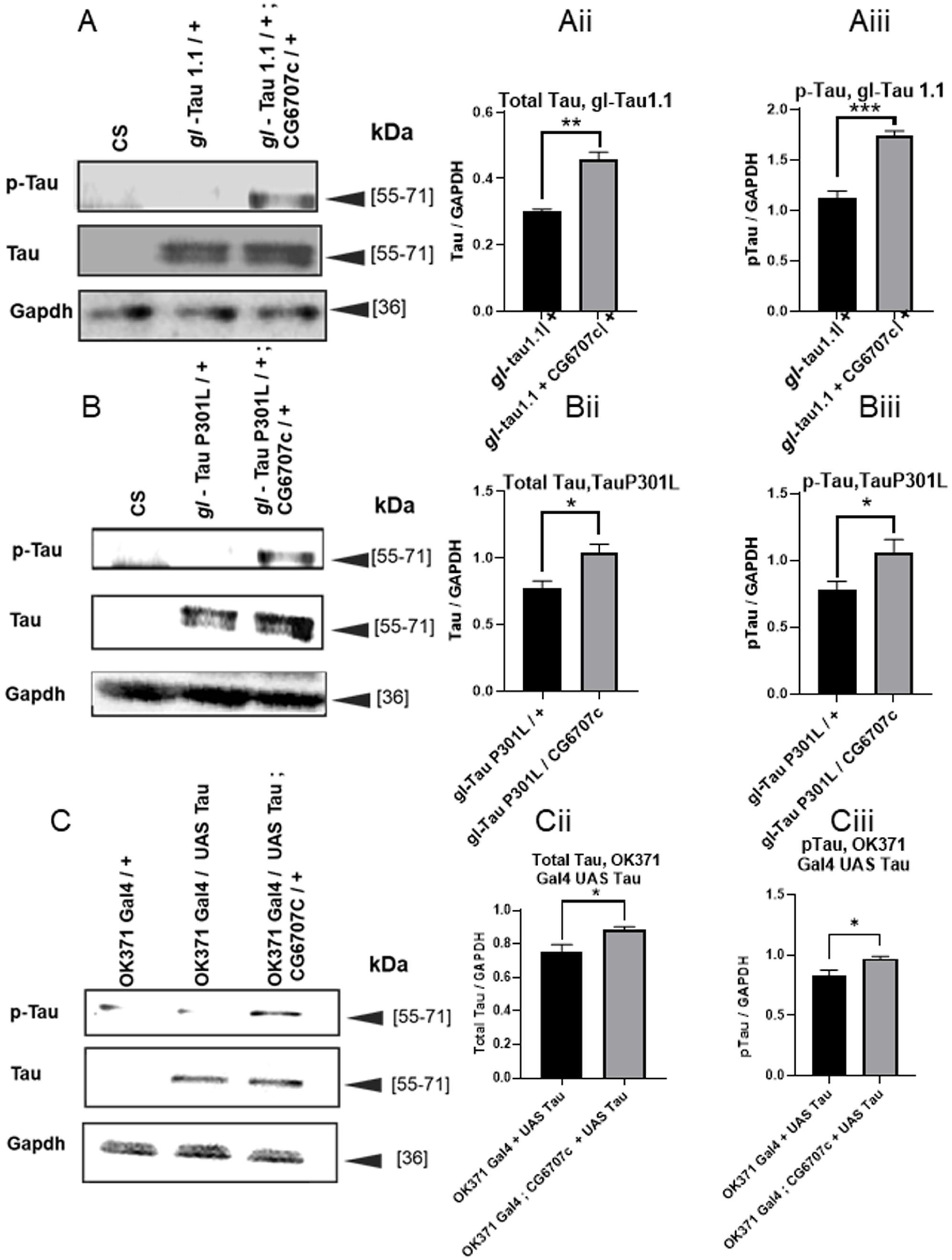
Downregulation of dTMEM leads to elevated levels of Tau in fly brains. (Ai, Aii): Bar graphs of immunoblots for total Tau and phosphorylated Tau (AT8 specific epitopes) in dTMEM downregulated flies with overexpressing wild-type Tau. (Aiii) Representative blot images of Tau overexpressing flies (+) and dTMEM downregulated flies with overexpressing P301L mutant Tau (glTau1.1 / +; CG6707c / +). (Bi, Bii): Bar graphs of immunoblots for total Tau and phosphorylated Tau (AT8 specific epitopes) in dTMEM downregulated flies with overexpressing P301L mutant Tau. (Biii) Representative blot images of Tau overexpressing flies (glTau P301L / +) and dTMEM downregulated flies with overexpressing P301L mutant Tau (glTauP301L / +; CG6707c / +). (Ci, Cii): Bar graph of immunoblots for total Tau in dTMEM downregulated flies with Tau being overexpressed in a few brain neurons derived by the OK371 Gal4 driver. (Ciii) Representative blot images of Tau overexpressing flies (OK371 Gal4 / UAS Tau 1.1) and dTMEM downregulated flies with Tau being overexpressed in few brain neurons by the OK371 Gal4 driver. (OK371 Gal4 / UAS Tau 1.1; CG6707c / +). N=3 blots, *: p < 0.05; **: p < 0.01; ***: p < 0.001; n.s: p ≥ 0.05. p value was calculated using Prism 7’s two tailed Unpaired t-test and One way Anova, multiple comparison.

To assess whether downregulation of dTMEM55 leads to an increase in AT8 signal in the neurons, we misxpressed human Tau using an enhancer trap line OK371-Gal4 [which expresses the desired protein in a defined subset of neurons from embryonic stage 15 to adulthood] in the dTMEM55 heterozygous mutant background. We detected an increase in AT8 signal in flies’ mis-expressing Tau along with downregulation dTMEM55 in comparison to flies only mis-expressing Tau alone without any dTMEM55 downregulation. This kind of increased p-Tau levels was observed in flies with downregulated dTMEM55 [using the heterozygous *dTMEM55* mutant, CG6707c / +] with wild type human Tau [UAS Tau 1.1] being expressed in neurons using the OK371 Gal4 driver [OK371 Gal4 / UAS Tau 1.1; CG6707c / +], Figure 6C, 6Ciii. This indicates that loss of function dTMEM55 elevates phosphorylated Tau levels in the neurons, a clinical-pathological manifestation known to be associated with the formation of neurofibrillary tangles in AD. Thus, downregulation of TMEM55b could be potentially leading to elevated p-Tau, whose excessive accumulation could lead to formation of NFTs, leading to neuronal degeneration in AD.

### TMEM55B is lowered in iPSC derived neurons derived from AD patients

#### A. Establishment and Characterization of an iPSC-Derived Neuronal Model of Alzheimer’s Disease

Familial Alzheimer’s disease (FAD) accounts for fewer than 5% of AD cases; however, sporadic and familial forms share substantial phenotypic and molecular overlap, likely reflecting convergence on common pathogenic pathways (Duara et al. 1993; Lippa et al. 1996). Among known FAD mutations, variants in presenilin-1 (PSEN1) are the most prevalent, with the M146L mutation being one of the best-characterized disease-causing alleles (Mangone et al. 1995; Morelli et al. 1998). To investigate the role of TMEM55B within a genetically defined human AD context, we established an iPSC-derived neuronal model using an isogenic pair of female cell lines: a wild-type control and a heterozygous PSEN1 M146L mutant generated by CRISPR/Cas9 editing (Hurley et al. 2023), (Reddy et al. 2016). Presenilin-1 (PSEN1), the catalytic core of the γ-secretase complex, mediates intramembrane cleavage of the amyloid precursor protein (APP), generating amyloid-β (Aβ) peptides that accumulate as extracellular plaques, a defining neuropathological hallmark observed in all postmortem Alzheimer’s disease (AD) brains (Haass and Selkoe 1993), (De Strooper et al. 2012). To date, over 300 pathogenic mutations in PSEN1 have been identified in association with familial Alzheimer’s disease (fAD), underscoring its central role in disease pathogenesis. Among these, the M146L mutation represents one of the most frequently occurring variants, whereas the L435F mutation, although rare, is associated with a particularly aggressive clinical phenotype characterized by extensive deposition of Aβ43-enriched cotton-wool plaques (Morelli et al. 1998), (Rogaeva et al. 2001). Functionally, the PSEN1 M146L mutation has been shown to enhance γ-secretase activity toward non-canonical substrates, including increased cleavage of the endoplasmic reticulum calcium sensor STIM1, thereby implicating dysregulated calcium homeostasis in disease progression (Tong et al. 2016). Moreover, studies utilizing human induced pluripotent stem cell (iPSC)-derived neurons demonstrate that the M146L mutation significantly elevates the Aβ42/Aβ40 ratio by selectively increasing Aβ42 production without altering Aβ40 levels, a biochemical signature strongly linked to amyloid aggregation and neurotoxicity in AD (Liu et al. 2014), (Sproul et al. 2014), (Schrank et al. 2020). Neuronal differentiation was achieved using NGN2-mediated transcription factor programming (Zhang et al. 2013b), a strategy that enables rapid and efficient generation of functionally mature neurons while preserving robust expression of mature neuronal markers (Ho et al. 2016; Hulme et al. 2022). This approach was selected over neural progenitor cell–based differentiation to accelerate neuronal maturation while maintaining disease relevance. The differentiation protocol was further optimized by integrating established principles of neuronal development to ensure reproducibility and high neuronal purity (Lancaster et al. 2013; Lancaster and Knoblich 2014a; Lancaster and Knoblich 2014b). Neuronal cultures were analyzed at two defined maturation stages, day 20 and day 40 in vitro, to model early and later disease-relevant states.

To validate cell identity and differentiation efficiency, iPSCs and derived neurons were analyzed by Western blotting and immunocytochemistry for pluripotency and neuronal markers. Both control and PSEN1 M146L iPSCs expressed high levels of OCT3/4, which was absent in day 20 and day 40 neurons, confirming loss of pluripotency upon differentiation (Supplementary Figure 10 B). Conversely, mature neuronal markers including class III β-tubulin (TUJ1), tau, and synapsin were robustly expressed in neurons at both time points and were undetectable in iPSCs, demonstrating successful neuronal lineage commitment. Morphological assessment and immunocytochemical analyses further confirmed the generation of a high proportion of mature neurons, underscoring the robustness and consistency of the NGN2-driven differentiation protocol. Previous invivo studies from our group demonstrated that TMEM55B depletion exacerbates tau-mediated neurodegeneration, leading to enhanced retinal degeneration, locomotor deficits, increased lethality, and neuronal cell loss in both wild-type tau and Tau P301L models (Figures 4 and 5). These findings position TMEM55B as a modulator of Tau-dependent neurotoxicity and implicate it in pathways governing Tau phosphorylation and clearance. Given the central role of Tau pathology in AD and the conserved relationship between TMEM55B loss and Tau toxicity across model systems, we prioritized Tau-related endpoints in the analysis of PSEN1 M146L iPSC-derived neurons. Consistent with this rationale, day 20 neurons exhibited mature neuronal morphology but did not display overt AD-associated phenotypes (Figure 9B, Biv). In contrast, day 40 neurons showed robust accumulation of phosphorylated Tau (p-Tau), reflecting age-dependent pathogenic changes characteristic of AD (Figure 9Biii, Bv). These observations validate the temporal disease relevance of the model and establish a framework for interrogating TMEM55B-dependent mechanisms underlying Tau pathology in human neurons.

#### b. TMEM55B is selectively reduced in late-stage AD iPSC-derived neurons

Using our iPSC-derived neuronal model of Alzheimer’s disease, which captures early and late stages of pathology, we examined TMEM55B expression at day 20 and day 40 of neuronal maturation. Western blot and immunocytochemistry analyses revealed no difference in TMEM55B levels between PSEN1 M146L and isogenic control neurons at day 20, indicating that TMEM55B is preserved during early stages of neuronal differentiation (Figure 7Ai–Bi, 7A; Figure 9B, 9Bi). In contrast, by day 40, TMEM55B expression was significantly reduced in PSEN1 M146L neurons relative to controls (Figure 7Ci–Di, 7A; Figure 9B, 9Bii). Immunofluorescence co-labeling with MAP2 confirmed these findings, showing diminished TMEM55B signal in aged AD neurons while nuclear staining (DAPI) remained consistent. These observations parallel reductions in TMEM55B mRNA reported in human neuropathological datasets from the ROSMAP cohort (Figure 1), demonstrating that TMEM55B downregulation occurs at both transcript and protein levels and manifests during later stages of neuronal maturation, consistent with its temporal association with AD progression.

**Figure 7:**
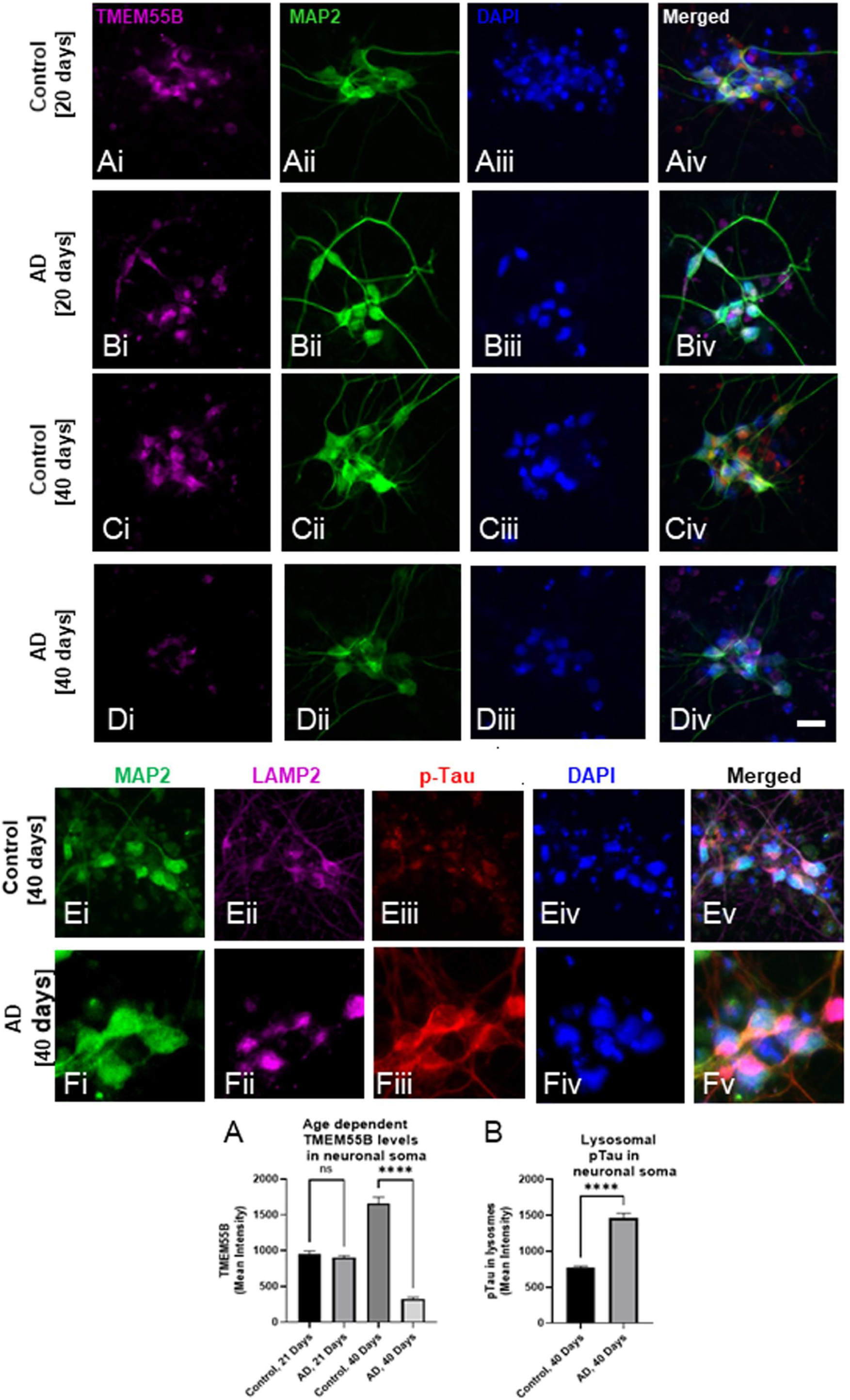
Age-dependent reduction of TMEM55B and somatic lysosomal accumulation of phosphorylated Tau in iPSC-derived Alzheimer’s disease neurons. (A) Quantitative immunofluorescence of iPSC derived differentiated human forebrain neurons of various ages were stained for MAP2, phospho-Tau, TMEM55B and DAPI. TMEM55b levels were lowered in the neuronal soma in the 40 days old AD forebrain neuronal culture. No significant changes were observed in the TMEM55B levels between 20 days old Control and AD patient’s iPSC derived forebrain neurons. N= 64 neurons. (B) Quantitative immunofluorescence of iPSC derived differentiated 40 days old human forebrain neurons were stained for MAP2, phospho-Tau, LAMP2 and DAPI. Phosphorylated Tau levels were elevated in the perinuclear lysosomal compartment of the neuronal soma in these 40 days old iPSC derived AD forebrain neurons. N= 48 neurons. *: p < 0.05; **: p < 0.01; ***: p < 0.001; n.s: p ≥ 0.05. p value was calculated using Prism 7’s two tailed Unpaired t-test and One way Anova, multiple comparison [Bonferroni]. Scale Bar: 20 µm

### Induced Pluripotent Stem Cells derived AD neurons show elevated phosphorylated Tau in the lysosomes located in the neuronal soma

To determine whether lowering TMEM55B in the aged AD neurons leads to any changes in the phosphorylated form of Tau in the lysosomes of these neurons we immunostained cells for p-Tau and LAMP2. *LAMP2* (along with LAMP1) is widely accepted as a structural lysosomal membrane protein marker used to identify lysosomes and late endo/lysosomal compartments in studies of autophagy and lysosome biology (Klionsky et al. 2021), (Eskelinen et al. 2002), (Saftig and Klumperman 2009). The formation of phosphorylated form of Tau and its accumulation is associated with the formation of neuropathological hallmarks (NFTs) in the AD brains. Therefore, an accumulation of p-Tau in the lysosomes upon downregulation of TMEM55B suggests defects in lysosomal mediated autophagic clearance of p-Tau, leading to neuronal degeneration by abnormal buildup of pathological forms of Tau.

We observed that an increase in p-Tau levels in the lysosomes located in the soma of the 40 day old AD iPSC derived forebrain neurons [Figure 7Fiii, 7B] in comparison to p-Tau levels in the soma of 40 days old Control neurons [Figure 7Eiii, 7B], [Figure 9Biii, 9Bv] and not in 20 days old AD iPSC neurons [Figure 9Biii, 9Biv]. These results in the iPSC derived neurons, suggests a possible defect in lysosome mediated autophagic clearance of p-Tau in the aged and severe AD patients, leading to neurodegeneration.

### TMEM55B loss in the iPSC derived AD neurons, lead to the accumulation of axonal lysosomes with a concomitant increase in the number of phospho-Tau positive lysosomes in these axons

Studies in both human AD brain tissue and mouse models of AD have revealed the massive accumulation of lysosomes in swollen axons near extracellular amyloid aggregates, forming part of the neuritic plaque (Terry et al. 1964). These organelles were observed to be multi-lamellar and multivesicular in nature by electron microscopy studies (Terry et al. 1964), indicative of the axonal organelles that build up due to retrograde transport blockade (Tsukita and Ishikawa 1980). These organelles were found to be relatively deficient in multiple lysosomal proteases and may arise due to a failure of optimal retrograde axonal lysosome transport near the amyloid plaques. Puertollano and coworkers (Willett et al. 2017) found that the lysosomal transmembrane protein TMEM55B recruits JIP4 to the lysosomal surface, inducing dynein dependent transport of lysosomes toward the microtubules minus-end. TMEM55B over-expression causes lysosomes to collapse into the cell center, whereas depletion of either TMEM55B or JIP4 results in dispersion toward the cell periphery. They showed that TMEM55B promotes lysosomal retrograde transport.

In order to analyse if TMEM55B loss in the iPSC derived AD neurons, lead to the accumulation of lysosomes in the axons, we analysed the LAMP2 staining in the axons of these neurons. We observed that there was a significant increase in the total number of lysosomes in the 40 days old iPSC derived AD neurons [Figure 8 Bi-Biii] in comparison to 40 days old iPSC derived control neurons [Figure 8 Ai-Aiii], Figure 8E. There is also an increase in the number of phospo-Tau positive lysosomes in these axons in 40 days old iPSC derived AD neurons [Figure 8 Di-Diii] in comparison to 40 days old iPSC derived control neurons [Figure 8 Ci-Ciii], Figure 8F. These results in the iPSC derived AD neurons, suggest possible lysosomal transport defects along the axons of the neurons. This in turn could probably lead to autophagic clearance defects, leading to neurodegeneration in aged and severe AD patients

**Figure 8:**
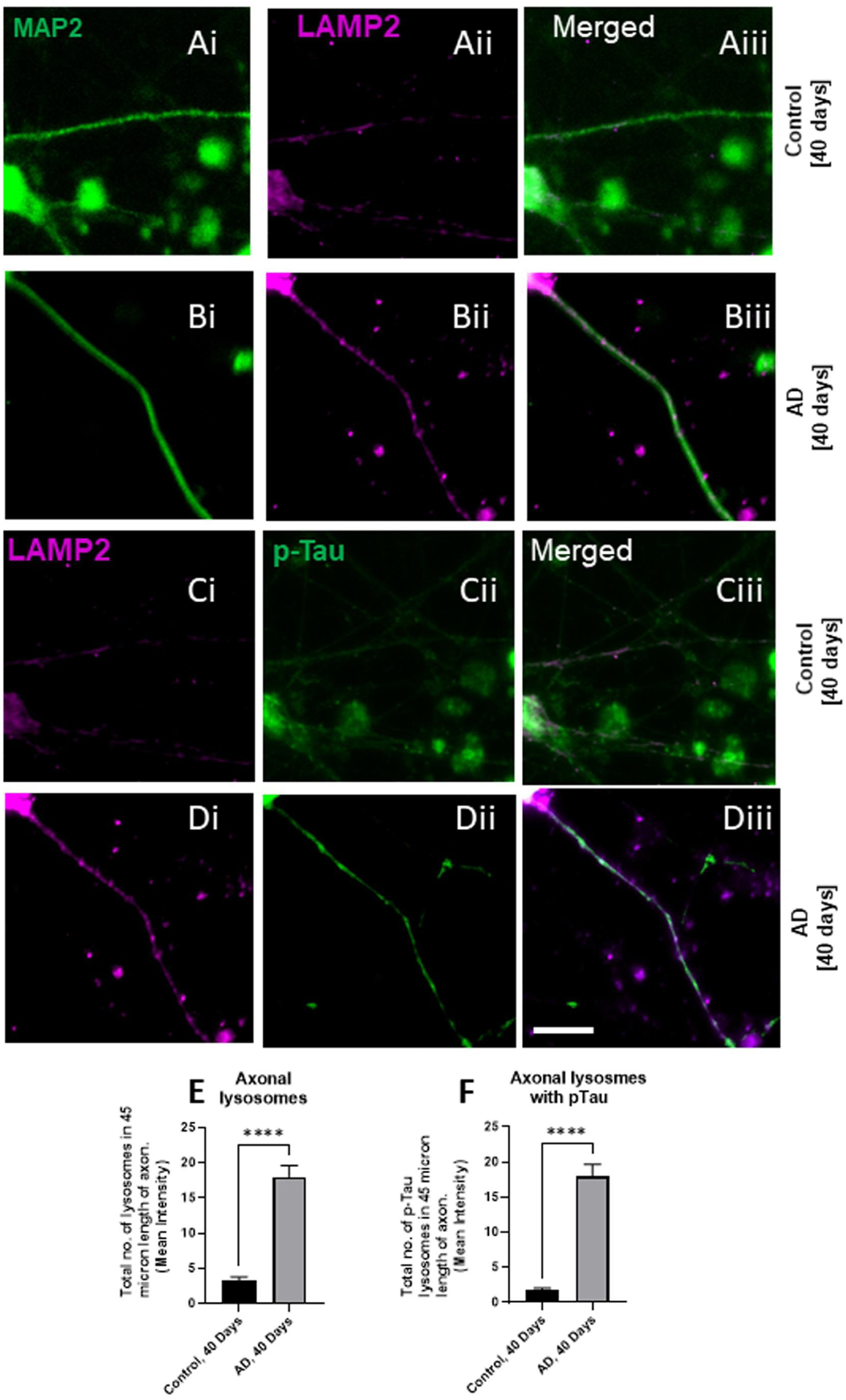
TMEM55B loss in iPSC-derived Alzheimer’s disease neurons promotes axonal lysosome accumulation and increased phosphorylated tau–positive lysosomes in the axons. (A) Quantitative immunofluorescence of iPSC derived differentiated 40 days old human forebrain neurons were stained for MAP2, phospho-Tau, LAMP2 and DAPI. There was an increase in the number of lysosomes (stained by LAMP2) in the axons of these iPSC derived AD patient’s forebrain neurons. Total number of lysosomes were counted by Fiji’s multipoint tool along the fixed 45-micron length of a single axon. (B) Quantitative immunofluorescence of iPSC derived differentiated 40 days old human forebrain neurons were stained for MAP2, phospho-Tau, LAMP2 and DAPI. Total number of lysosomes, which also stained positive for phospho-Tau, were counted by Fiji’s multipoint tool along the fixed 45-micron length of a single axon. An increase in the number of lysosomes (stained by LAMP2) also staining positive for phosphorylated Tau was observed in the axons of 40 days old, iPSC derived AD patient’s forebrain neurons, in comparison to the axons of the iPSC derived control individual’s forebrain neurons. N = 21 axons from different neurons. *: p < 0.05; **: p < 0.01; ***: p < 0.001; n.s: p ≥ 0.05. p-value was calculated using Prism 7’s two tailed Unpaired t-test. Scale Bar: 20 µm.

### Autophagic flux defects and age dependent abnormal accumulation of pTau, and lowering of TMEM55 levels in iPSC-derived AD neurons

Autophagic flux is used to represent the dynamic process of autophagy and includes the whole process of autophagy, including auto phagosome formation, maturation, fusion with lysosomes, subsequent breakdown and the release of macromolecules back into the cytosol (Zhang et al. 2013a). The most straightforward method for monitoring autophagy is the detection of LC3 protein processing. Autophagic flux can be detected by LC3-II turnover using Western blot analysis in the presence and absence of lysosomal degradation inhibitors, such as pepstatin A, E64d, bafilomycin A1, chloroquine and NH4Cl (Ju et al. 2010), (Klionsky et al. 2008b). If autophagic flux is occurring, the level of LC3-II will be increased in the presence of a lysosomal degradation inhibitor because the transit of LC3-II through the autophagic pathway will be blocked. Chloroquine (CQ) is widely used to inhibit autophagic flux; however, its precise mechanism was clarified by Mauthe et al. (2018), who demonstrated that CQ primarily blocks autophagosome–lysosome fusion rather than simply alkalinizing lysosomes. CQ disrupts SNARE-dependent membrane fusion, leading to accumulation of LC3-II–positive autophagosomes and p62, indicative of defective cargo degradation (Mauthe et al. 2018). Extending these findings, Nnah et al. (2019) identified key endocytic regulators of autophagic flux, showing that disruption of endosomal trafficking phenocopies CQ-induced fusion blockade and exacerbates neurodegeneration (Kodani et al. 2019). Together, these studies establish endolysosomal trafficking and fusion as critical determinants of late-stage autophagic flux. We evaluated the extent of autophagic flux in iPSCs and iPSC derived neurons in the presence and absence of lysosomal degradation inhibitor, chloroquine by evaluating LC-II and LC-I levels. We observed an increased LC-II levels in the presence of chloroquine in the iPSC derived 20 and 40-day-old AD neurons [Fig 9 A’ii and A’iii]. However, no change in the levels of LC-II levels was observed in the presence of chloroquine in the iPSCs [Fig 9 A’i]. We observed a robust, statistically significant decline in autophagic flux in the 20-day and 40-day old iPSC derived AD neurons in comparison to the age matched iPSC derived Control neurons [Fig 9 Aii and Aiii]. There was no autophagic flux defects in the AD iPSCs and the levels of autophagic flux in the AD iPSCs were comparable to the Control iPSCs [Fig 9 Ai]. Our results indicates that autophagic flux defects are prominent in AD neurons. These results additionally establish that these neurons could be potentially used for studying the effects of various lysosomal proteins in mediating autophagic flux to enhance clearance of Tau and other proteins known to form toxic aggregates in neurodegenerative diseases.

**Figure 9:**
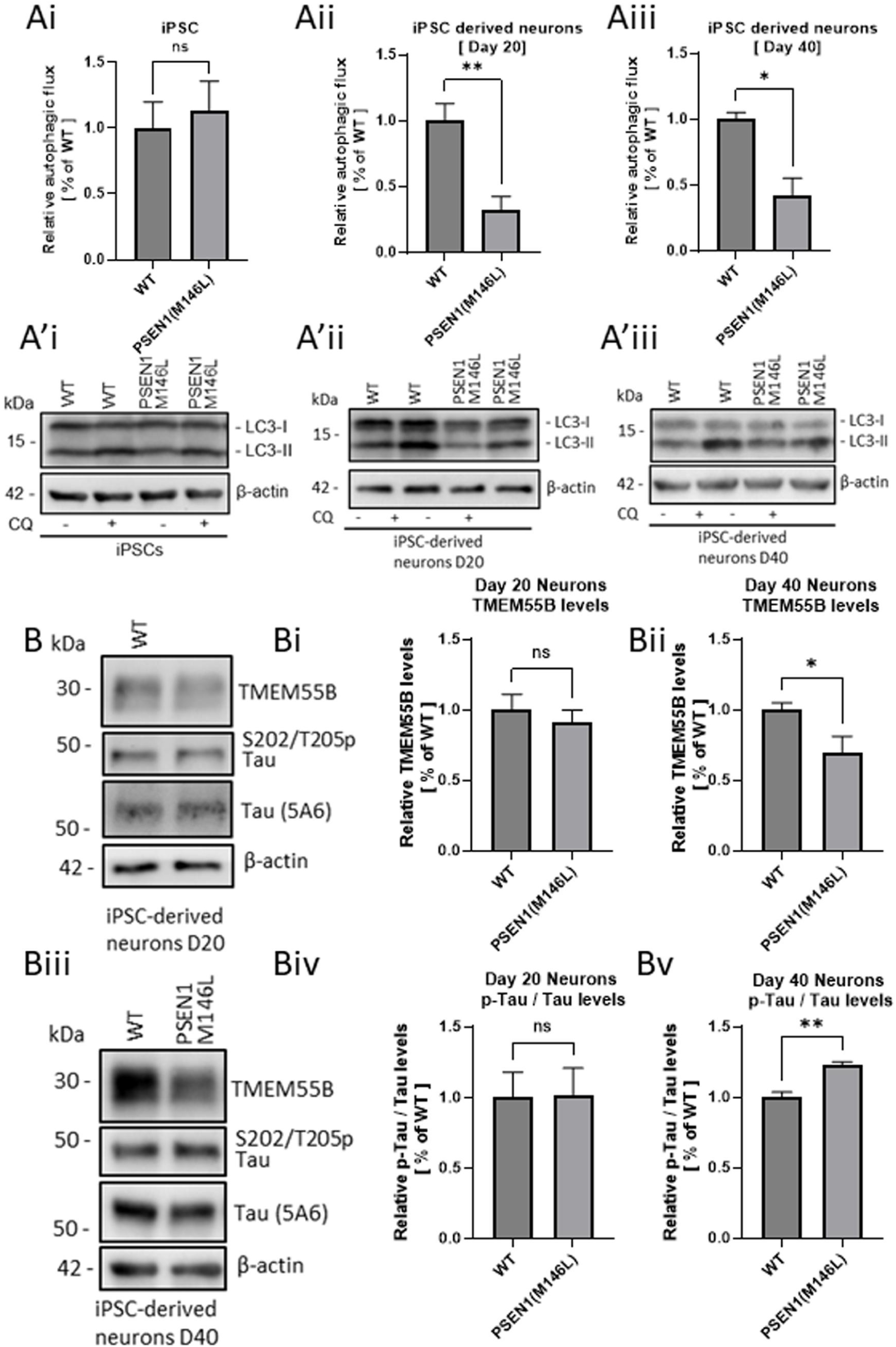
Progressive Loss of TMEM55B Is associated with Autophagic Flux Defects and accumulation of Phosphorylated Tau in iPSC-Derived Alzheimer’s Disease Neurons. Western blots measuring autophagic flux in iPSCs (Ai, A’i); 20 days old (Aii, A’ii) and 40 days old iPSC derived forebrain neurons from Control and AD patients (Aiii, A’iii). Immunoblots depicting the changes in TMEM55B levels (B, Bi, Bii) and phosphorylated Tau levels (Biii, Biv, Bv) in 20 days and 40 days old iPSC derived Control and AD neurons. N = 4 blots,*: p < 0.05; **: p < 0.01; ***: p < 0.001; n.s: p ≥ 0.05. p value was calculated using Prism 7’s two tailed Unpaired t-test and One way Anova, multiple comparison.

### Loss of TMEM55 leads to lysosomal positioning defects in HEK293T cells

Recently, Puertollano and co-workers (Willett et al. 2017), found a novel mechanism of lysosome positioning regulation. They found that the lysosomal protein TMEM55B interacts and recruits the dynein adaptor JIP4 to lysosomal membranes, thus inducing retrograde transport of lysosomes along microtubules. They also found that reduction of either TMEM55B or JIP4 caused a dramatic accumulation of lysosomes at the cell periphery without affecting the distribution of early endosomes or the Golgi apparatus. Since TMEM55B is a target of TFEB and TFE3, they proposed that, by regulating TMEM55B, expression levels [either by TFEB/3 activation by starvation or cholesterol accumulation in lysosomes]; TFEB and TFE3 modulate lysosomal positioning in response to nutrient deprivation and lysosomal stress.

To assess the distribution of lysosomes or late endosomes in si-Control HEK293T cells and si-RNA mediated TMEM55B knockdown HEK293T cells. In the ICC of these cells, we used antibody against Rab7 to probe for the compartmental localization of late endosomes and lysosomes, and evaluated TMEM55B levels by counterstaining with an anti-TMEM55B antibody. We found that TMEM55B was significantly reduced in the si-TMEM55B cells in comparison to the control cells [Supplementary Figure 11Ai-Avi, B]. The Rab7 staining was more localized more towards the peripheral regions of the si-TMEM55B HEK293T cells (Supplementary Figure 11Ax-Axii, C). On the other hand, the Rab7 staining was more localized more in the peri-nuclear regions of the si-Control HEK293T cells (Supplementary Figure 11Avii-Aix, C). These findings support the results observed by Puertollano and co-workers, and thus hint towards a possible mechanism of TMEM55B downregulation in AD patients, leading to lysosome positioning defects. These lysosome position defects could explain the continued presence of lysosomes and late endosomes in the peripheral and not the perinuclear region, preventing their fusion to the autophagosomes and impaired clearance of p-Tau leading to its excessive cellular accumulation.

### dTMEM55 interacts with lysosomal cation channel Trpml to regulate locomotor function, cell death, lethality and lifespan

Free Ca2+ stored within lysosomes, is a significant source of intracellular Ca2+ comparable to stores in the ER (Patel and Docampo 2010), is emerging as an important signal that coordinates adaptive responses upon its release into the cytosol. Lysosomal Ca2+ efflux via TRPML1 is activated in several stress conditions, including starvation (Li et al. 2016), (Medina et al. 2015), lysosome deacidification (Lee et al. 2015), (Raychowdhury et al. 2004) and oxidative stress (Zhang et al. 2016), as well as during physiological processes such as phagocytosis of large particles by macrophages (Samie et al. 2013). Ballabio, Medina and coworkers (Medina and Ballabio 2015), (Medina et al. 2015) meanwhile showed that lysosomal Ca^2+^ signaling controls the activities of the phosphatase calcineurin and of its substrate TFEB, a master transcriptional regulator of lysosomal biogenesis and autophagy. Lysosomal Ca^2+^ release via mucolipin 1 (MCOLN1) activates calcineurin, which binds and dephosphorylates TFEB, thus promoting its nuclear translocation. Genetic and pharmacological inhibition of calcineurin suppressed TFEB activity during starvation and physical exercise, while calcineurin overexpression and constitutive activation had the opposite effect. Induction of autophagy and lysosomal biogenesis via TFEB required MCOLN1-mediated calcineurin activation, linking lysosomal calcium signaling to both calcineurin regulation and autophagy induction.

ChIP–seq analysis in TFEB-FLAG HeLa cells identified 46 TFEB-bound genes among 623 annotated endocytic genes, while transcriptomic profiling of TFEB-GFP–expressing HeLa cells revealed 35 endocytic genes upregulated relative to wild-type controls (Nnah et al. 2019). These datasets defined a subset of putative TFEB-regulated components of the endocytic pathway, including MCOLN1. Independent studies have further suggested TMEM55B as a potential transcriptional target of TFEB and TFE3 (Sardiello et al. 2009{Pastore, 2016 #194; Martina et al. 2016; Pastore et al. 2016). To extend these findings, we next asked whether concurrent perturbation of TFEB-regulated lysosome-associated genes might exert additive or synergistic effects on neurodegenerative phenotypes. To address this, we genetically downregulated both genes and systematically assessed their combined impact using multiple established assays of neurodegeneration. This approach allowed us to evaluate whether coordinated disruption of TFEB-dependent lysosomal pathways exacerbates neuronal dysfunction beyond the effects observed upon individual gene perturbation.

In order to verify whether loss of dTMEM and lysosomal transient receptor protein cation channel, trpml leads to enhanced locomotory defects, lethality, decline in survival of the animals and cell death we performed various fly locomotory, lethality, survival assays, TUNEL and cleaved caspase 3 immunostainings in the fly brain. Our results indicated that loss of dTMEM and trpml, leads to enhanced locomotory defects in 21 days old flies [Figure 10A, Supplementary Figure 12Aiii], enhanced lethality [Figure 10B] and reduced lifespan [Figure 10C] in an age dependent manner. Scatter plots were employed to represent individual climbing assay measurements, as neurobehavioral assays often exhibit substantial inter-individual variability. Displaying individual data points enables assessment of data distribution, variability, and outliers, thereby strengthening the interpretation of genotype- and age-dependent behavioral effects. Loss of dTMEM55 and trpml, also leads to enhanced death assayed by cleaved caspase 3 [Supplementary Figure 13, A-E] and TUNEL staining [Supplementary Figure 13, F-G] in 21 – 28 days old fly brains. These results hint towards a possible genetic interaction between the two TFEB regulated genes, MCOLN1 and TMEM55B, in enhancing neuronal degeneration in AD.

**Figure 10:**
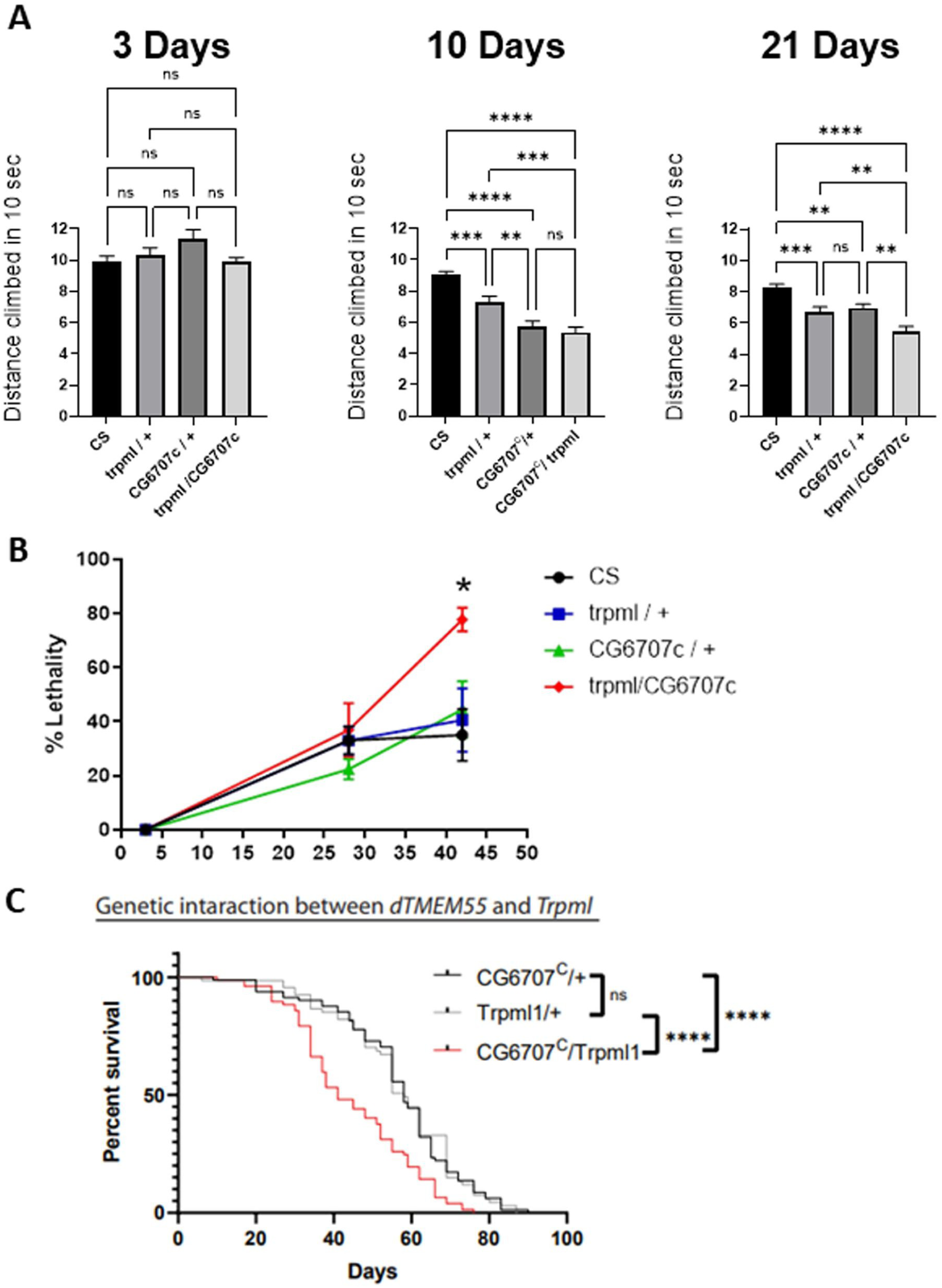
Loss of dTMEM and trpml leads to enhanced locomotory defects, lethality and decline in survival of the animals. (A) Bar graphs representing locomotory function of adult flies with a measurement of distance climbed by adult male flies (3 days, 10 days old and 21 days) in 10 seconds compared between various genotypes, N > 60. (B) Line graphs depicting quantitative extent of lethality in flies of various genotypes listed below, N = 8 tubes [each tube = 10-12 flies]. (C) Aging curve depicting the lifespan of flies of various genotypes, N = 5 tubes [each tube = 20 flies]. Genotypes: CS: wild type CantonS; trpml / +: trpml heterozygous mutant; CG6707c / + = heterozygous dTMEM mutant; trpml / CG6707c = trpml and dTMEM double mutants. p < 0.05; **: p < 0.01; ***: p < 0.001; n.s: p ≥ 0.05. p-value was calculated using Prism 7’s two tailed Unpaired t-test and One way Anova, multiple comparison.

## Discussion

Autophagosomal–lysosomal dysfunction is an early feature of Alzheimer’s disease (AD), preceding amyloid-β and Tau pathology (Nixon and Yang 2011). AD brains show enlarged APP-positive endosomes (Takahashi et al. 2002), lysosomal deficits, and autophagic vacuole accumulation (Cataldo et al. 1997), (Nixon and Yang 2011), (Nixon et al. 2005). Cathepsin polymorphisms increase AD risk (Bhojak et al. 2001), (Papassotiropoulos et al. 1999), whereas cystatin B/C deletion improves phenotypes (Mi et al. 2007), (Yang et al. 2011), (Yang et al. 2014). Neurodegeneration is linked to impaired autophagy and trafficking (Dobrowolski and De Robertis 2011), (Sorkin and von Zastrow 2009), (Taelman et al. 2010), (Killisch et al. 1991), (Lipinski et al. 2010), (Nixon 2013). mTORC1–TFEB signaling governs lysosomal biogenesis (Laplante and Sabatini 2012), (Inoki et al. 2003), (Tee et al. 2003), (Sardiello et al. 2009), (Settembre et al. 2012), (Settembre et al. 2013), (Sardiello et al. 2009), (Martina et al. 2014), and TFEB targets include several genes, including MCOLN1 (Nnah et al. 2019).

TMEM55B was long considered a lysosomal protein of unknown function, though it regulates nuclear phosphatidylinositol-5-monophosphate (Ungewickell et al. 2005), (Zou et al. 2007). Puertollano and coworkers (Willett et al. 2017) showed TMEM55B recruits JIP4 to lysosomes, promoting dynein-dependent retrograde transport, and depletion causes peripheral lysosome accumulation without affecting endosomes or Golgi. TMEM55B is a TFEB/TFE3 target induced by starvation and cholesterol stress (Willett et al. 2017). S-palmitoylation regulates its trafficking (Rudnik et al. 2022), while ERK/MAPK-mediated phosphorylation controls perinuclear clustering (Takemasu et al. 2019). TMEM55B mediates PLEKHM1 degradation, ESCRT repair, and TFE3 activation (Jeong et al. 2024). Knockout zebrafish show oxidative-stress sensitivity. TMEM55B associates with V-ATPase/Ragulator to regulate mTORC1 (Hashimoto et al. 2018) and controls PI(4,5)P2-dependent LDLR and cholesterol metabolism (Qin et al. 2020).

Even though a few groups have ventured in deciphering some putative roles of this molecule in autophagy, a clear-cut mechanistic underpinning in its role in regulating Tau mediated neurodegeneration using invitro, invivo, patient derived clinical datasets and iPSC derived differentiated neurons have not been previously shown. In this study, we report the critical role of the lysosomal transmembrane protein TMEM55B [TMEM55] in mediating neurodegeneration in transgenic in vivo fly and *invitro* cell culture models of Alzheimer’s disease.

Integrated analysis of clinical, microarray and bulk RNA-sequencing datasets from the ROSMAP cohort, derived from dorsolateral prefrontal cortex, revealed that *TMEM55B* transcript levels were significantly reduced exclusively in severe AD patients. Expression of housekeeping genes (*RPL13A*, *GAPDH*) remained stable across control and AD groups, confirming that the observed changes were not due to global transcriptional decline. In the same cohort, severe AD samples exhibited downregulation of the lysosomal gene *MCOLN1* and concomitant upregulation of the AD-associated gene *MAPT*, consistent with progressive Tau pathology. These findings indicate stage-specific dysregulation of *TMEM55B*, absent in mild or moderate AD. Because bulk ROSMAP data represent mixed cortical cell populations, we next examined neuronal-specific expression using single-nucleus RNA-sequencing from the ROSMAP consortium. Analysis of neuronal nuclei from dorsolateral prefrontal cortex demonstrated significantly lower *TMEM55B* levels in AD compared with controls. Collectively, multiple human post-mortem NGS datasets suggest a potential association between neuronal *TMEM55B* downregulation and AD-related neurodegeneration. Our findings indicate that the observed phenotypes are restricted to late-stage Alzheimer’s disease (AD) cases, consistent with the clinical reality that early-stage AD is largely asymptomatic and lacks overt clinical manifestations. While molecular alterations are known to initiate during the early phases of the disease, these changes often remain subtle and do not reach statistical or pathological prominence until later stages. In line with this framework, we observed that the molecular changes described here become robust and statistically significant only in severe AD. Importantly, although the most pronounced alterations were detected in severe AD cases, these results remain highly impactful, as they position TMEM55B as a potential biomarker and disease-associated factor that precedes or accompanies progression to severe AD. TMEM55B dysregulation was specifically associated with severe AD and was not observed in mild or moderate cases, suggesting a stage-dependent relationship with disease severity.

Human TMEM55B gene shares 99% homology with *Drosophila* dTMEM55. Therefore, we decided to create a fly transgenic model to understand how TMEM55B downregulation can lead to AD. We generated various transgenic dTMEM55 loss of function flies to evaluate how downregulation of TMEM55B affects neurodegeneration in AD and understand the mechanisms underlying abnormal Tau accumulation. Complex brain functions like memory, circadian rhythms, and movement are affected in patients with neurodegenerative diseases. In *Drosophila*, these complex brain functions are impaired along with degeneration of neuronal structures. The climbing assay is one of the simplest and routinely used method for measuring the motor function of flies. Previous study of Roy et al and others (Roy and Jackson 2014), (Nitta and Sugie 2022), have used locomotory assays as a means to evaluate neurodegeneration. In our current study we have used similar paradigms like climbing assays to evaluate how downregulation of dTMEM55 affects age dependent locomotory defects in flies. Both dTMEM55 mutant flies and flies with RNAi mediated TMEM55B downregulation using a motor neuron and pan-neuronal driver showed locomotory defects in 10 days old flies. However, the 3-day old flies in these transgenics, did not show any defects in these climbing assays. In a similar fashion, the dTMEM55 knockout flies also displayed climbing defects in an age dependent manner. Since behavioural output like locomotory or climbing assays are reliable readouts for neuronal dysfunction, all these results lead to the conclusion that downregulation of TMEM55B results in neuronal dysfunction in an age dependent manner. In order to assess whether neuronal dysfunction phenotype is specifically associated with the loss of function of dTMEM55 and not any other gene we overexpressed dTMEM55 and human TMEM55B in the cells where dTMEM55 levels were specifically downregulated. We were able to rescue the climbing defects in 10-day old flies, indicating the vital and specific role of dTMEM55 in maintaining optimum neuronal function and output. Additionally, rescue by the human TMEM55B indicates that the human TMEM55B shares similar functional attributes as of dTMEM55. These results strongly support that information obtained through the neuronal dysfunction phenotype from fly TMEM55B loss of function models and can be extended to neuronal degeneration in human AD. Lifespan or lethality assays in flies is another common read out for testing neuronal toxicity in evaluating any gene’s function in neurodegenerative disease (Nitta and Sugie 2022). Using RNAi mediated TMEM55B downregulation using motor neuron driver lethality assays were performed. These results indicate that dTMEM55 downregulation enhances lethality in 43 days old flies. Lethality assay is one of the quantitative approaches to evaluate neurotoxicity and is widely used to assess the effect of various genes in modulating toxicity in various neurodegenerative diseases (Nitta and Sugie 2022). Therefore, these results conclude that TMEM55B potentiates neurodegeneration in AD by enhancing neuronal toxicity.

We additionally validated that all our studies in the various fly TMEM55B models, are due to altered TMEM55B levels by using RT-PCR to probe for TMEM55B RNA levels. We found that the TMEM55B levels were lowered in dTMEM55 loss of function flies and elevated in the dTMEM55 gain of function flies.

Lysosomes are important for maintenance of protein and organelle homeostasis in cells. A lysosome’s optimum functioning is important for neuronal longevity and health. Maintenance of optimum axonal lysosomal transport is important for regulating neuronal development, its maintenance and functioning. It is also known that perturbations to optimal axonal lysosomes, or abnormal accumulation of decreased number of axonal lysosomes, are linked to altered neuronal function and health. Any change in optimal axonal lysosome number has been known to lead to several neurological diseases including neurodegenerative diseases (Paumier and Gowrishankar 2024). Investigations in both human AD brain tissue and mouse models of AD have shown the excessive accumulation of lysosomes in swollen axons near extracellular amyloid aggregates, which formed part of the neuritic plaque (Terry et al. 1964), one of the major neuropathological hallmarks of AD. These multilamellar and multivesicular vesicles were observed by electron microscopy (Terry et al. 1964) and resembled axonal organelles that build up due to retrograde transport defects or blockage (Tsukita and Ishikawa 1980). Abnormal lysosome trafficking and signaling has been implicated in various diseases, including neurodegenerative diseases and various molecular players have been shown to have an important role in regulating these processes (Lie and Nixon 2019). In our studies we found that downregulation of dTMEM55 resulted in abnormal accumulation of lysosomes in the larval axons in comparison to the control larval axons. These results indicate that lowered TMEM55B could lead to neurodegeneration by affecting normal axonal transport of lysosomes.

Age dependent neurodegeneration has been observed due to expression of β-amyloid (Aβ) or *Tau* encoded by the gene microtubule-associated protein tau (MAPT), both of which are associated with the prominent AD pathologies, in *Drosophila* brain (Iijima et al. 2004), (Finelli et al. 2004), (Crowther et al. 2005), (Wittmann et al. 2001). AD is an age dependent neurological disorder that causes irreversible loss of neurons, especially in the brain cortex and hippocampus. In the brains of AD patients, senile Abeta plaques of Abeta aggregates and neurofibrillary tangles (NFTs), comprising of aggregates of highly phosphorylated Tau protein, are observed. These are the major neuropathological hallmarks of AD (Scheltens et al. 2021). *Drosophila* is routinely used as a simple, genetically tractable in *vivo* model for studying the molecular pathogenesis of AD (Bouleau and Tricoire 2015), (Jeon et al. 2020).

Among the various *Drosophila* models, flies that express Aβ have been widely used to study AD disease models. Although there is no protein homologous to Aβ in *Drosophila*, flies expressing human Aβ in all nervous systems have been shown to have impaired learning, accumulation of amyloid plaques, neurodegeneration, and shortened lifespan. All these neuronal generation phenotypes in flies mimicked clinical symptoms observed in patients with AD (Iijima et al. 2004), (Finelli et al. 2004), (Crowther et al. 2005).

The expression of Tau using pan-neuronal Elav-Gal4 driver led to reduced lifespan in flies with an increase in the number of vacuoles in the cortex and neuropile of flies in an age dependent manner (Wittmann et al. 2001). The R406W mutation in the MAPT gene, responsible for the cause of frontotemporal dementia with Parkinsonism linked to chromosome 17 tau with AD-like clinical features (Reed et al. 1997), (van Swieten et al. 1999), (Saito et al. 2002), resulted in more severe Tau-associated pathology (Wittmann et al. 2001). Additionally, the expression of Tau in cholinergic neurons by Choline acetyltransferase-Gal4 (Cha-Gal4) (Yasuyama and Salvaterra 1999) significantly reduced the number of cholinergic neurones after 60 days (Wittmann et al. 2001). Cholinergic neurons have been shown to be particularly vulnerable to neurodegeneration in AD and can also be affected in other tauopathies (Geddes et al. 1993). One of the simple methods for evaluating neuronal toxicity of expressed human genes and alleles is the rough eye phenotype in flies. The *Drosophila* eye consists of around 700–800 basic structural units called ommatidia, each consisting of 8 photoreceptor neurones (primary sensory neurons), 4 cone cells, and 2 primary pigment cells (Kumar 2012). Even though these ommatidia are generally regularly aligned, they may become disorganized if ectopic genes induce morphological differentiation, proliferation, or cell death in ommatidial cells, thereby causing morphological abnormalities of the compound eye, leading to the formation of the rough eye. Gene toxicity is evaluated via a qualitative assessment of the structural deformities of the eye. The structural abnormality of the eye, if induced by a human gene, can be utilized for conducting a modifier screen. Such a screen can be easily performed using the extent of rescue of the structural abnormality as an indicator. This method does not require any specific high-priced equipment other than epi-microscopy and therefore has the potential to efficiently evaluate neuronal toxicity. Previously, Roy et al. (Roy and Jackson 2014), have used the rough eye phenotype as a readout to evaluate the interaction between Tau and α-Syn in augmenting neurotoxicity in a fly model of Parkinson’s Disease. It was shown that misexpression of Tau and α-Syn enhanced a rough eye phenotype in the *Drosophila* Tauopathy model. Misexpression of both Tau and α-Syn resulted in a more severe rough eye, than when Tau is mis expressed alone, indicating that α-Syn enhances the Tau mediated rough eye phenotype, retinal degeneration and neurotoxicity. As a step forward, we wanted to evaluate whether downregulation of dTMEM55 modulates Tau toxicity. We downregulated dTMEM55 expression using a mutant dTMEM55 fly [CG6707c/+] crossing it to a fly overexpressing human wild-type full-length (2N/4R) tau using a *gl*-Tau construct (human tau cDNA fused directly to the eye-specific *gl* promoter (Ambegaokar and Jackson 2011). *gl*-Tau misexpression causes a rough phenotype of the external eye associated with abnormalities of photoreceptor neurons and other cell types in the underlying retina (Jackson et al. 2002). In the *gl*-Tau flies, the degree of the rough eye phenotype is intermediate and more pronounced anteriorly. These flies have reduced eyes, with the posterior region of the eye remaining more normal. Misexpression of both human Tau with a concomitant downregulation of dTMEM55 resulted in a more severe rough eye, with the abnormal phenotype extending farther posteriorly. We also assayed the extent of rough eye phenotype in the *gl*-Tau flies, with the neuronal knockdown of dTMEM55. Additionally, we have shown that Tau mediated roughness increased significantly in eyes of flies with specific dTMEM55 downregulation in the neurons (using pan-neuronal Elav Gal4 driver and UAS dTMEM55 RNAi), in comparison to when Tau was mis-expressed alone. We also observed that downregulation of dTMEM55 enhances eye roughness in flies mis-expressing TauP301L [a Tau mutation most frequently observed in patients with frontotemporal dementia with parkinsonism linked to chromosome 17 (FTDP-17), in comparison to flies expressing TauP301L alone, without any dTMEM55 downregulation. The specific role of dTMEM55’s downregulation in enhancing Tau mediated rough eye phenotype or retinal degeneration was shown by expressing a Duplication transgene of dTMEM55 [CGDup], which had the potential to restore the dTMEM55 to optimum levels, against the loss of function dTMEM55 genetic background. In these flies, the rough eye phenotype was significantly reduced in comparison to flies where Tau was misexpressed against a dTMEM55 loss of function background. All these results concluded, that TMEM55B had the potential to enhance both wild type Tau and mutant Tau P301L phenotypes in a fly AD model, an indicator of its plausible role in potentiating Tau mediated neurodegeneration in Alzheimer’s Disease. Motor behaviour has been used to probe for neuronal function in various neurodegenerative models. Tau has previously been shown to be involved in axonal transport defects and locomotor dysfunction in a *Drosophila* model (Mudher et al. 2004). To examine whether downregulation of dTMEM55 modulates Tau mediated adult locomotor behaviour, we used the motor neuronal *OK371*-GAL4 driver to express Tau in postmitotic neurons. We tested motor behaviour using a simple climbing assay (negative geotaxis) in which a single adult male (3,10 and 21 days post-eclosion) is tapped to the bottom and allowed to climb up an empty vial for 10s. The distance travelled by the individual fly within the given time frame is a measure of its motor performance or climbing index. We observed that 21 days old flies expressing Tau alone or flies with the loss of function dTMEM55 genetic background had severe motor deficits, with respect to the controls. However, 21 days old flies, misexpressing wild type Tau against a loss of function dTMEM55 genetic background showed significantly higher motor behaviour defects than age matched flies, expressing Tau alone or flies having only dTMEM55 downregulated. To assess whether downregulation of dTMEM55 enhances Tau mediated neuronal toxicity, we used fly lethality assay. Mis-expression of Tau alone using the motor neuronal driver enhanced lethality in 28 days old flies. However, this lethality was further exacerbated when Tau was mis-expressed against a dTMEM55 loss of function genetic background. This result concludes that downregulation of dTMEM55 enhances Tau mediated neuronal toxicity.

The microtubule-binding protein tau is a component of neurofibrillary tangles (NFT) in Alzheimer’s disease (AD) and related disorders that are collectively referred to as tauopathies (Morishima-Kawashima et al. 1995), (Johnson and Bailey 2002). In the NFT, Tau is hyperphosphorylated and it is widely believed that tau kinases such as glycogen synthase kinase-3β (GSK)-3β, microtubule affinity-regulating kinase (MARK) and cyclin-dependent kinase 5 (Cdk5) plays a role in the formation of insoluble tau aggregates and NFT (Cho and Johnson 2003), (Augustinack et al. 2002). The phosphorylated Tau forms the NFT which is one of the neuropathological hallmarks in AD. To assess whether downregulation of dTMEM55 leads to any change in the p-Tau (pSer202/pThr205) levels in our model of TMEM55B’s role in AD, we performed immunoblots with head extracts. We probed the blot with AT8, a monoclonal antibody that recognizes Tau phosphorylated at serine 202 and threonine 205, T-46, which detects total Tau, and GAPDH as a loading control. We detected an increase in AT8 signal in flies mis-expressing Tau along with downregulation dTMEM55 in comparison to flies only mis-expressing Tau alone without any dTMEM55 downregulation. This kind of elevated p-Tau levels was observed in flies with downregulated dTMEM55 with either wild type Tau [Tau 1.1] or mutant Tau [P301L Tau] mis-expression [Figure 6Aii, 6Bii]. This indicates that loss of function dTMEM55 elevates phosphorylated Tau levels, a clinical-pathological manifestation known to be associated with neurofibrillary tangles in AD. Thus, downregulation of TMEM55b could be potentially leading to elevated p-Tau, whose excessive accumulation could lead to formation of NFTs, leading to neuronal degeneration in AD. All these assays, including retinal degeneration, climbing defects and lethality showed that the downregulation of dTMEM55 enhanced Tau mediated neuronal degeneration, dysfunction and toxicity in the fly transgenic AD models, indicating that these neuronal defects might lead to neurodegeneration in AD patients. We observed that the loss of function of dTMEM55 [*Drosophila* TMEM] leads to locomotion defects, enhances lethality, accumulation of lysosomes in the axonal nerves, enhances Tau-mediated neurodegeneration. Biochemical analysis revealed that downregulation of dTMEM leads to elevated levels of Tau in fly brains.

To establish the translational relevance of our fly findings, we tested whether the TMEM55B loss-of-function phenotypes observed in *Drosophila* dTMEM55B models could be effectively recapitulated in a human system using iPSC-derived neurons. We firstly characterized the iPSC derived neurons by western blot with the help of markers specific for iPSCs and differentiated neurons. Induced pluripotent stem cells (iPSCs) are generated through the reprogramming of somatic cells to a pluripotent state (Shi et al. 2017). These cells possess the capacity to differentiate into multiple specialized lineages and represent a powerful platform for modelling human development, degenerative disorders, and therapeutic interventions (Marchetto et al. 2011). Several iPSC-derived models of human forebrain neurons have been used in understanding mechanisms underlying several brain related diseases, as these cells are known to mediate higher order brain functions, including consciousness (Baxter and Chiba 1999), emotion (Morgane et al. 2005), and sleep (Schwartz and Roth 2008). As a result, malfunctions in these cells can cause a wide range of neurological disorders, including neurodegenerative diseases like Alzheimer’s (Auld et al. 2002) and Huntington’s (McColgan and Tabrizi 2018), as well as neurodevelopmental diseases such as autism (Donovan and Basson 2017) and epilepsy (Heath 1976).

For our studies, two iPSC cell lines were induced, one being the wild type, representing cell lines from healthy patients and other being the PSEN1-M146L cell line {a mutation in the PSEN1 gene (Mangone et al. 1995), (Morelli et al. 1998), known to cause familial AD}, representing AD patients’ cell line. We performed Western blot on the iPSCs and the differentiated neurons of these cell lines, using antibody against OCT4 [octamer-binding transcription factor 4], a primary cell marker used to identify iPSCs. OCT4 is known as the key transcription factor crucial for maintaining pluripotency in stem cells; and its expression is known to signify the undifferentiated state of an iPSC. The iPSCs showed high levels of OCT4, while the 20-days and 40-days old neurons lacked OCT4 signal. This indicated that the 20-days and 40-days old neurons were mature differentiated neurons and had lost the stem cell potency. Additionally, we also tested iPSCs, 20-days and 40-days old neurons with antibody against various mature differentiated neurons like Tuj-1 [class III beta-tubulin, marker of neurons in the central and peripheral nervous systems from the early stage of neural differentiation], Tau [MAPT, microtubule associated protein Tau, a protein found in neurons that helps maintain the stability of microtubules] and Synapsin [Synapsin is considered a reliable neuronal marker because it is a protein exclusively expressed in neurons, specifically located at the presynaptic terminal, and is directly associated with synaptic vesicles, making it a strong indicator of the presence of synapses within a tissue sample]. All the three neuronal markers were elevated in the mature differentiated neurons [both 20-days and 40-days] and were missing from the iPSCs. All these results indicated that the protocol used to induce iPSCs to neuronal stem cells, then induce the neuronal stem cells to neuronal progenitor cells and then differentiate NPCs to adult mature neurons yielded a high percentage of mature neurons. Additionally, these neurons tested positive for all neuronal markers [like Tuj-1, Tau and Synapsin] and negative for iPSC marker like OCT4. Then we proceeded to test the levels of TMEM55B in mature differentiated Control and AD neurons. We labelled the differentiated neurons, with an antibody against MAP2, [Microtubule-associated protein 2, a protein known to aid in the organization and regulation of the neuronal cytoskeleton], and counterstained it with antibody against TMEM55B.The MAP2 antibody used targets MAP2c as well as MAP2b. MAP2c is a low molecular-weight form of Microtubule-Associated Protein 2 (MAP2) found abundantly in neuron axons. MAP2c is a key MAP found in axons, often along with tau. It helps bundle microtubules (Higgs and Das 2022), influences their spacing (similar to tau), and interacts with actin, playing roles in axonal development and function by regulating microtubule organization and potentially signalling pathways (Roger et al. 2004). MAP2c promotes microtubule stabilization and bundling, interacts directly with filamentous actin, and coordinates actin–microtubule cross-talk, thereby regulating cytoskeletal organization, neurite initiation, and axonal growth (Pacheco and Gallo 2016). DAPI was used to stain the nucleus of these neurons. We found that there was no change in the TMEM55B levels between 20 days old control and AD neurons. However, we found that TMEM55B levels are significantly lowered in 40 days old AD neurons in comparison to 40 days old control neurons. These results indicated that TMEM55B is lowered in aged AD neurons and not the early AD neurons. In the next step, we wanted to test whether lowering of TMEM55B in the aged AD neurons leads to any changes in the phosphorylated form of Tau in the lysosomes of these neurons. The formation of phosphorylated form of Tau and its accumulation is associated with the formation of neuropathological hallmarks (NFTs) in the AD brains. Therefore, an accumulation of p-Tau in the lysosomes upon downregulation of TMEM55B will hint towards some kind of defects in lysosomal mediated autophagic clearance of p-Tau, leading to neuronal degeneration by abnormal buildup of pathological forms of Tau. We observed that there was an increase in p-Tau levels in the lysosomes located in the soma of the 40 days old iPSC derived AD neurons in comparison to p-Tau levels in the soma of 40 days old iPSC derived Control neurons. These results in the iPSC derived neurons, hints towards a possible defect in lysosome mediated autophagic clearance of p-Tau in the aged and severe AD patients, leading to neurodegeneration. Additionally, quantitative analysis of 40-day-old iPSC-derived AD neurons revealed a significant increase in the total number of axonal lysosomes compared with age-matched control neurons. Notably, axonal lysosomes positive for phosphorylated Tau were also markedly increased. These observations suggest defects in axonal lysosomal transport in AD neurons, potentially impairing autophagic degradation. Importantly, this phenotype recapitulates findings from transgenic *Drosophila* model, where downregulation of dTMEM55 led to abnormal lysosomal accumulation in larval axons compared with controls. Together, these data indicate that impaired lysosomal transport may contribute to defective autophagic flux, leading to defective clearance of protein aggregates and organelles, thereby contributing to progressive neuronal dysfunction and degeneration observed in aged and severe Alzheimer’s disease patients.

Autophagic flux, encompassing autophagosome formation, maturation, lysosomal fusion, and degradation, is essential for neuronal homeostasis, and its dysfunction contributes to AD, PD, HD, and ALS (Williams et al. 2006), (Rubinsztein et al. 2007), (Wong and Cuervo 2010). Defects in induction, maturation, or lysosomal clearance impair protein degradation (Zhong et al. 2009), (Pickford et al. 2008), (Ravikumar et al. 2005), (Webb et al. 2004), (Iwata et al. 2005), (Kimura et al. 2008), (Wong and Cuervo 2010), (Lee et al. 2010), and is assessable via LC3-II turnover or p62 degradation (Mizushima et al. 2010), (Ju et al. 2010), (Klionsky et al. 2008b), (Tanida et al. 2006), (Bjorkoy et al. 2005), (Mizushima and Yoshimori 2007). We chose the most straightforward method for monitoring autophagy, which is the detection of LC3 protein processing. LC3 proteins are specifically cleaved at the C-terminus by Atg4 to become LC3-I, which then conjugates to phosphatidylethanolamine to form LC3-II. Based on the observation that LC3-II is degraded in autolysosomes, the level of LC3-II or GFP-LC3-II is widely used as a marker for monitoring the autophagic process (Klionsky et al. 2012). However, an increased level of LC3-II or an accumulation of GFP-LC3 puncta is not always indicative of autophagy induction and may represent a blockade in autophagosome maturation (Fass et al. 2006). Autophagic flux can be detected by LC3-II turnover using Western blot analysis in the presence and absence of lysosomal degradation inhibitors, such as pepstatin A, E64d, bafilomycin A1, chloroquine and NH4Cl (Ju et al. 2010), (Klionsky et al. 2008b). If autophagic flux is occurring, the level of LC3-II will be increased in the presence of a lysosomal degradation inhibitor because the transit of LC3-II through the autophagic pathway will be blocked.

Autophagic flux is a dynamic, multistep process involving autophagosome biogenesis, maturation, fusion with lysosomes, and degradation of sequestered cargo. Disruption of autophagic flux, particularly at late stages, is increasingly recognized as a central pathological feature in neurodegenerative disorders. Chloroquine (CQ) has long been used as a pharmacological inhibitor of autophagy, although its precise mechanism of action remained incompletely understood until recent studies provided detailed mechanistic insights. Additionally, Dobrowolski et al. demonstrated that lysosomal inhibition by chloroquine enhances canonical Wnt signaling by promoting accumulation of multivesicular endosomes (MVEs), leading to sequestration of GSK3 within late endosomes. This mechanism stabilizes β-catenin and other GSK3 substrates, revealing a key regulatory role for the endolysosomal pathway in Wnt signal transduction (Dobrowolski et al. 2012). Two landmark investigations, Mauthe et al., 2018 and Nnah et al., 2019, demonstrated that CQ primarily blocks autophagic flux by impairing autophagosome–lysosome fusion through disruption of endo-lysosomal trafficking pathways and vesicular fusion machinery (Mauthe et al. 2018), (Kodani et al. 2019). Mauthe et al. (2018) performed a comprehensive mechanistic analysis using biochemical, ultrastructural, and live-cell imaging approaches to define how CQ inhibits autophagy. Contrary to the classical model that attributed CQ-mediated flux inhibition to lysosomal alkalinization and impaired hydrolase activity, the authors demonstrated that CQ predominantly disrupts autophagosome–lysosome fusion. CQ treatment led to a pronounced accumulation of LC3-II–positive autophagosomes and elevated levels of the autophagy adaptor p62/SQSTM1, reflecting defective cargo degradation. Notably, lysosomal acidification was only modestly affected, indicating that CQ does not primarily act by neutralizing lysosomal pH. Instead, CQ impaired the recruitment and assembly of the SNARE fusion complex composed of syntaxin 17 (STX17), SNAP29, and VAMP8, which is essential for autophagosome–lysosome membrane fusion. Furthermore, CQ altered endo-lysosomal trafficking dynamics, resulting in enlarged, dysfunctional endosomal and lysosomal compartments, thereby preventing efficient autophagosome docking and fusion. These findings redefined CQ as a potent inhibitor of vesicular trafficking and membrane fusion rather than solely a lysosomotropic alkalinizing agent (Mauthe et al. 2018). Extending these mechanistic insights, Nnah et al. (2019) conducted a systematic functional screen of 623 endocytic genes to identify regulators of autophagic flux. This large-scale analysis uncovered 46 endocytic components that critically modulate autophagy, many of which phenocopied CQ-induced flux blockade when genetically suppressed. Loss of these genes resulted in robust accumulation of autophagosomes, increased LC3-II and p62 levels, and impaired degradation of long-lived proteins, indicating severe defects in late-stage autophagy. Mechanistically, disruption of endocytic trafficking impaired lysosomal maturation, positioning, and functional competence, thereby preventing effective autophagosome–lysosome fusion. Importantly, several of these endocytic regulators were transcriptionally controlled by transcription factor EB (TFEB), highlighting a coordinated regulatory axis linking endocytosis, lysosomal biogenesis, and autophagic clearance. Pharmacological treatment with CQ further exacerbated these genetic defects, revealing convergence between chemical inhibition and genetic perturbation of endocytic pathways (Kodani et al. 2019).

Collectively, these studies establish a unified model in which chloroquine inhibits autophagic flux primarily by disrupting endo-lysosomal trafficking and SNARE-dependent autophagosome–lysosome fusion. This mechanistic framework has significant implications for neurodegenerative diseases, where defects in endocytosis, lysosomal function, and autophagy contribute to progressive neuronal dysfunction. The insights provided by these studies not only refine the interpretation of CQ-based autophagy assays but also highlight endocytic trafficking pathways as promising therapeutic targets for restoring autophagic flux in neurodegenerative disorders.

We evaluated the extent of autophagic flux in iPSCs and iPSC derived neurons in the presence and absence of lysosomal degradation inhibitor, chloroquine by evaluating LC-II and LC-I levels. We observed an increased LC-II levels in the presence of chloroquine in the iPSC derived 20 and 40-day-old AD forebrain neurons. However, no change in the levels of LC-II levels was observed in the presence of chloroquine in the iPSCs. We observed a robust, statistically significant decline in autophagic flux in the 20 day and 40-day-old iPSC derived AD neurons in comparison to the age matched iPSC derived Control neurons. There was no autophagic flux defects in the AD iPSCs and the levels of autophagic flux in the AD iPSCs were comparable to the Control iPSCs. Our results indicates that autophagic flux defects are prominent in AD neurons. These results additionally establish that these neurons could be potentially used for studying the effects of various lysosomal proteins in mediating autophagic flux to enhance clearance of Tau and other proteins known to form toxic aggregates in neurodegenerative diseases. The observation that autophagic flux defects are evident in 20-day-old neurons, whereas reductions in TMEM55B levels are detected only in 40-day-old AD neurons, suggests a temporal separation between the onset of autophagy impairment and TMEM55B downregulation. This pattern raises the possibility that early defects in autophagic flux may involve additional regulatory factors, with altered TMEM55B expression potentially contributing at later stages. However, further studies will be required to determine whether TMEM55B plays a direct or indirect role in modulating autophagic flux during disease progression.

Abnormal lysosomal positioning is a hallmark of Alzheimer’s disease (AD), with lysosomes accumulating in dystrophic axons around amyloid plaques (Terry et al. 1964; Cataldo et al. 1994). Mouse studies indicate this arises from impaired retrograde axonal transport and lysosomal maturation, rather than increased anterograde trafficking, which enhances amyloidogenic APP processing by limiting lysosomal degradation (Gowrishankar et al. 2015; Gowrishankar et al. 2017).

Lysosomal positioning is determined by the balance of the bidirectional lysosomal movement, changes under various conditions and affects lysosomal functions such as autophagy. Evidence have directed to the fact that the distribution of lysosomes within cells is regulated in response to a variety of stimuli, and alterations in this regulation is associated with various pathologies. For example, under starvation conditions, autophagosomes and lysosomes move toward the cell centre, facilitating the fusion between both organelles and degradation of the autophagosomal content (Kimura et al. 2008), (Korolchuk et al. 2011). Several protein complexes have been implicated in the regulation of lysosomal positioning. Anterograde movement is mediated by BORC, Arl8, SKIP, and Rab7–FYCO1 complexes (Rosa-Ferreira and Munro 2011), (Pu et al. 2015), (Matsuzaki et al. 2011), (Raiborg et al. 2016), while retrograde transport relies on Rab7, RILP, LAMP-1/2, and ALG-2/MCOLN1 interactions recruiting dynein–dynactin (Cantalupo et al. 2001), (Johansson et al. 2007), (Huynh et al. 2007), (Krzewski et al. 2013), (Li et al. 2016), (Vergarajauregui et al. 2009). All these findings suggested that lysosomal transmembrane proteins regulate retrograde lysosomal transport. Additionally, overexpression of Rab34, Rab36, Rabring7, or Rapsyn causes the clustering of lysosomes in the perinuclear area (Chen et al. 2010), (Wang and Hong 2002), (Mizuno et al. 2003), (Aittaleb et al. 2015). Our results showed that the knockdown of TMEM55B led to lysosomal positioning defects in HEK293T cells and iPSC-derived AD forebrain neurons. Collectively, our data suggest that TMEM55B may act as a conserved regulator of lysosomal positioning, with its downregulation potentially contributing to Tau-associated neurodegenerative processes in AD. TMEM55B deficiency appears to disrupt retrograde lysosomal transport, which could lead to axonal lysosome accumulation, impaired autophagic clearance, and Tau accumulation. These observations support a possible mechanistic link between TMEM55B dysfunction and neuronal vulnerability, highlighting TMEM55B as a potential candidate mediator of lysosome-dependent proteostasis in susceptible neuronal populations.

Ballabio, Medina, and colleagues demonstrated that lysosomal Ca²⁺ signaling plays a critical role in regulating the phosphatase calcineurin and its substrate TFEB, a master transcription factor governing lysosomal biogenesis and autophagy (Medina and Ballabio 2015), (Medina et al. 2015). Lysosomal Ca²⁺ release through mucolipin 1 (MCOLN1) activates calcineurin, which dephosphorylates TFEB, promoting its nuclear translocation and transcriptional activation of lysosomal and autophagic genes. Genetic or pharmacological inhibition of calcineurin suppresses TFEB activity under conditions such as starvation and physical exercise, whereas calcineurin overexpression or constitutive activation enhances TFEB function. Importantly, induction of autophagy and lysosomal biogenesis via TFEB requires MCOLN1-mediated calcineurin activation, establishing a mechanistic link between lysosomal calcium signaling, calcineurin regulation, and autophagy induction. In parallel, our functional studies in *Drosophila* models revealed that TMEM55-mediated neurodegenerative phenotypes were further exacerbated in MCOLN1 loss-of-function flies, indicating that TMEM55 and MCOLN1 act additively to modulate lysosomal positioning, transport, and neuronal homeostasis. These findings suggest that TMEM55 and MCOLN1 cooperate to maintain lysosome-dependent proteostasis, and that disruptions in either pathway can enhance neuronal vulnerability. Taken together, these observations support a model in which TMEM55 and MCOLN1 might jointly regulate lysosomal positioning, retrograde transport, and calcium-dependent signaling to ensure efficient autophagic flux and clearance of pathological proteins. Dysregulation of this network may possibly contribute to impaired autophagic clearance, Tau accumulation, and progressive neuronal degeneration, providing mechanistic insight into lysosome-dependent pathways that could underlie neurodegenerative processes in Alzheimer’s disease and related disorders.

Overall, our data suggests that the downregulation of TMEM55 can potentially cause neurodegeneration in AD cases, due to lysosome transport defects [Schematic model, Figure 11A, 11B], preventing the autophagic clearance of Tau. In summary, our study identifies TMEM55B as a critical regulator of lysosomal positioning and autophagic competence in neurons, with direct relevance to Alzheimer’s disease–associated neurodegeneration. Integrating human clinical–pathological datasets with in vivo *Drosophila* models and human iPSC-derived neurons, we demonstrate that TMEM55B expression is selectively reduced in severe AD and that its loss of function exacerbates lysosomal mislocalization, impairs autophagic flux, and promotes pathological tau accumulation.

**Figure 11:**
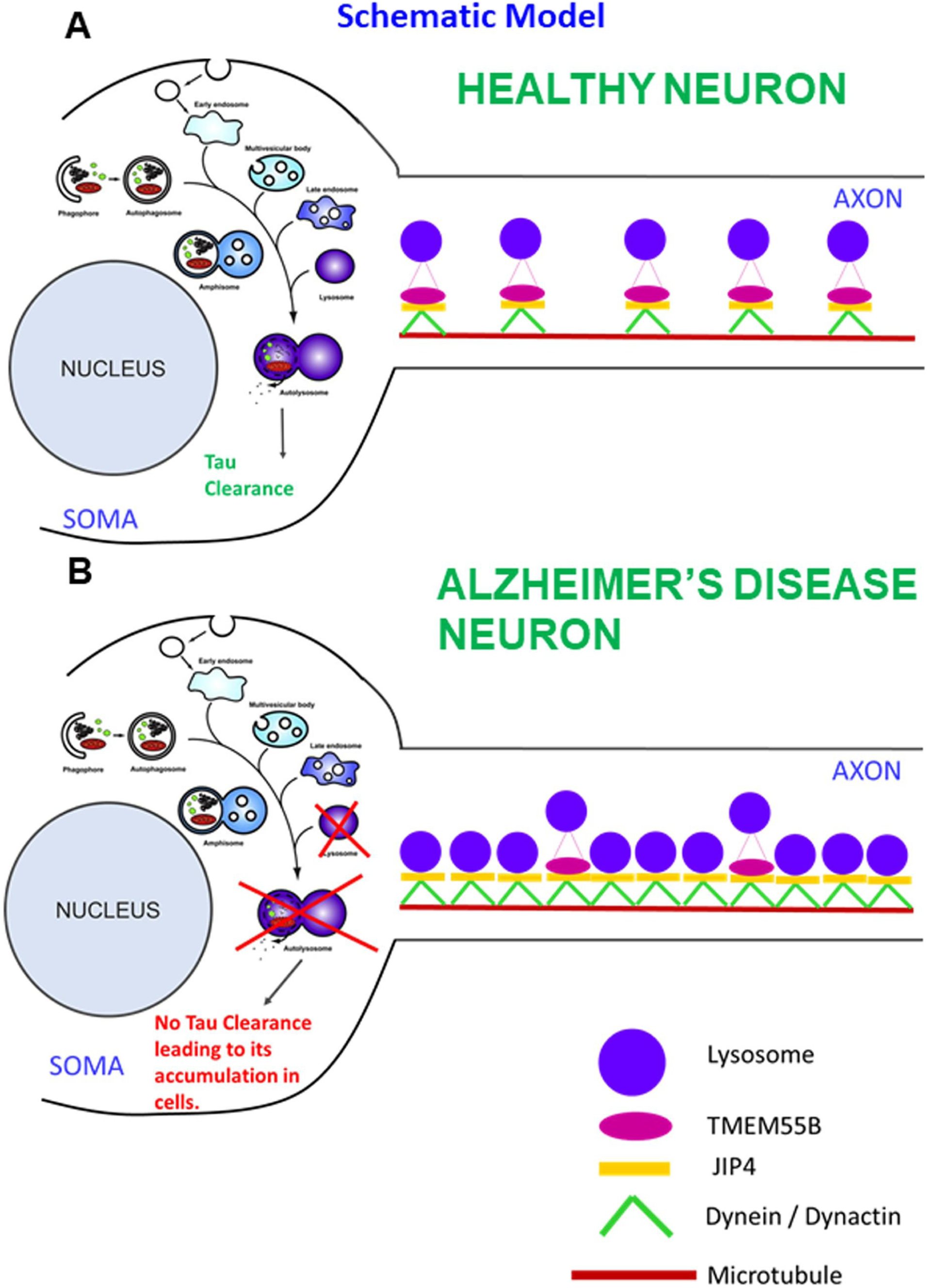
Schematic model showing the mechanism underlying TMEM55B loss of function in modulating Tau dependent neurodegeneration. (A) The neuron from healthy individual having normal levels of TMEM55B have efficient lysosomal transport from neuronal periphery towards the neuronal center. This enables the lysosomes to fuse with the amphisome [fused auto phagosome and endosome] to form autolysosome to promote autophagic clearance of Tau. (B) The neuron from AD patients having downregulated TMEM55B will have lysosomal transport defects and show lysosomal accumulation in the axons. This will prevent lysosomes from fusing with the amphisome to form autolysosomes, leading to impaired autophagic clearance of Tau.

Mechanistically, TMEM55B deficiency may impair retrograde lysosomal transport, potentially leading to reduced lysosomal clearance of Tau and increased neuronal vulnerability. These findings suggest that TMEM55B could act as a disease-severity–associated factor in neurodegeneration, indicating that lysosomal trafficking pathways may contribute to late-stage AD pathology. Overall, our results underscore the potential importance of lysosomal homeostasis in maintaining neuronal integrity and provide a rationale for further investigation of TMEM55B-dependent pathways as possible therapeutic targets in lysosome-mediated neurodegenerative disorders.

## Supporting information

"D:\NOD resubmission\NOD resubmission FINAL PACKAGE\FINAL SUBMISSION 2026\Table 1 DIOPT.zip"

## Acknowledgements

The ROSMAP analysis results published here are in whole or in part based on data obtained from the AD Knowledge Portal (https://adknowledgeportal.org). Samples for this study were provided by the Rush Alzheimer’s Disease Center, Rush University Medical Center, Chicago. Data collection was supported through funding by NIA grants P30AG10161, R01AG15819, R01AG17917, R01AG30146, R01AG36836, U01AG32984, U01AG46152, the Illinois Department of Public Health, and the Translational Genomics Research Institute.’ Additional, study data were provided by the Rush Alzheimer’s Disease Center, Rush University Medical Center, Chicago. Data collection was supported through funding by NIA grants P30AG10161 (ROS), R01AG15819 (ROSMAP; genomics and RNAseq), R01AG17917 (MAP), R01AG30146, R01AG36042 (5hC methylation, ATACseq), RC2AG036547 (H3K9Ac), R01AG36836 (RNAseq), R01AG48015 (monocyte RNAseq) RF1AG57473 (single nucleus RNAseq), U01AG32984 (genomic and whole exome sequencing), U01AG46152 (ROSMAP AMP-AD, targeted proteomics), U01AG46161(TMT proteomics), U01AG61356 (whole genome sequencing, targeted proteomics, ROSMAP AMP-AD), the Illinois Department of Public Health (ROSMAP), and the Translational Genomics Research Institute (genomic).

This work was supported by an NIH/NIA [National Institute on Aging] grant to Radek Dobrowloski and Wilma Friedman (5R01AG062475-03). We are grateful to Dr. Radek Dobrowolski for his guidance in initiating this work and for introducing us to the field of lysosome-mediated autophagic dysfunction in Alzheimer’s disease. We thank Ms. Angelid Pabon for technical assistance with initial fly and cell culture experiments. We acknowledge using stocks from Bloomington Drosophila Stock Center and Rutgers software support for access to licensed softwares. We would also like to thank lab members and colleagues for their critical review of the manuscript.

## Author contributions

Conceptualization – Radek Dobrowolski and Bidisha Roy; Data curation – Bidisha Roy, Raluca Contu, Maciej Jeziorek, Ching-On Wong; Formal analysis – Bidisha Roy, Raluca Contu, Ching-On Wong; Funding acquisition – Radek Dobrowolski, Wilma Friedman, Bidisha Roy; Investigation - Bidisha Roy, Raluca Contu, Maciej Jeziorek; Methodology - Bidisha Roy, Raluca Contu, Maciej Jeziorek; Project administration – Bidisha Roy; Supervision – Radek Dobrowolski and Wilma Friedman; Validation and Visualization – Bidisha Roy; Writing – original draft, review and editing – Bidisha Roy; Writing of iPSC derived neuronal culture protocol –Raluca Contu; Manuscript Editing – Wilma Friedman, Ching-On-Wong, Raluca Contu.

## Supplementary figures

**Supplementary Figure 1:**
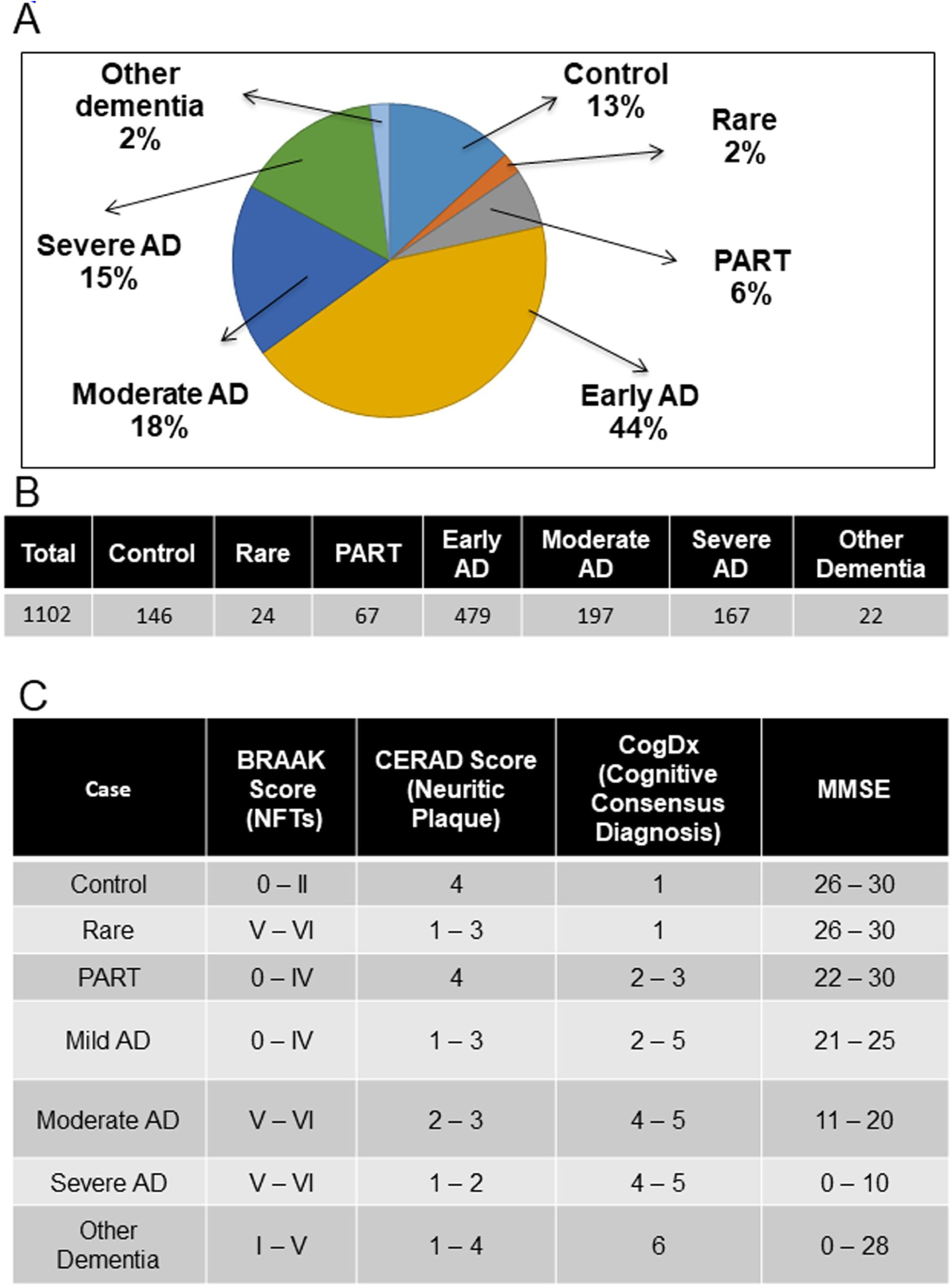
Stratification of the ROSMAP patient cohort based on clinical and pathological criteria. (A), (B) Pie chart and table representing percentage and an absolute number of patients with various forms of Alzheimer’s disease and other Tauopathies, in the Religious Order Study – Memory and Aging Project cohort study. (C) Table showing the correlation of various Tauopathies (Severe AD, Moderate AD, Early AD, Other Tauopathies like Primary Age-Related Tauopathy) and rare forms (high neuropathological hallmarks but no cognitive impairment) with respect to BRAAK and CERAD score.

**Supplementary Figure 2:**
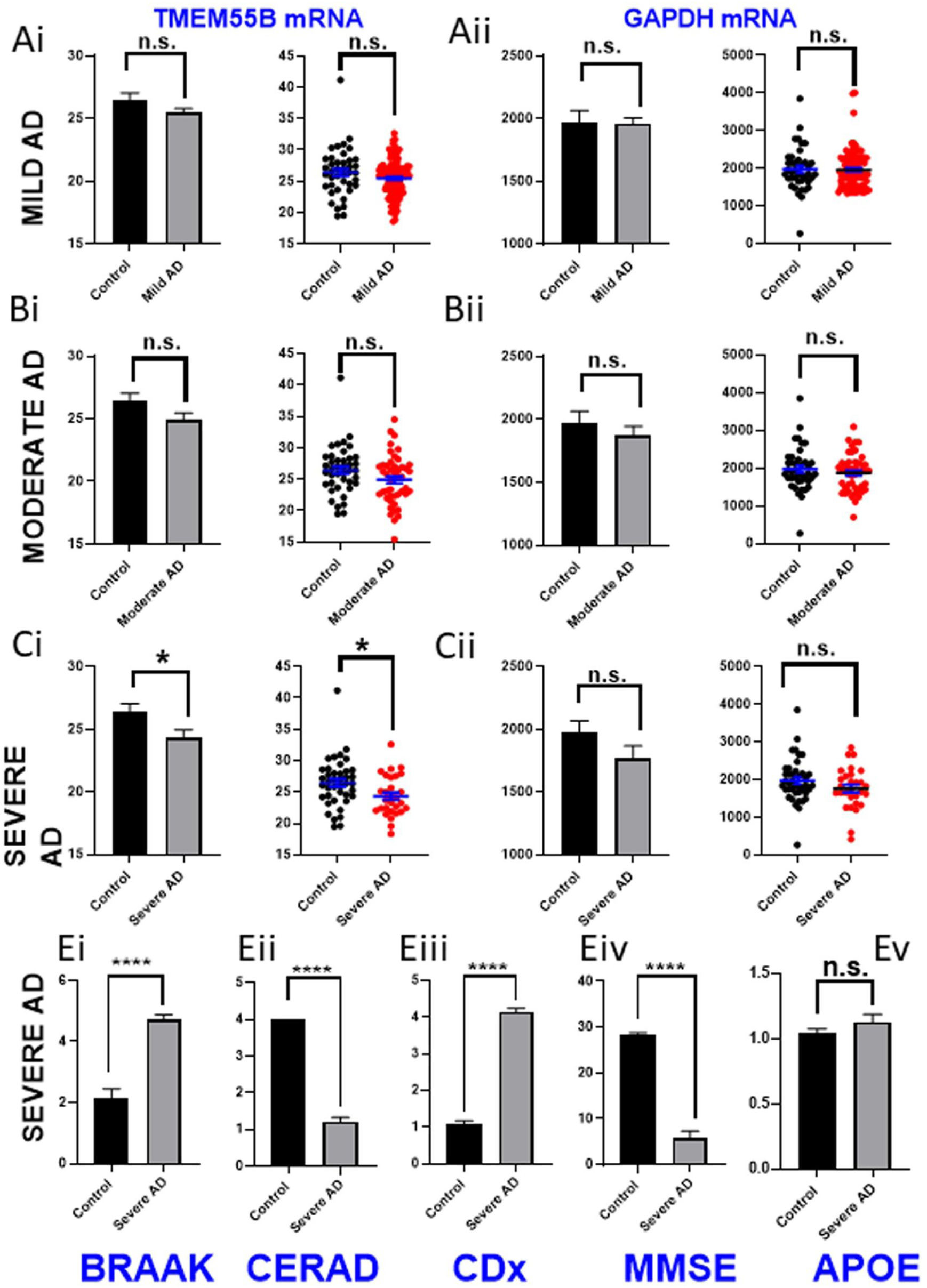
Quantitative levels of TMEM55B mRNA levels in control, mild, moderate, and severe AD patients using ROSMAP RNA Seq dataset and clinicopathological characterization of Control and Severe AD patients from the ROSMAP dataset. **(Ai-Cii)** Bar graphs and scatter plots representing quantitative mRNA levels of TMEM55B and housekeeping gene GAPDH in mild (Ai, Aii), moderate (Bi, Bii), and severe AD (Ci, Cii) cases respectively, using the RNA Seq dataset of ROSMAP studies. *: p < 0.05; **: p < 0.01; ***: p < 0.001; n.s: p ≥ 0.05. p-value was calculated using Prism 7’s two-tailed Unpaired t-tests. **(Ei-Ev)** Comparison of BRAAK stage (Ei), CERAD Score (Eii), CDx (Eiii), MMSE (Eiv), and APOE AD risk (Ev) between control and severe AD patients. Control patients: N= 41; Severe AD patients: N=29; Moderate AD patients: N= 47; Mild Ad patients: N=100. *: p < 0.05; **: p < 0.01; ***: p < 0.001. p-value was calculated using Prism 7’s two-tailed Unpaired t-tests.

**Supplementary Figure 3:**
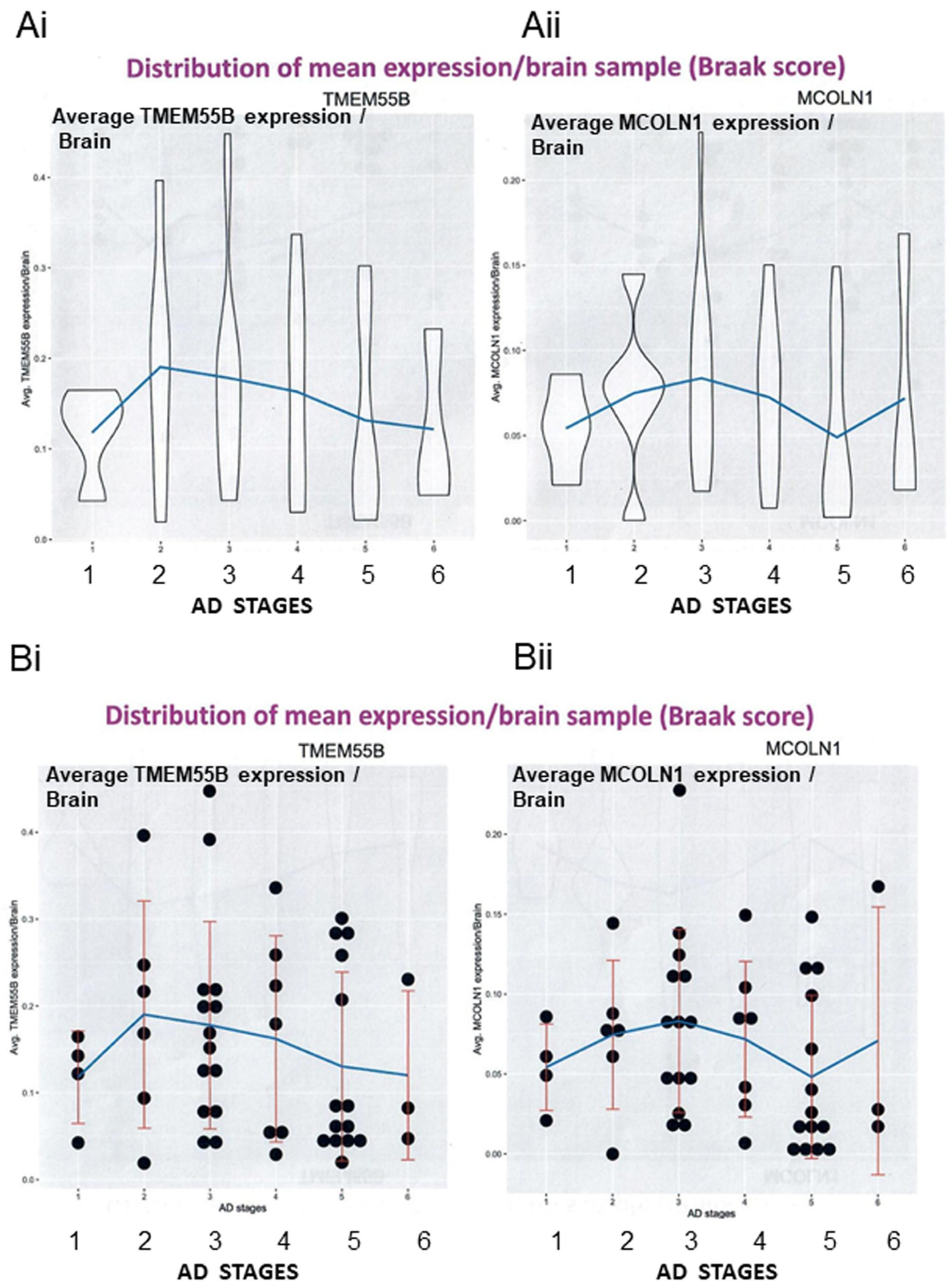
TMEM55B RNA levels are lowered in single neurons of AD patients. (Ai, Bi): Violin plot and dot plots indicating single cell RNA Seq analysis of neuron’s nuclear dataset from the ROSMAP patient cohort in correlation with the BRAAK stage of the patient samples showed initial elevated levels of TMEM55B at Braak stage 2, followed by a decline at BRAAK stage 6. (Aii, Bii): In comparison, the trendline of MCOLN1 in the violin and dot plots is different from TMEM55B (B), showing an initial increase at Braak stage 3, but without a considerable decline at Stage 6; The single nucleus analysis was done using the neurons from the gray matter of the dorsal-prefrontal cortex. N = 465 patients.

**Supplementary Figure 4:**
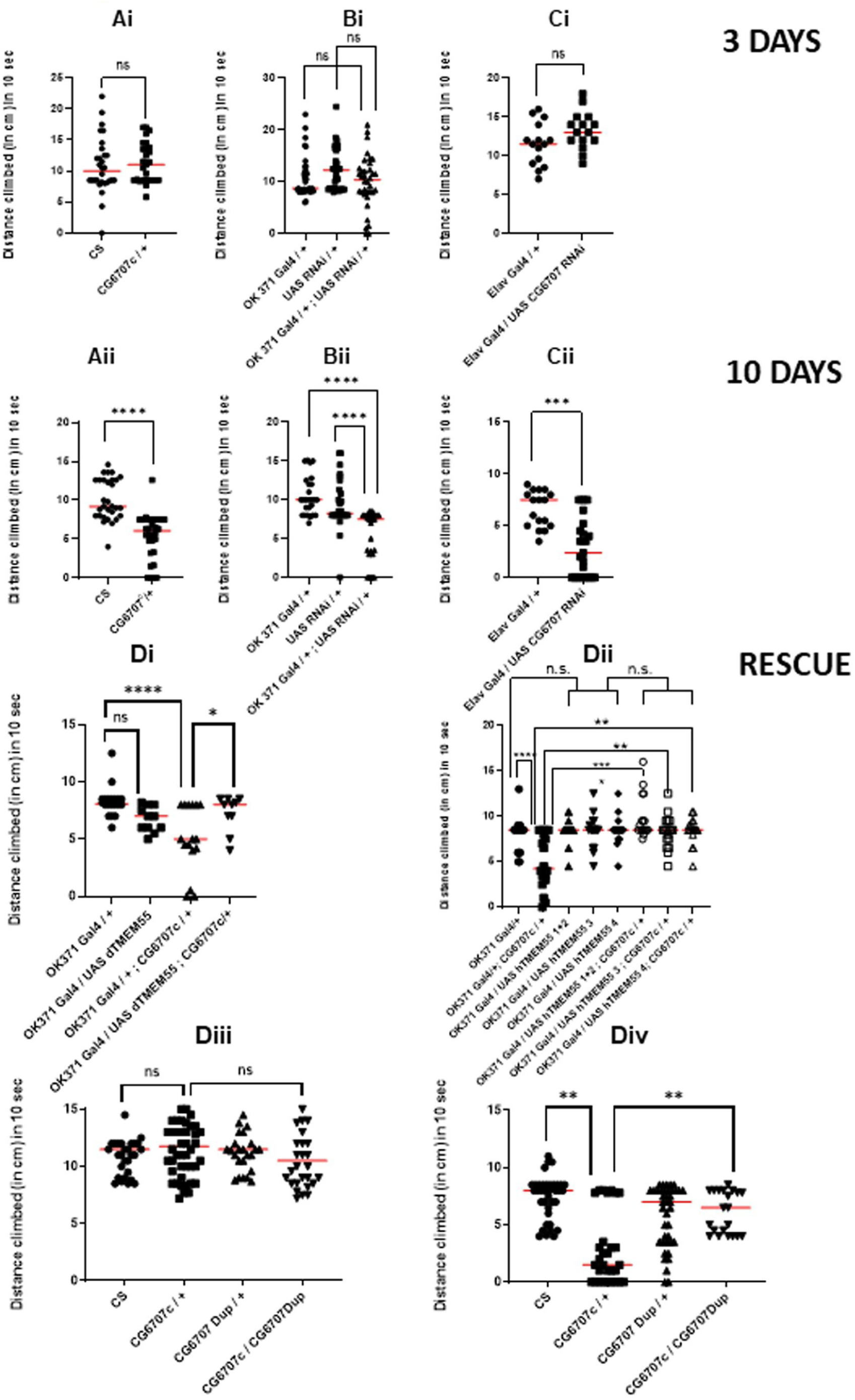
Loss of dTMEM55 results in progressive locomotor impairment in adult flies. Scatter plots show distance climbed in a 10-s negative geotaxis assay in 3-day-old and 10-day-old adult male flies. **(A)** Comparison between wild-type CS and loss of function piggyBac insertion dTMEM mutants (CG6707c or CG6707*^c01122^*). **(B)** RNAi-mediated knockdown of dTMEM in glutamatergic neurons compared with Gal4 driver-alone and UAS responder-alone controls. **(C)** Pan-neuronal RNAi knockdown of dTMEM compared with genetic controls. **(D)** Rescue experiments showing that (i) re-expression of fly dTMEM in glutamatergic neurons, (ii) expression of human TMEM55 isoforms, and (iii–iv) genomic duplication of dTMEM improve locomotor performance in CG6707*^c^* mutants. Each data point represents an individual fly; horizontal bars indicate mean ± SEM. N ≥ 16 flies per group unless stated otherwise. *: p < 0.05; **: p < 0.01; ***: p < 0.001; n.s: p ≥ 0.05. p-value was calculated using Prism 7’s two-tailed Unpaired t-test and One-way ANOVA, multiple comparisons.

**Supplementary Figure 5:**
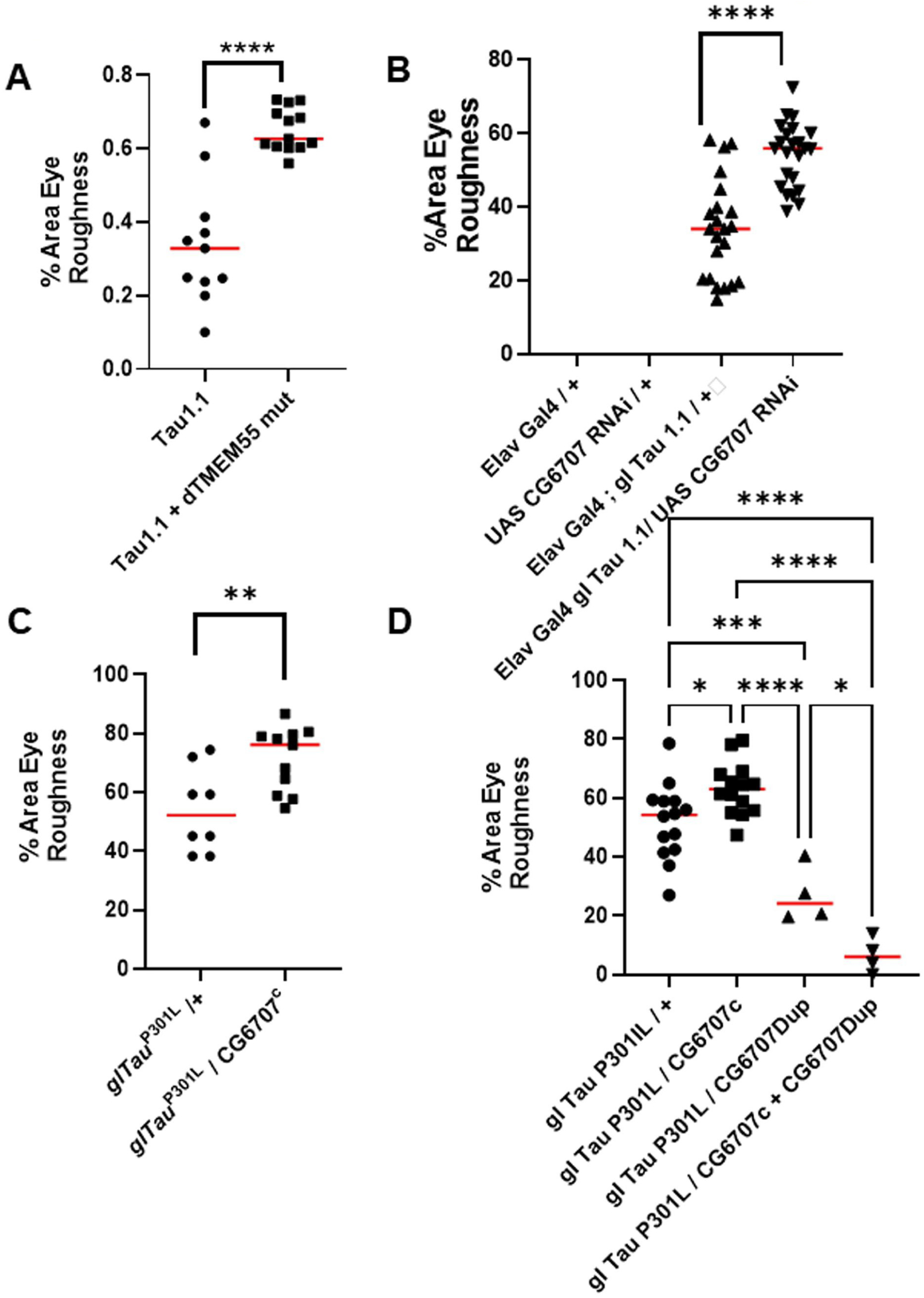
Scatter plots showing enhanced Tau-mediated retinal degeneration in transgenic flies with downregulated TMEM55. (A-D) Scatter plots depicting the severity of Tau-induced retinal degeneration across individual transgenic flies with downregulated TMEM55. Each data point represents an individual fly scored for eye roughness or retinal degeneration, enabling assessment of inter-individual variability in neurodegenerative phenotypes. The use of scatter plots, rather than simple bar graphs, allows visualization of the full distribution of degeneration severity, including phenotypic heterogeneity and potential outliers, which may be masked by mean-based representations. This approach provides a more sensitive and biologically informative evaluation of the extent to which TMEM55 downregulation exacerbates Tau-mediated retinal degeneration. [N > 5], *: p < 0.05; **: p < 0.01; ***: p < 0.001; n.s: p ≥ 0.05. p-value was calculated using Prism 7’s two-tailed Unpaired t-test and One-way ANOVA, multiple comparisons.

**Supplementary Figure 6:**
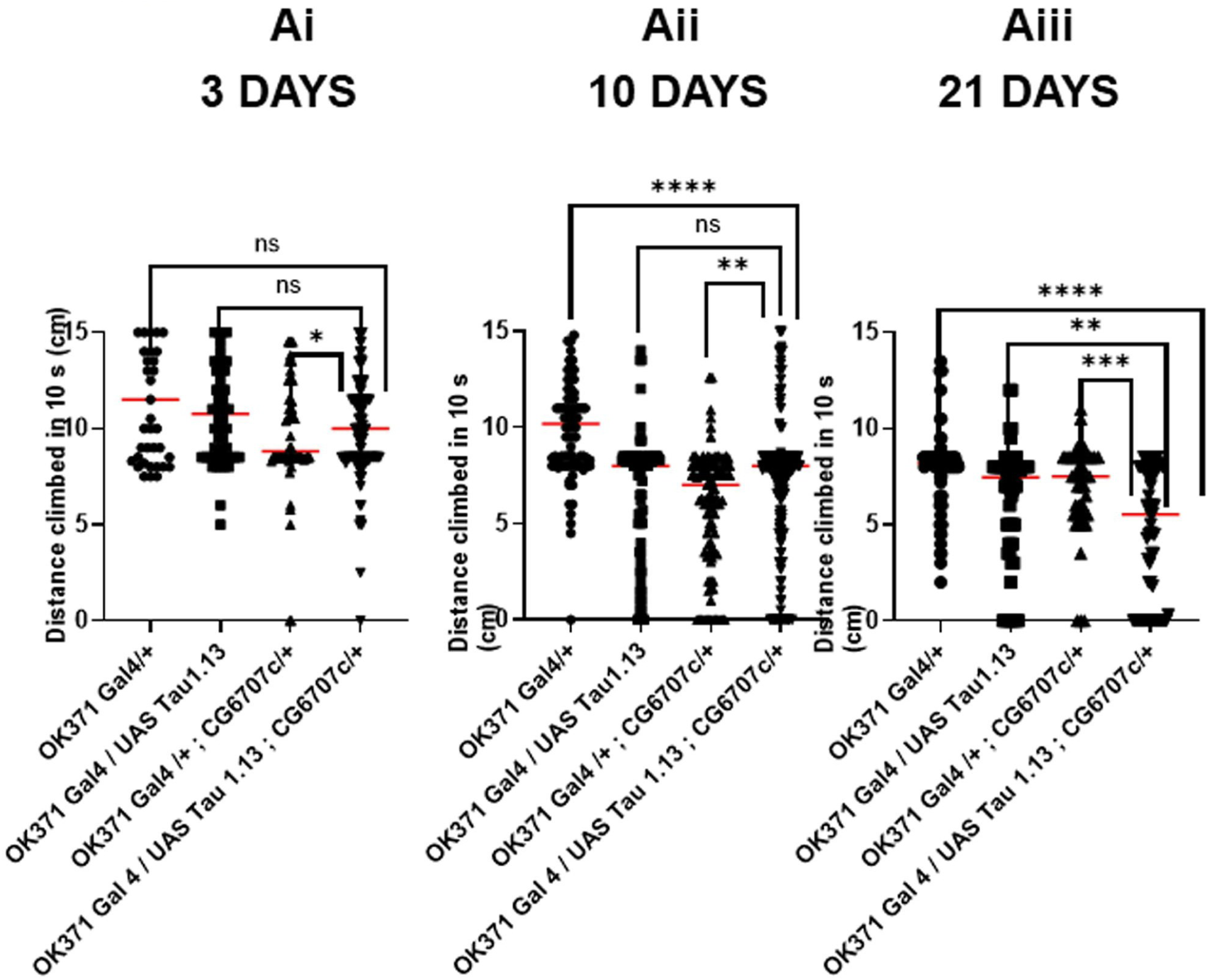
Downregulation of dTMEM exacerbates age-dependent Tau-mediated locomotory dysfunction in adult flies. Ai–Aiii) Scatter plots depicting locomotory performance of adult male flies measured as distance climbed (cm) within 10 seconds at 3 days (Ai), 10 days (Aii), and 21 days (Aiii) of age across the indicated genotypes. Each data point represents an individual fly (N = 17 males per genotype), enabling visualization of inter-individual variability in locomotor ability. Red horizontal lines denote the mean performance for each genotype. The use of scatter plots, rather than simple bar graphs, allows assessment of the full distribution of locomotory defects, including variability, age-dependent progression, and potential outliers, providing a more sensitive evaluation of how dTMEM downregulation enhances Tau-mediated locomotory dysfunction. *: p < 0.05; **: p < 0.01; ***: p < 0.001; n.s: p ≥ 0.05. p-value was calculated using Prism 7’s One-way ANOVA, multiple comparisons.

**Supplementary Figure 7:**
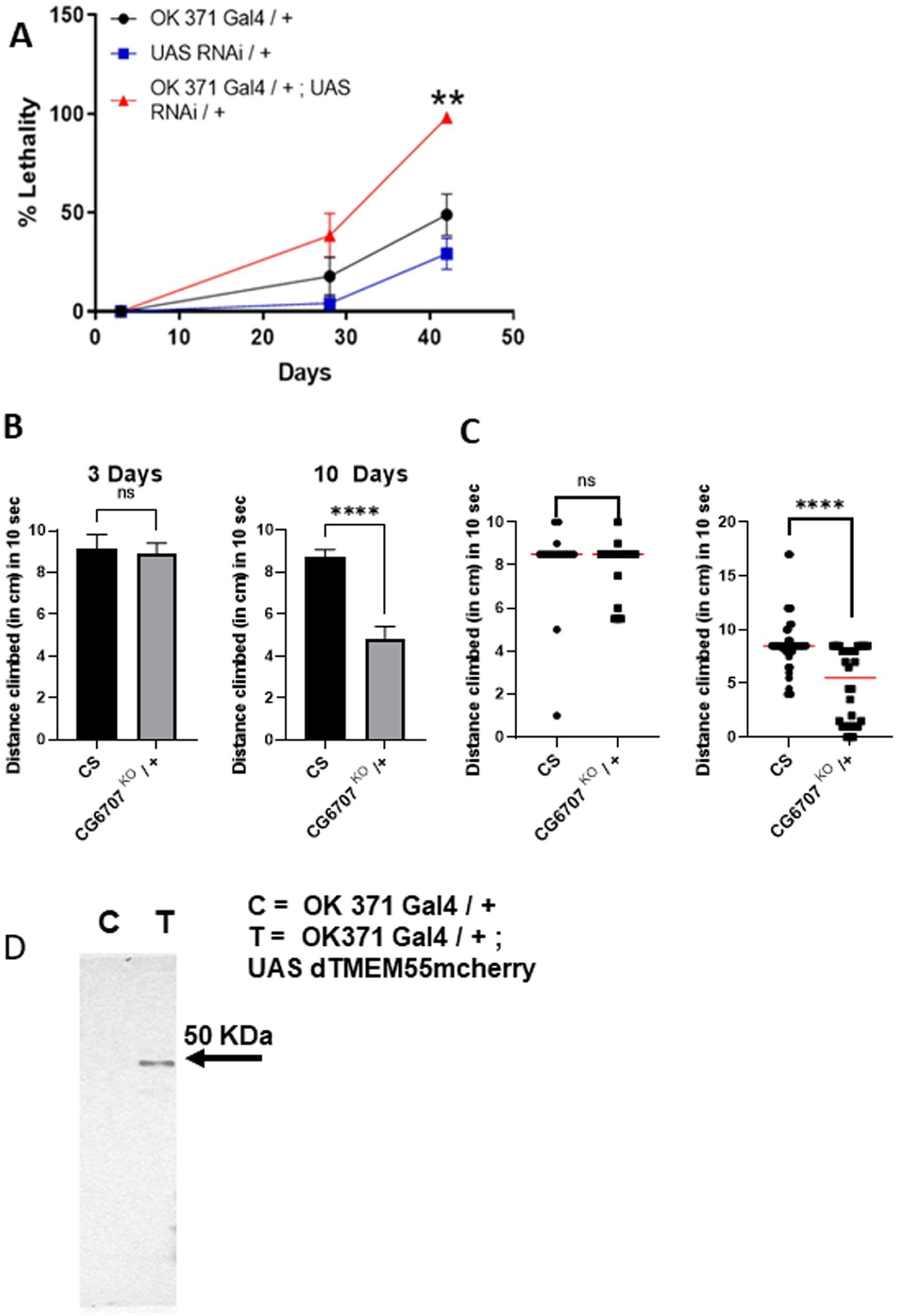
(A) Lethality assay comparing flies with RNAi-mediated dTMEM in glutamatergic neurons with respect to the Gal4 driver and UAS responder alone. N = 4 tubes [each tube = 10-12 flies]. *: p < 0.05; **: p < 0.01; ***: p < 0.001; n.s: p ≥ 0.05. p-value was calculated using Prism 7’s two-tailed Unpaired t-test and One-way ANOVA, multiple comparisons. (B) dTMEM knock-out flies generated using CRISPR technology, lead to locomotory defects in an age-dependent fashion. Distance climbed by adult male flies (3 days and 10 days old) in 10 seconds compared between wild-type CS and CRISPR technology-generated dTMEM knockout flies (CG6707 ^KO^). N > 20 flies. (C) CRISPR-mediated knockout of dTMEM results in age-dependent locomotor impairment. Scatter plot showing the distance climbed in 10 s by individual adult male flies at 3 and 10 days of age, comparing wild-type CS controls and CRISPR-generated dTMEM knockout flies (CG6707 ^KO^). Each point represents a single fly; N > 20 flies per genotype and age group. (D) Immunoblots showing the 50KDa dTMEM + mCherry band, in comparison to the control line lacking any expression of dTMEM (OK371 Gal4 driver alone). *: p < 0.05; **: p < 0.01; ***: p < 0.001; n.s: p ≥ 0.05. p-value was calculated using Prism 7’s two tailed Unpaired t-test.

**Supplementary Figure 8:**
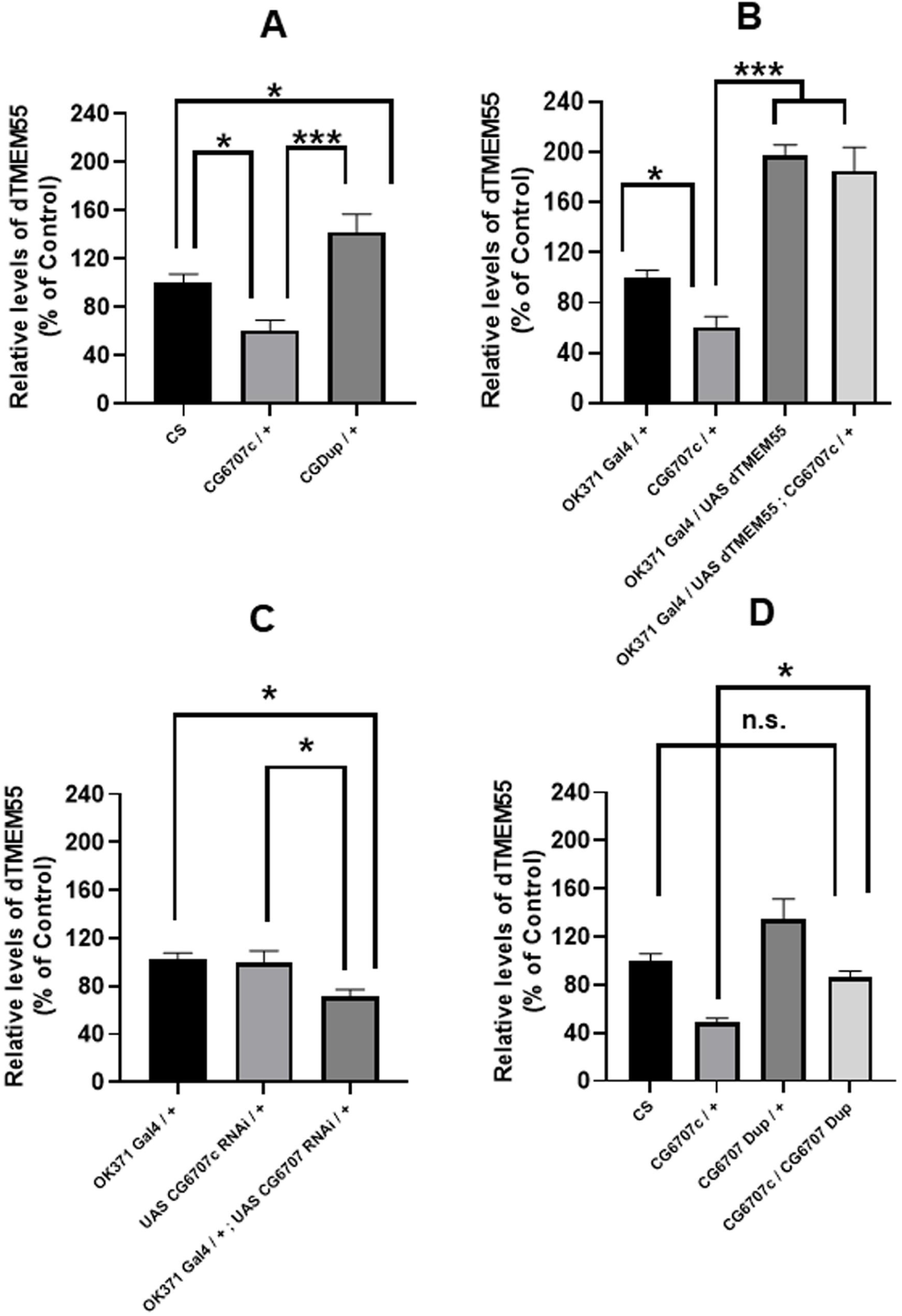
RT PCR to determine dTMEM levels in the various genetic transgenic animals. (A) dTMEM mRNA transcripts are lower and higher than the wild type in the CG6707c / + (dTMEM mutant) and CGDup / + (dTMEM Duplication) lines. (B) dTMEM mRNA levels are restored in CG6707c / CGDup transgenic combination. (C) Downregulation of dTMEM using the UAS RNAi line leads to decrease levels than the Gal4 driver or UAS responder alone. (D) Overexpression of dTMEM using the UAS dTMEM mcherry line leads to an increase in the levels of endogenous dTMEM. Genotypes: CS: Wild type; CG6707c / +: dTMEM heterozygous mutant; CGDup / +: heterozygous dTMEM duplication; CG6707c / CGDup: double transgenic with dTMEM mutation and dTMEM duplication; OK371 Gal 4/ +: Gal4 driver; UAS CG6707 RNAi / +: UAS responder; OK371 Gal4 / UAS CG6707 RNAi: dTMEM knockdown in glutamatergic neurons; UAS CG6707 mCherry: UAS responder overexpressing dTMEM; OK371 Gal4 / UAS CG6707 mCherry: dTMEM overexpression in glutamatergic neurons. N = 5 sets. *: p < 0.05; **: p < 0.01; ***: p < 0.001; n.s: p ≥ 0.05. p-value was calculated using Prism 7’s two-tailed Unpaired t-test and One-way ANOVA, multiple comparisons.

**Supplementary Figure 9:**
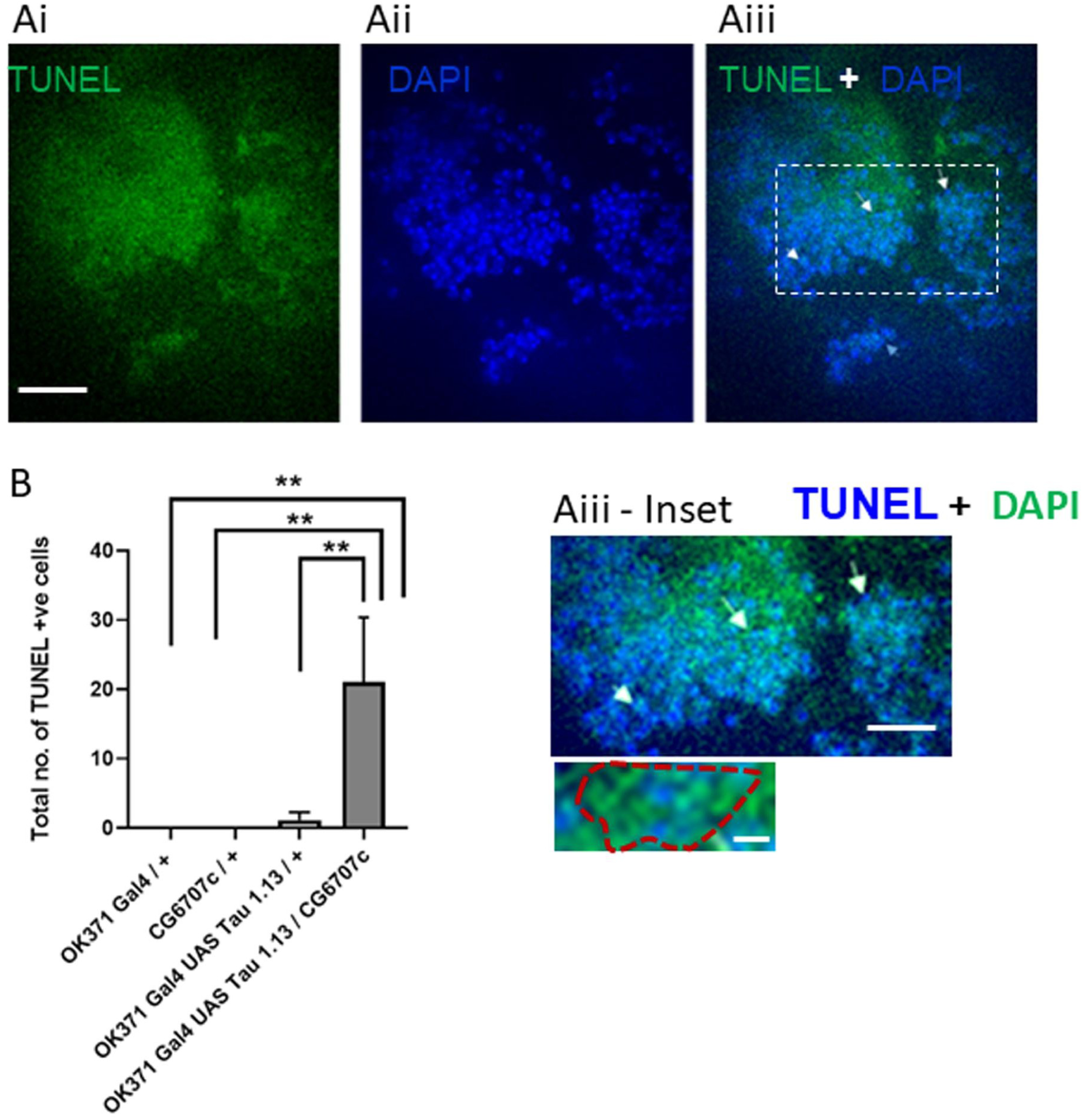
Downregulation of TMEM55 enhances Tau-mediated cell death. (A) Immunofluorescence image of TUNEL positive cells in the optic lobe region per brain hemisphere of dTMEM flies overexpressing Tau. (B) Quantitative bar graph representing the total number of Tunel positive cells in the optic lobe region per hemisphere, in a single fly brain in various genotypes listed below. Genotypes: This includes the control (OK371 Gal4 / +); Tau over-expressing (OK371 Gal4 / UAS Tau 1.13); dTMEM55 mutant (OK371 Gal4 / +; CG6707c / +) and dTMEM55 mutants overexpressing Tau (OK371 Gal4 / +; CG6707c / UAS Tau 1.13) flies. N > 40 flies for A; N = 8 tubes [each tube = 10-12 flies] for B; N = 6 brains for *: p < 0.05; **: p < 0.01; ***: p < 0.001; n.s: p ≥ 0.05. All flies analyzed were 20 –21 days old. The P-value was calculated using Prism 7’s two-tailed Unpaired t-test and One-way ANOVA, multiple comparisons. Scale Bar: 30 µm for Ai, Aii, Aiii; 16 µm and 4 µm for Aiii, Inset 1 and 2 respectively.

**Supplementary Figure 10:**
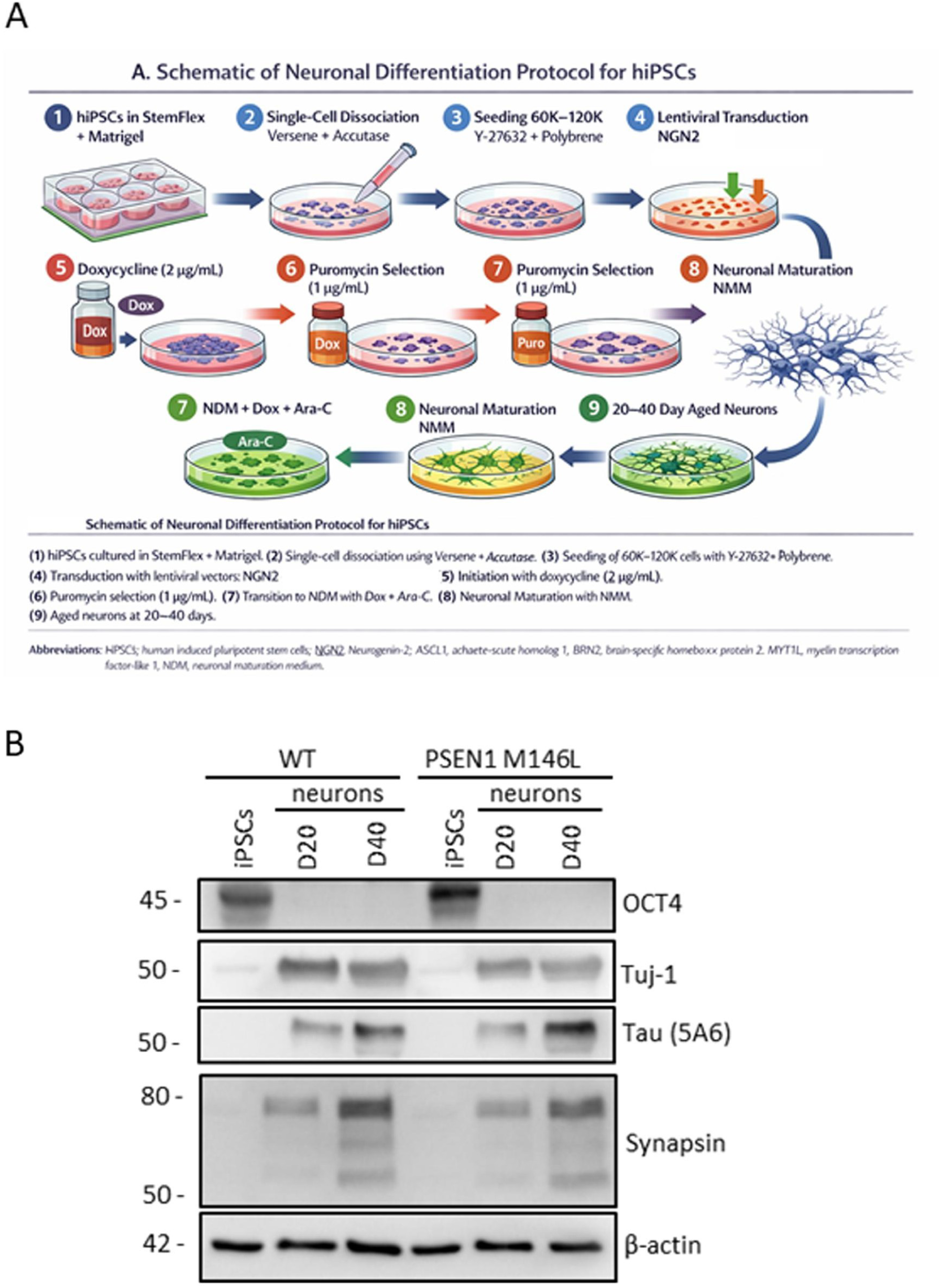
(A) Schematic overview of the hiPSC-to-neuron differentiation and maturation protocol. Human induced pluripotent stem cells (hiPSCs) maintained under feeder-free conditions in StemFlex medium on Matrigel-coated surfaces were dissociated into single cells and seeded onto poly-D-lysine/Matrigel–coated culture ware in the presence of the ROCK inhibitor Y-27632 and polybrene. Cells were transduced with lentiviral vectors expressing neurogenin-2 (NGN2). Neuronal fate commitment was induced by doxycycline-mediated activation of NGN2 expression, followed by puromycin selection to enrich for successfully transduced cells. Cultures were sequentially transitioned from neuronal induction medium (NIM) to neuronal differentiation medium (NDM), with transient cytosine β-D-arabinofuranoside (Ara-C) treatment applied where necessary to eliminate residual proliferating non-neuronal cells. From day 7 onward, neurons were maintained in neuronal maturation medium (NMM) with regular half-medium changes to support neuronal survival and functional maturation. Neurons were aged for 20 or 40 days in vitro prior to downstream biochemical and immunocytochemical analyses. **Abbreviations:** hiPSC, human induced pluripotent stem cell; NGN2, neurogenin-2; NIM, neuronal induction medium; NDM, neuronal differentiation medium; NMM, neuronal maturation medium; Ara-C, cytosine β-D-arabinofuranoside; Dox, doxycycline. (B) **Characterization of iPSC derived control and AD neurons.** (A) Immunoblot image derived from of 20 days and 40 days old, iPSC derived control and AD forebrain neuronal extracts, processed using antibodies against stem cell marker, OCT4; differentiated neuronal marker like Tuj1 and Tau; synaptic marker like Synapsin.

**Supplementary Figure 11:**
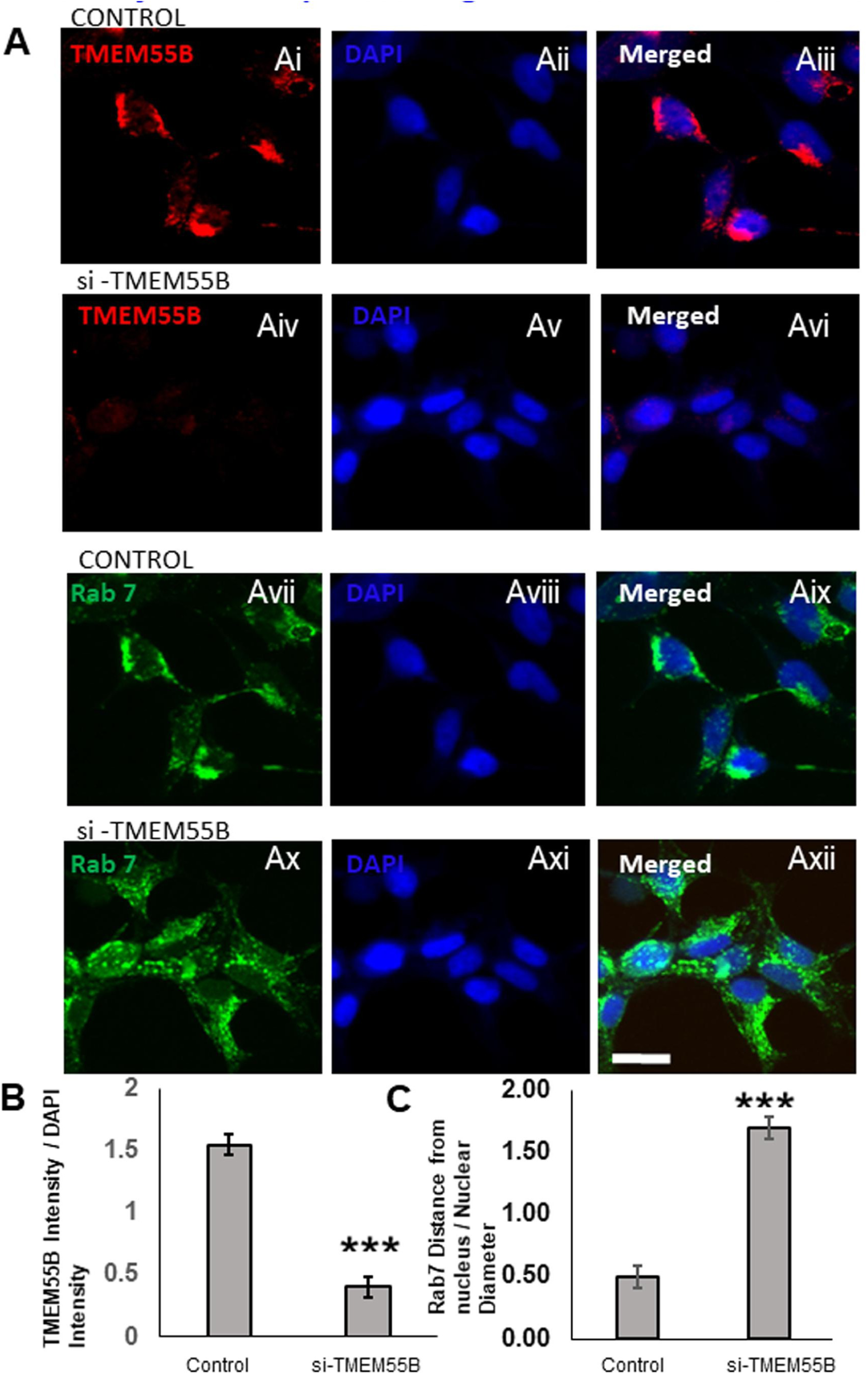
Downregulation of TMEM55B leads to lysosomal positioning defects in HEK293T cells. (Ai-Axii, B, C): Quantitative immunofluorescence of control and si-RNA mediated TMEM55B knock down HEK293T cell stained for TMEM55B, RAB7 and DAPI. (Ai-Avi, B): TMEM55B levels were lowered in the HEK293T cells’ soma in the si-RNA mediated TMEM55b knock down cells significantly in comparison to the control HEK293T cells. N= 46 cells. (Ax-Axii, C) Rab7 staining spanned a greater length in the soma and covered more distance from the nucleus, covering both the perinuclear and peripheral regions of the HEK293T cells treated with si-TMEM55B RNA. On the other hand, Rab7 staining remained restricted to the perinuclear space and spanned areas closer to the nucleus in the control HEK293T cells (Avii-Aix, B). N= 54 cells.*: p < 0.05; **: p < 0.01; ***: p < 0.001; n.s: p ≥ 0.05. p value was calculated using Prism 7’s two tailed Unpaired t-test and One-way Anova, multiple comparison. Scale Bar: 20 µm

**Supplementary Figure 12:**
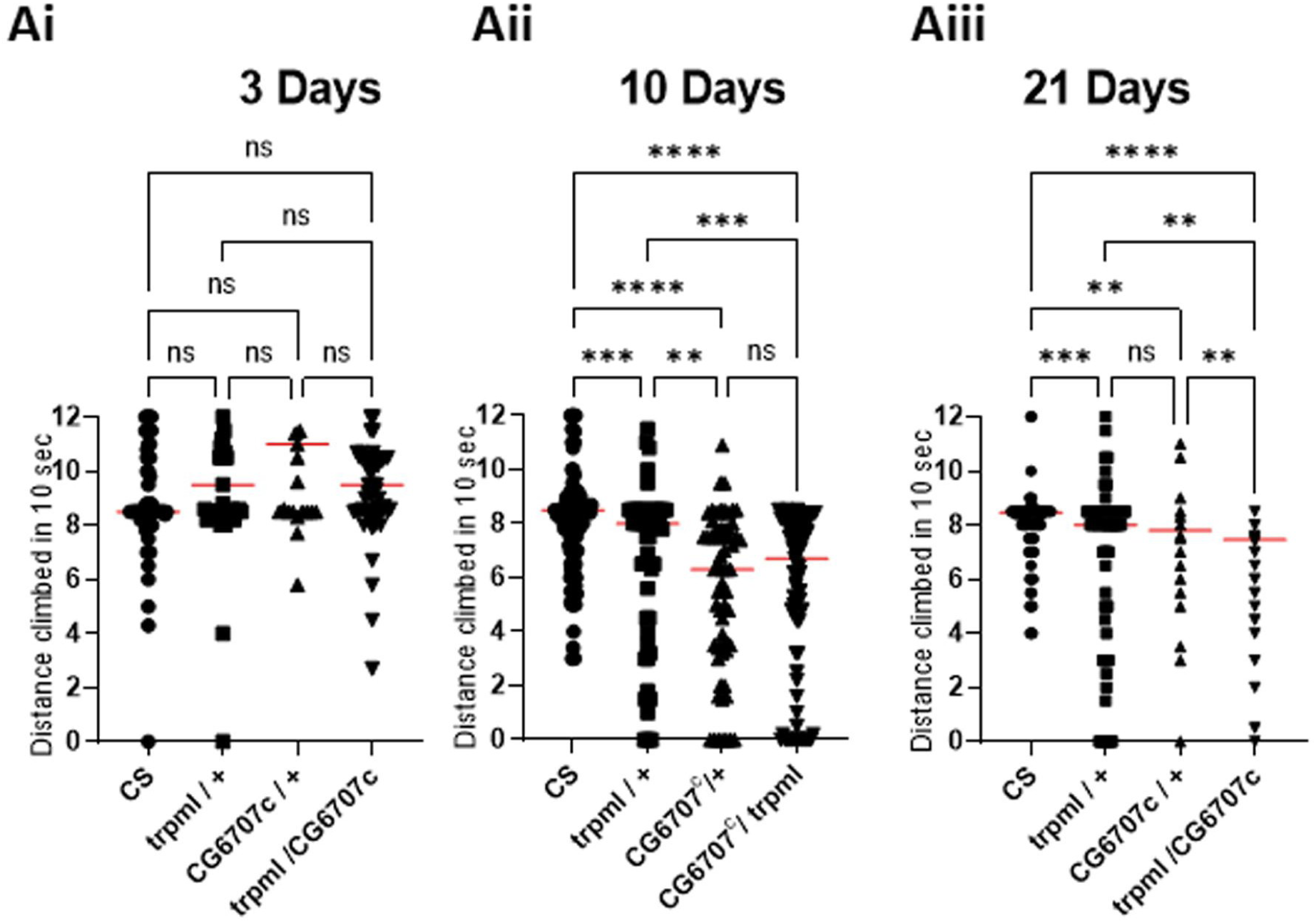
Concomitant loss of dTMEM and trpml exacerbates age-dependent locomotory defects in *Drosophila.* Scatter plots showing locomotory performance of adult male flies measured as the distance climbed in 10 seconds at 3 days (Ai), 10 days (Aii), and 21 days (Aiii) of age across the indicated genotypes. Each data point represents an individual fly (N > 60 per genotype). Scatter plots were used instead of bar graphs to display individual measurements and biological variability, allowing visualization of data distribution and potential outliers that may be masked by summary statistics alone, particularly in age-dependent behavioral assays. Red horizontal bars indicate mean values. Statistical significance is indicated as shown, p < 0.05; **: p < 0.01; ***: p < 0.001; n.s: p ≥ 0.05. p-value was calculated using Prism 7’s two tailed Unpaired t-test and One-way Anova, multiple comparison.

**Supplementary Figure 13:**
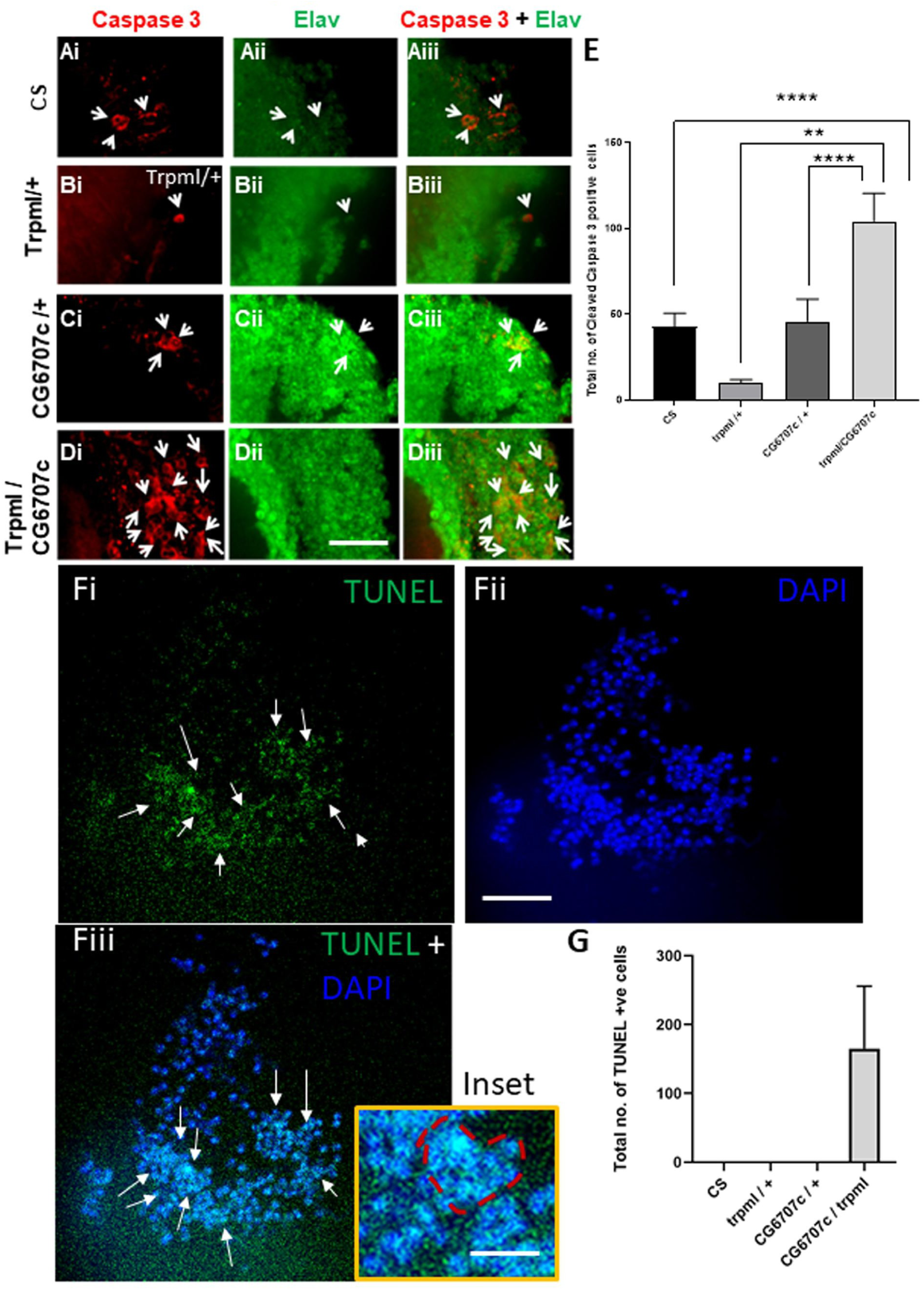
Loss of dTMEM and trpml leads to increased accumulation of nuclear cleaved caspase 3 and increase in the number of TUNEL positive cells. (Ai – Diii) Immunohistochemically staining of optic lobe of *Drosophila* brain with anti-Cleaved Caspase 3 (red) and anti-Elav (green) antibodies. (E) Bar graphs representing the total number of cleaved caspase 3 positive cells in the optic lobe region per brain hemisphere of flies with varied genotypes, N = 5 brains. (Fi-Fiii) TUNEL positive cells in the optic lobe of trpml / CG6707c double mutant animals along with quantitative bargraph (G), N = 5 brains. Genotypes: CS: wild type CantonS; trpml / +: trpml heterozygous mutant; CG6707c / + = heterozygous dTMEM mutant; trpml / CG6707c = trpml and dTMEM double mutants. p < 0.05; **: p < 0.01; ***: p < 0.001; n.s: p ≥ 0.05. p-value was calculated using Prism 7’s two tailed Unpaired t-test and One-way Anova, multiple comparison. Scale Bar: 24 µm for Ai-Diii; 21 µm for Fi-Fiii; 9 µm for Inset.

